# Comparative population genomics unveils congruent secondary suture zone in Southwest Pacific Hydrothermal Vents

**DOI:** 10.1101/2024.05.31.596862

**Authors:** Adrien Tran Lu Y, Stéphanie Ruault, Claire Daguin-Thiébaut, Anne-Sophie Le Port, Marion Ballenghien, Jade Castel, Pierre-Alexandre Gagnaire, Nicolas Bierne, Sophie Arnaud-Haond, Camille Poitrimol, Eric Thiébaut, François H Lallier, Thomas Broquet, Didier Jollivet, François Bonhomme, Stéphane Hourdez

## Abstract

How the interplay of biotic and abiotic factors shapes current genetic diversity at the community level remains an open question, particularly in the deep sea. Comparative phylogeography of multiple species can reveal the influence of past climatic events, geographic barriers, and species life history traits on spatial patterns of genetic structure across lineages.

To shed light on the factors that shape community-level genetic variation and to improve our understanding of deep-sea biogeographic patterns, we conducted a comparative population genomics study on seven hydrothermal vent species co-distributed in the Back-Arc Basins (BABs) of the Southwest Pacific region. Using ddRAD-seq, we compared the range-wide distribution of genomic diversity across species and discovered a shared phylogeographic break. Demogenetic inference revealed shared histories of lineage divergence and a secondary contact. Low levels of asymmetric gene flow probably occurred in most species between the Woodlark and North Fiji basins, but the exact location of contact zones varied from species to species. For two species, we found individuals from the two lineages co-occurring in sympatry in Woodlark Basin. Although species exhibit congruent patterns of spatial structure (Eastern vs Western sites), they also show variation in the degree of divergence among lineages across the suture zone. Our results also show heterogeneous gene flow across the genome, indicating possible partial reproductive isolation between lineages and early speciation.

Our comparative study highlights the pivotal role of historical and contemporary factors, underscoring the need for a comprehensive approach—especially in addressing knowledge gaps on the life history traits of deep-sea species.

## Introduction

Hydrothermal vents are one of the most emblematic chemosynthesis-based ecosystems that host a highly specialized fauna. This vent fauna depends on local hydrothermal activity and is likely to share historical patterns of colonization linked to the tectonic history of the ridge system (Plouviez et al. 2009; Matabos et al. 2011; Matabos and Jollivet 2019). In contrast to other deep-sea ecosystems, vents represent a linear but highly fragmented and relatively unstable ecosystem based on chemosynthetic primary producers, which cannot live elsewhere. Hydrothermal activity is linked to specific geological features associated with the volcanic and tectonic activities of ocean ridges or submarine volcanoes (Hourdez and Jollivet 2020). Plate tectonics has previously been cited as a driver of most of the biogeographic distribution of the vent fauna (Tunnicliffe 1992), that may lead to allopatric speciation and possible secondary contacts (Hurtado et al. 2004; Johnson et al. 2006; Faure et al. 2009; Plouviez et al. 2009; Matabos et al. 2011; Johnson et al. 2013).

In contrast to the linear setting of mid-oceanic ridges such as the Mid-Atlantic Ridge (MAR) or the East Pacific Rise (EPR), the fauna of the vents of the Southwest Pacific is distributed across several geological Back-Arc-Basins (BABs) separated by abyssal plains, ridges and volcanic arcs, forming a fragmented, discontinuous complex (Figure 1). These BAB formations are estimated to be between 12 to 1 million years (My) old (Schellart et al. 2006) . An earlier study on a few species of gastropods highlighted contrasting phylogeographic patterns, including some closely related species, suggesting alternative dispersal strategies and evolutionary history to cope with fragmentation in response to a common geological history of the vent habitat in this region (Poitrimol et al. 2022). This situation raises questions regarding the drivers of the spatial distribution of genetic diversity of BABs hydrothermal fauna.

**Figure 1:**
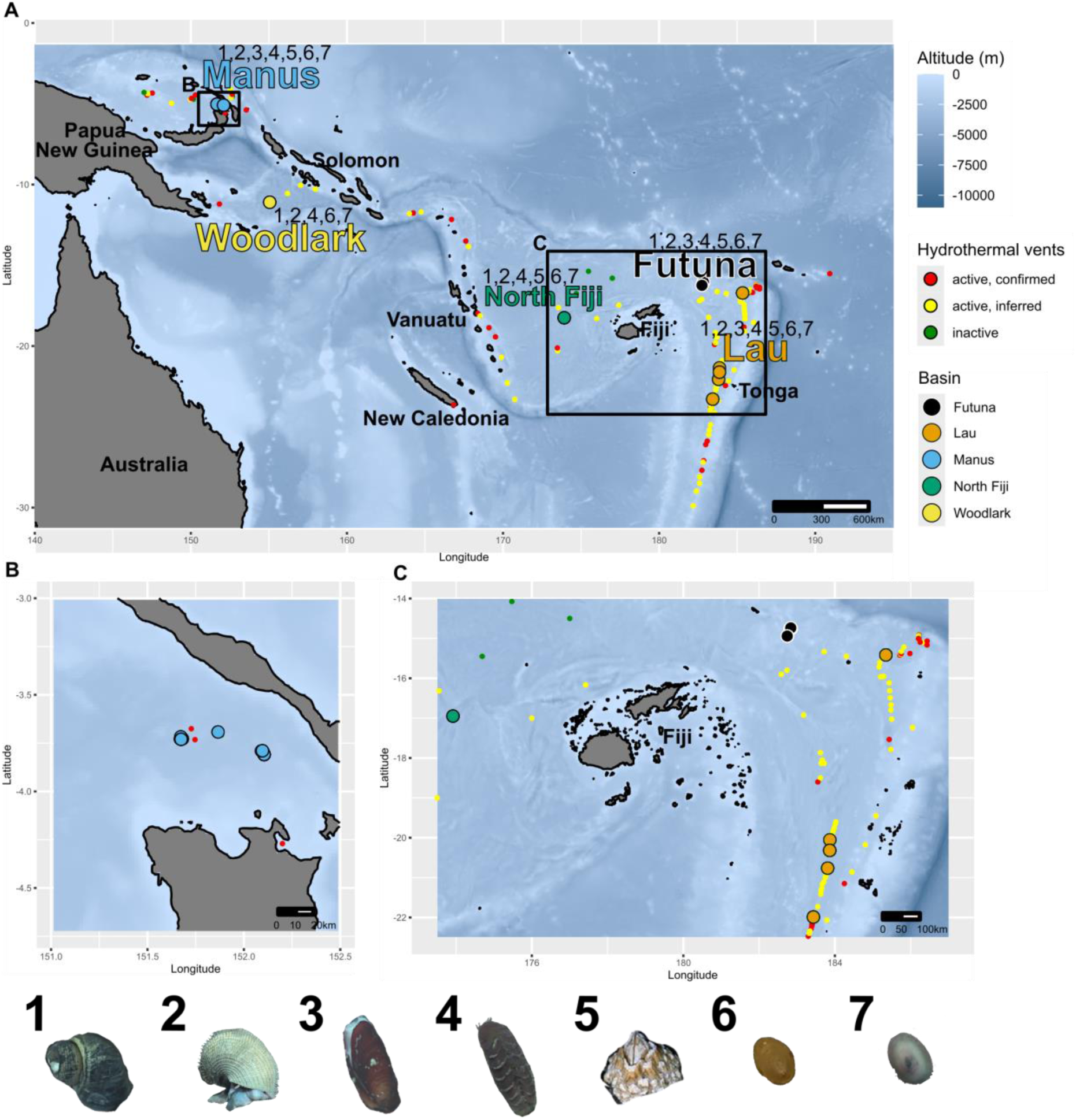
(A) Sampling areas in the South-west Pacific Ocean. Colors represent the different BABs. (B) Sampling areas for Manus BAB and (C) for North Fiji, Futuna and Lau BABs. Numbers on the panel (A) indicate the species sampled in each BAB. 1: I.nautilei, 2: A. kojimai, 3: B. manusensis 4: B. segonzaci, 5: E. ohtai, 6: S. tollmanni and 7: L. schrolli (& L. aff. schrolli). Small points represent hydrothermal vents. Red, active and confirmed. Yellow, Active and inferred. Green, Inactive. Vents activity data taken from InterRidge Vents Database V3.4.

The vent communities inhabiting these Southwest Pacific BABs appear as a single biogeographic unit (Bachraty et al. 2009; Moalic et al. 2012; Tunnicliffe et al. 2024). In contrast to other hydrothermal communities, which are mainly composed of tubeworms, mussels and shrimps, this fauna consists mainly of large symbiotic Provannidae gastropods, such as *Ifremeria nautilei*, *Alviniconcha spp*., and deep-sea *Bathymodiolus* mussels. These large engineer species create specific habitats for a wide assemblage of invertebrate species, including annelids from different families (e.g. *Polynoidae*, *Alvinellidae*, *Siboglinidae*), limpets (*Lepetodrilus spp*., *Shinkailepas spp*.), barnacles, holothurians, and crustaceans (copepods, amphipods, shrimps, and crabs) (Desbruyères et al. 2006).

While these vents support thriving oases of life, they also produce metal sulfide deposits, which attract the interest of deep-sea mining companies. The future management of the Southwest Pacific vent fauna will rely on understanding population delimitation and connectivity (gene flow and dispersal), which are of crucial importance in conservation biology (Van Dover 2011; Gena 2013; Van Dover et al. 2017; Niner et al. 2018; Washburn et al. 2019). Anthropogenic exploitation of vent resources has already begun on a Japanese site in the Northwest Pacific and some prospects have been set up in the Manus BAB (Solwara prospects), while the potential consequences of these activities are not yet understood (Carver et al. 2020).

Connectivity and renewal of vent populations is mostly driven by larval dispersal due to the sedentary nature and the strict relationship of the communities with the vent fluid. Some vagile fauna, such as fish, crabs, or shrimp, may contribute to connectivity through adult migration in response to local environmental changes, but only across very limited spatial scale (vent fields) (Lutz et al., 1994; Shank et al., 1998). Direct connectivity assessment is not technically feasible for minute larvae numbering in millions (Levin 1990; Vrijenhoek 2010). As a consequence, demographic connectivity needs to be assessed by indirect methods such as population genetics, larval dispersal modeling or recent method of elemental fingerprints tracking (Mouchi et al. 2024).

Dispersal modeling in the region suggested possible but limited larval exchange between distant BABs (Mitarai et al. 2016). However, in the context of the unstable and fragmented habitat of deep-sea hydrothermal vents, metapopulation theory predicts long-distance dispersal could mitigate risks of inbreeding and local extinction (Hamilton and May 1977; McPeek and Holt 1992). Initial genetic analyses of several vent species along mid-oceanic ridges however revealed conflicting evidence on dispersal capabilities, with some suggesting almost panmictic populations at the ridge scale while others hint at patterns of isolation by distance and stepwise (re)colonization (Audzijonyte and Vrijenhoek 2010; Teixeira et al. 2012).

Studying and disentangling the origin and maintenance of species genetic diversities is the main objective of phylogeography (Avise et al. 1987; Avise 2000; Avise 2009). Expanding this approach, to a multispecies comparative dataset within a given biome or ecosystem can highlight the effect of several factors shaping the genetic diversity (Hickerson et al. 2010; Papadopoulou and Knowles 2016). Using new methods and approaches for large genomic datasets, we can start to disentangle past and present connectivity patterns and offer a unique opportunity to describe species distribution patterns at the community level and to improve scientific guidelines for conservation (Gagnaire 2020; De Jode et al. 2023).

Our study investigated genetic diversity and connectivity patterns across a biogeographic hydrothermal system in the Southwest Pacific, using comparative population genomics analysis on seven key vent species that form the region’s primary assemblage. These species, representing different taxonomic groups but sharing similar environments and ranges across the Southwestern Pacific, were examined for phylogeographic patterns through genome-wide analysis, while exhibiting different life-history traits (lecithotrophic vs. planktotrophic larvae). It revealed a clear phylogeographic break encompassing all seven species around the Solomon-Vanuatu archipelago islands, with an additional contact zone on the Woodlark Ridge in two species. Based on inferred demogenetic histories, we propose a scenario of vicariance in which the dispersal capacities of certain species may have modulated the recontact of previously isolated faunal units. Our findings on genetic diversity and gene flow in these communities highlight the need to understand population connectivity across different geographical regions.

## Material and methods

### Sampling

Seven hydrothermal-vent species from four main vent habitats have been sampled over five West Pacific regions (Manus, Woodlark, North Fiji, Lau back-arc basins) and the Futuna volcanic arc (see SI Table 1 and Figure 1 A, B, C). These species include the emblematic symbiotic snails *Ifremeria nautilei* and *Alviniconcha kojimai*, the vent mussel *Bathymodiolus manusensis*the limpets *Shinkailepas tollmanni*, *Lepetodrilus schrolli*, and *L.* aff. *schrolli* which live on the shells of *I. nautilei* and the vent mussels, the barnacle *Eochionelasmus ohtai*, and, finally, the large scaleworm *Branchinotogluma segonzaci.* All taxa analyzed presumably represent monotypic species with the exception of *Lepetodrilus* limpets. For these, a taxonomic separation has been recently proposed by Chen & Sigwart (2023) by their genetic differences (Plouviez et al. 2019) with the sympatric co-occurrence of their mitochondrial haplotypes in Woodlark (Poitrimol et al. 2022) without knowing whether the speciation process was already achieved. As these entities are likely to hybridize and clearly fall into the grey zone of speciation, we chose to treat them as one complex unit (*L. schrolli & L.* aff*. schrolli*) to model their past demographic history in a single framework. Altogether, we chose these seven taxa as they occupy nearly the same habitat in at least three distinct basins on both the western and eastern sides of this Pacific region and display very different dispersal-related life histories (with a larval development assumed to be lecithotrophic for *I. nautilei*, *B. segonzaci*, *E. ohtai* whereas it is planktotrophic for *A. kojimai*, *B. manusensis* and and *S. tollmanni,* see discussion).

Animal collections were made during the Chubacarc cruise in 2019 (chief scientists S. Hourdez & D. Jollivet) on board the RV *L’Atalante* with the ROV Victor 6000 (Hourdez and Jollivet 2019). All species, except for *B. segonzaci* (collected on vent chimney walls), were sampled from diffuse venting areas with the tele-manipulated arm of the ROV and brought back to the surface in thermally insulated boxes. *B. segonzaci* were collected using the slurp gun of the ROV and kept in 5 L bottles until the ROV recovery. On board, large animals were dissected to separate tissues and individually preserved in 80% ethanol, and/or directly used for DNA extractions. A hierarchical sampling scheme was implemented wherever possible, with two replicate sites sampled within each vent field (locality) and one to three vent fields sampled per basin, yielding a total of 21 sampling localities across vent communities (SI Table 1 & 2). For each locality and species, a minimum of 24 individuals were preserved or directly processed for DNA extraction, whenever sample availability allowed.

The sampling scheme was not fully achievable for *B. manusensis*, which was found only at Manus, Futuna, and one single Lau Basin site (Mangatolo) in sympatry with *Bathymodiolus septemdierum* (formerly known as *B. brevior*). It was absent at La Scala in the Woodlark Basin and southern Lau Basin sites along the North Fiji Ridge, where only *B. septemdierum* was present. DNA extractions were conducted directly on board for *I. nautilei*, *A. kojimai*, *B. manusensis*, *S. tollmanni*, *L. schrolli* & *L.* aff*. schrolli*, and *E. ohtai* from specific tissues (e.g., foot, mantle, whole body). For *B. segonzaci*, DNA extractions were performed later in the lab on ethanol-preserved samples. Extractions used either a modified CTAB 2%/PVPP 2% protocol (Jolly et al. 2003) or the NucleoSpin® Tissue 96 kit (Macherey-Nagel, Germany).

### Preparation of the ddRAD Libraries

The preparation of ddRAD genomic libraries was standardized by following the protocol described in Daguin-Thiébaut et al. (2021) and used for *I. nautilei* in Tran Lu Y et al. (2022). These seven libraries comprise 294 samples for *A. kojimai* (generated in Castel et al. (2022) and only raw data were reused), 469 individuals for *S. tollmanni*, 282 individuals for *E. ohtai*, 195 individuals for *B. manusensis*, 282 individuals for *B. segonzaci*, and 546 individuals for *L. schrolli & L.* aff. *schrolli*. All libraries were produced from gDNA digested with the enzymes *Pst1* and *Mse1,* except for *B. manusensis* that was digested with *Pst1* and *Msp1*.

These libraries also included 8 to 47 replicates (samples replicated twice or three times as controls) used for quality control and parameter calibration. Single-end (only for *B. manusensis*) or paired-end 150 (all other taxa) sequencing was performed on HiSeq 4000 (*I. nautilei*) or Novaseq 6000 Illumina (all other taxa), by the Genoscope, France (*I. nautilei*), or Novogene Europe (Cambridge, UK; all other taxa). The Fastqc Software (V.0.1.19) was used to check the sequence quality of the raw reads prior to the *de novo* assembly for each species.

A genomic assembly was performed for each species independently with the “*de novo”* Stacks2 module (Rochette et al. 2019) after demultiplexing individuals with the Process_radtags module. Parameter calibration followed the recommendations of (Mastretta-Yanes et al. 2015; Paris et al. 2017) (see SI Calibration). For all species, the parameter calibration, data filtering, and the Stacks modules used followed the methods described in (Tran Lu Y et al. 2022). However, for *S. tollmanni* and *L. schrolli* & *L.* aff. *schrolli,* adjustments were made to reduce the individual loss due to greater allele divergence (SI Table 2) (SI Table 3 & 4). A MAF filter at 0.05 was applied except for demographic inferences for which singletons were not filtered but masked .

### Population structure and admixture across the southwest Pacific

For each species, independent population genomics analyses were conducted to examine population structure, phylogeographic, and admixture patterns. Principal Component Analysis (PCA) was performed with SNPRelate (V.1.21.7) to assess spatial genetic diversity (Zheng et al. 2012). Admixture proportions were analyzed using Admixture (V.1.3.0) (Alexander and Lange 2011) with k values from 1 to 8 and ten runs per k. Population trees, including migration edges and F3 statistics, were generated in Treemix (V.1.13) (Pickrell and Pritchard 2012) with ten replicates and 0–5 migration events. The degree of genetic differentiation was estimated with pairwise *F*_st_ values between groups of individuals (genetic units or metapopulations, basins or localities) and with Analyses of Molecular Variance (AMOVA) with Arlequin (V.3.5.2.2) (Excoffier and Lischer 2010) and the statistical significance of *F*_st_ was assessed with 10,000 permutations of genotypes between populations.

For visualizations, all plots were generated using R (V.4.0.3) and ggplot2 (V.3.3.6). For *S. tollmanni*, individuals from Woodlark were subdivided in two groups based on genetic assignment (See Results).

The net divergence (*D_a_*) between populations was calculated from absolute divergence (*D_xy_*) corrected by average nucleotide diversity (*π*), following the (Nei and Li 1979) formula. Genetic diversity indices (*He*, *Ho, π*) were estimated for each genetic unit using the Stacks (V.2.52) population module across the final dataset. Indices were also estimated at basin scale.

### Evolutionary history of vent species and metapopulation connectivity

#### Relative gene flow direction

Gene flow patterns were assessed using the Divmigrate (Sundqvist et al. 2016) module within the R package diveRsity (V.1.9.90) (Keenan et al. 2013), which applies allele frequencies and *F*_st_ derived estimators to calculate a migration matrix normalized between 0 and 1 (where 1 indicates 100% gene flow and 0, none). No filter threshold was applied. Statistical significance of gene flow patterns between genetic units was tested using 1,000 bootstrap and non-overlapping 95% confidence intervals considered significant.

#### Demogenetic history of species metapopulations

To understand the demographic history of populations and identify the best model of population divergence, we utilized the ∂a∂i software (V2.1.0) (Gutenkunst et al. 2009) to fit joint allele frequency spectra to specific population models for each taxon independently. We focused on scenarios where an ancestral population splits into two daughter populations, with or without migration. This approach was suitable since all vent species analyzed present primarily a major subdivision into two genetic units (Eastern vs. Western populations).

One main advantage of using this ∂a∂i approach is its consideration of linked selection and heterogeneous migration across the genome, which are critical for accurate demographic inference (Ewing and Jensen 2016; Ravinet et al. 2017). To explore patterns of divergence and past and present genetic connectivity, we reused the approach used in (Tran Lu Y et al. 2022) for *I. nautilei* for all taxa. These models encompassed 28 potential scenarios derived from four major divergence models: Strict Isolation (SI), Isolation with Migration (IM), Ancient Migration (AM), and Secondary Contact (SC), initially developed in (Rougeux et al. 2017). The models accounted for various demographic and evolutionary processes, including changes in population size (G), the effects of barrier loci (from hybrid counterselection or local adaptation) through heterogeneous migration along the genome (2m), and linked selection (2N).

Due to the lack of external groups for allele state identification, we employed folded joint allele frequency spectra (folded JAFS). Each model was fitted at least 10 times independently for each species to assess convergence, and we compared models using the Akaike Information Criterion (AIC) for each simulation. Given the lack of information on the biology of these taxa, we estimated model parameters and divergence times (i.e., Ts since divergence, Tsc since secondary contact, and absolute divergence time) in generations using a uniform mutation rate of 10^-8^ (Lynch 2010; Popovic et al. 2023) across all species to facilitate comparisons of relative divergence times. Parameter uncertainties were calculated and reported at the 95% confidence level using the Fisher Information Matrix (FIM) on the best-fit model for each species.

## Results

### Calibration parameters and filtering steps

The 150 bp paired-end sequencing produced an average of 3.7, 3.3, 3.0, and 2.8 million paired reads per individual for *S. tollmanni*, *E. ohtai*, *B. segonzaci*, and *L. schrolli* & *L.* aff*. schrolli*, respectively. For *B. manusensis*, single read data yielded 2.9 million reads per individual. Raw reads for *A. kojimai* were sourced from (Castel et al. 2022), while *I. nautilei* results were directly reused from (Tran Lu Y et al. 2022).

We identified the optimal assembly parameters with Stacks (V.2.52) for each species, employing the de novo assembly pipeline for bi-allelic loci previously used for *I. nautilei* in (Tran Lu Y et al. 2022), After testing various parameter combinations, we selected assembly parameters ranging from 4 to 6 for m, 4 to 11 for M, and 5 to 11 for n, depending on the species (see SI Calibration & SI Table 5). Additionally, we applied the same methodologies, analyses, modules, and filtering parameters from (Tran Lu Y et al. 2022) to facilitate comparisons of genetic patterns across species (see SI Calibration). The *de novo* assemblies and filtering steps resulted in a variable number of SNPs, ranging from 2,904 to 47,547, derived from 159 to 414 individuals retained for each of the seven species (see SI Table 4 & 5).

### Population structure and admixture

Population structure analyses revealed consistent patterns in the spatial distribution of genetic diversity across the seven species. Principal Component Analysis (PCA) demonstrated a clear genetic separation between Manus individuals and those from the Eastern zones (North Fiji, Futuna, and Lau, hereafter NF/F/L) along the first component (PC1), explaining 1.94% to 26.03% of the total variance (Figure 2). Notably, the site La Scala on the Woodlark Ridge (discovered during the Chubacarc expedition, (Boulart et al. 2022)) shows contrasting results. First *I. nautilei*, *A. kojimai*, and *E. ohtai* were clearly divided into two genetic groups: Manus-Woodlark (M/W) and NF/F/L (Figure 2 A, B, D). Conversely, genetic relationships for Woodlark individuals differed slightly among other species. *S. tollmanni* displayed the two genetic groups, within Woodlark individuals, clustering either with Manus or NF/F/L, while one individual appeared admixed (Figure 2 C). For *L. schrolli* and *L.* aff. *schrolli*, all Woodlark individuals were positioned as intermediates between Manus (*L. schrolli*) and NF/F/L (*L.* aff. *schrolli*), being genetically closer to North Fiji individuals, which slightly diverged from the Futuna/Lau (F/L) individuals on PC1 (Figure 2 G). A similar pattern was also noted for *B. segonzaci*, with Woodlark individuals distinctly separated (but not intermediate) from both Manus and NF/F/L groups (i.e.closer to Manus: Figure 2 E). *B. manusensis*, while being absent from La Scala, displayed a similar differentiation pattern (Figure 2 F).

**Figure 2:**
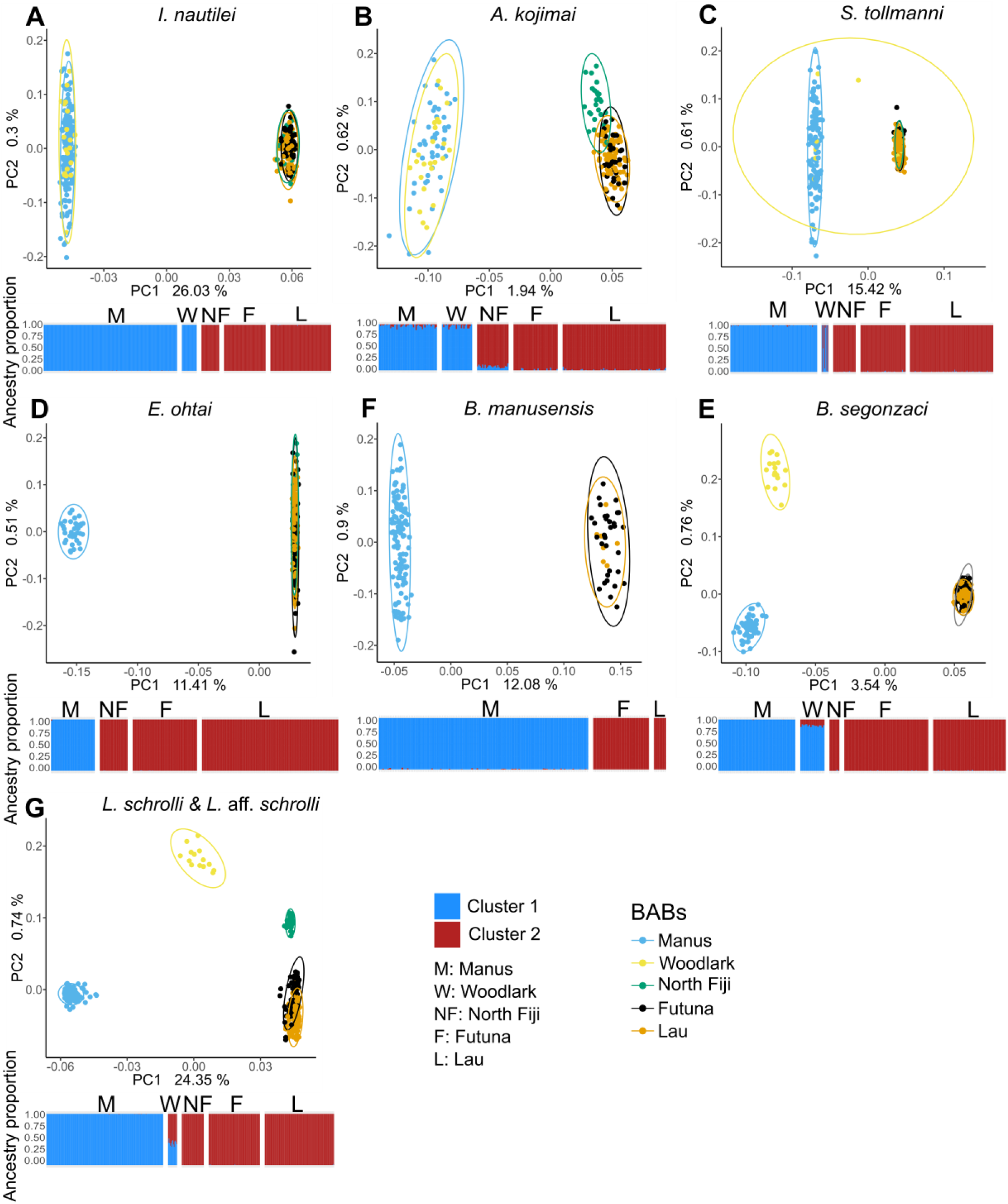
All PCA (PC1 & PC2) and Admixture plots for the best number of genetic clusters (K = 2) for each species (A, B, C, D, E, F, G). Colors in PCA plots represent regions (Manus, Woodlark, North Fiji and Lau Back-Arc-Basins, and the Futuna Volcanic Arc). Open ellipses represent the multivariate normal distribution of each group of points (basins) at 95% in PCA plots. Colors in Admixture plots represent each inferred genetic cluster. M: Manus, W: Woodlark, NF: North Fiji, F: Futuna, L: Lau.

The second component (PC2), accounting for 0.62-0.76% of the total genetic variance, revealed slight regional differences for some species (Figure 2 B, E, G). North Fiji individuals exhibited a subtle genetic differentiation from the F/L group on PC2 in *A. kojimai* and *L.* aff*. schrolli*, while *B. segonzaci* and *L. schrolli* showed a similar pattern between Woodlark and Manus individuals. Among the species analyzed, the *L. schrolli & L.* aff*. schrolli* complex exhibited the most pronounced basin-specific signature (Figure 2 G and SI Figure 1).

The shared distribution of genetic variation was corroborated by admixture analyses for all species, consistently identifying k = 2 as the optimal number of clusters (SI Figure 2). These genetic units correspond to M/W individuals on one side and NF/F/L on the other. Except for some individuals from Woodlark and North Fiji BABs, where genome admixture varied from 10% to 50% depending on the species, most individuals exhibited low levels of shared ancestry (Figure 2). Woodlark individuals displayed contributions from both genetic clusters for *L. schrolli/ L.* aff. *schrolli*, *S. tollmanni*, and *B. segonzaci*. In *S. tollmanni*, the Woodlark population comprised a mix of parental types and a putative F1 hybrid. Conversely, all Woodlark individuals showed approximately equal (∼50%) shared ancestry from both *L. schrolli* (Manus group) and *L.* aff. *schrolli* (Eastern group) when k = 2. For this later taxon, admixture analyses indicated additional clusters for Woodlark and North Fiji as k increased (k = 3 to 5, SI Figure 3 & 4).

F3 statistics revealed significant negative values, indicating admixture in Woodlark for two species. The first species, *S. tollmanni*, exhibited individuals from the Western population (Manus type: Woodlark2) with sources from both Manus (M) and Eastern (NF/F/L) groups, and the Eastern one (NF/F/L type: Woodlark1). The Woodlark *Lepetodrilus* individuals demonstrated a similar pattern of admixture, with sources from both Manus (M) and NF/F/L (SI Figure 5).

Analysis of overall genetic differentiation (*F*_st_) and net divergences (*D_a_*) between Western and Eastern groups revealed three main patterns (Table 1). The first group, comprising limpets *Lepetodrilus* and *Shinkailepas*, showed a high differentiation (0.271-0.360) and divergence (0.013-0.019). The second group (*I. nautilei*, *E. ohtai*, and *B. manusensis*) also exhibited high genetic differentiation (0.203-0.387) but lower to moderate divergences (0.002-0.007). The final group, including *B. segonzaci* and *A. kojimai*, had low genetic differentiation (0.018-0.038) and moderate divergences (0.007).

**Table 1:**
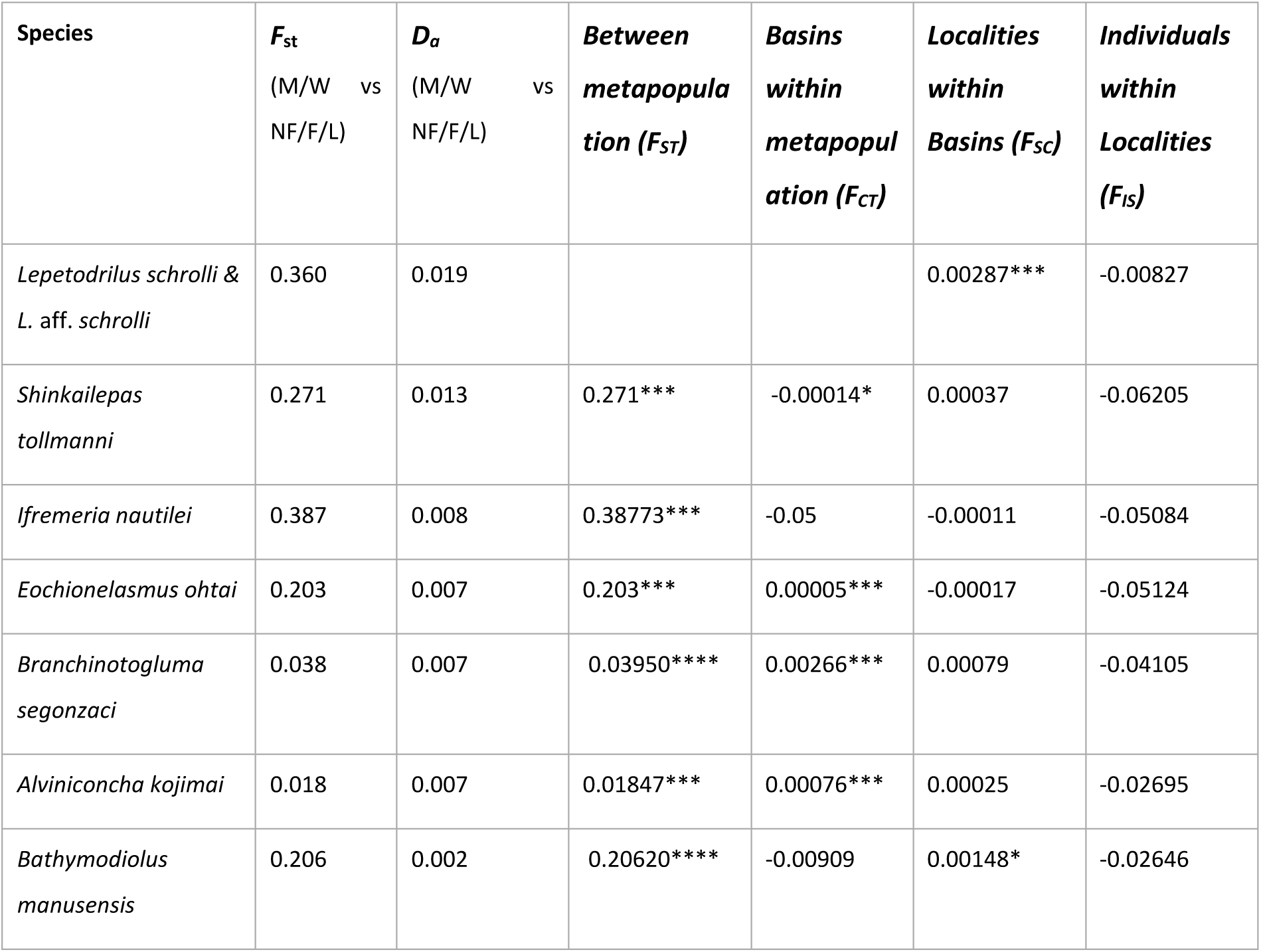
Fixation index (Fst) and net nucleotide divergence (Da) measured between the two (Eastern and Western) metapopulations for each species for the two first columns. Other columns display results of AMOVA analysis performed with Arlequin in a two step procedure (Basin within metapopulation and then Localities within basin). Significativity threshold for 10000 permutations for AMOVA (*:0.05, **:0.01, ***:<0.001)

When populations were divided into basins, pairwise *F*_st_ values indicated differentiation within each metapopulation, ranging from 0 to 0.364, depending on the species (see SI Table 6). Most significant genetic differentiation between BABs occurred in comparisons between Western and Eastern BABs (Manus or Woodlark against Lau, North Fiji, and Futuna). Some differentiation was also observed at the basin scale within groups, particularly the separation of North Fiji and Futuna/Lau populations for *A. kojimai* and *L.* aff*. schrolli.* However, no significant differentiation was noted between Lau and Futuna, except for *L.* aff*. schrolli*. These patterns are corroborated by the AMOVA results, which indicate that most variation occurs between metapopulations, with only minimal differentiation observed among certain basins within metapopulations (e.g., Woodlark and North Fiji) (Table 1). Notably, *L. schrolli* and *L.* aff. *schrolli* exhibited some little but significant variation among localities within basins.

Treemix analyses revealed a consistent two-group pattern of population differentiation across species, with varying optimal numbers of migration events (ranging from 0 to 2) between the Eastern and Western groups (SI Figure 6 and 7). This suggests a primary East to West migration event for all species, except *E. ohtai* and *B. manusensis*. Additionally, a secondary migration event was observed only for *I. nautilei*, *A. kojimai*, and *S. tollmanni*, occurring between two basins within the same genetic group (i.e., metapopulation), although the interacting basins varied across species. Notably, *B. segonzaci* displayed a distinct pattern with two migration events between the Eastern and Western groups: the first from Futuna to Woodlark and the second from Lau to Manus.

### Genetic diversity of species

Regardless of the statistics used (*Ho*, *He* and *π*: SI figure 8 & 9), *S. tollmani* displayed a gene diversity twice higher than that of the other species. The level of genetic diversity of species slightly differed between the Eastern and Western groups, but not always in the same direction. *I. nautilei*, *B. manusensis* and *L. schrolli/L.* aff. *schrolli* exhibited a slightly higher gene diversity in M/W compared with NF/F/L whereas it was the opposite for the other species (*A. kojimai*, *S. tollmani*, *E. ohtai* and *B. segonzaci*). These statistics displayed exactly the same pattern of distribution when calculated per basin (see SI Figure 9).

### Evolutionary history of populations and connectivity at the multispecies scale

#### Relative migration rates

The Divmigrate analysis between the Eastern and Western groups revealed a robust and common pattern of bidirectional but asymmetrical gene flow in all species. The main direction of gene flow is westward from NF/F/L (1) to M/W (2), while gene flow in the opposite direction was about half to a two-third (Figure 3). The species *L. schrolli/L.* aff. *schrolli*, however, displayed a more complex bidirectional pattern with much stronger gene flow between Futuna and Lau than between Futuna/Lau and North Fiji and virtually no gene flow between Woodlark, Manus and the others BABs.

**Figure 3:**
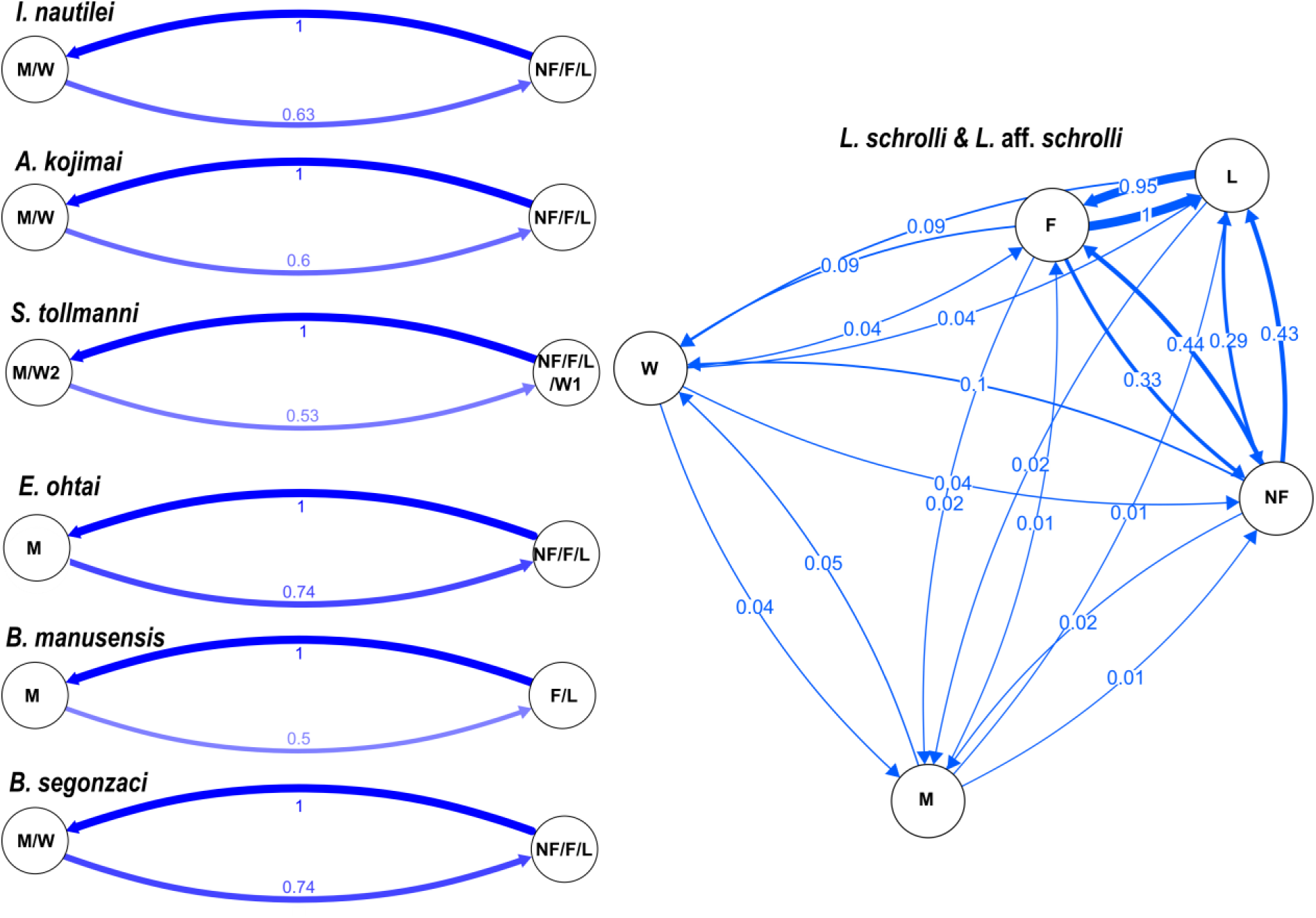
Relative migration (Nm) network plot between population estimated with Divmigrate. Populations are grouped into two metapopulations (East vs West) for each species, except for L. schrolli & L. aff. schrolli for which populations correspond to basins (L: Lau, F: Futuna, NF: North Fiji, W: Woodlark and M: Manus). M/W means Manus/Woodlark and NF/F/L North Fiji/Futuna/Lau. For *S. tollmanni*, the Woodlark population is subdivided between individuals genetically related to Lau/Futuna (W1) and Manus (W2), one F1 hybrid excluded from the analysis.

#### Demo-genetic inferences Modes of divergence

The folded JAFS (Figure 4 & SI Figure 10) showed the distribution of allelic variants between the Eastern and Western metapopulations for each species. Of the 28 models tested for each species, the SC (Secondary Contact) model was almost always the best-fitting model selected by the Weighted AIC (SI Figure 11 & 12). The exceptions are *I. nautilei* (Tran Lu Y et al. 2022) and *B. manusensis*, for which the SC model is slightly, but not significantly, better than the IM (Isolation with Migration) model. The Strict Isolation (SI) model was the worst model for all species. This indicated that all species are still able to maintain low levels of gene flow at the regional scale despite the large geographical distances and the abyssal plain separating the different regions. Increasing the complexity of the SC or IM population model by the addition of genomic heterogeneity (2N and/or 2m) or demographic change (G) parameters improved the model fit (see SI Figure 11 & 12). Capturing linked selection (2N) improved the model fit more than the G or 2m parameters for most species except *E. ohtai*. Alternatively, adding heterogeneous gene flow (2m) to the SC model improved the model fit for *E. ohtai* (SC2mG), although the two metapopulations appeared to be well separated. This situation of semi-permeable barrier also holds for *S. tollmanni* and *L. schrolli/L. aff. schrolli,* for which genetic admixture is locally suspected. For these two latter species but also for *A. kojimai*, *B. segonzaci*, and *I. nautilei,* the SC2N2mG was the best model after evaluating all possible parameter combinations. For *B. manusensis*, it was however not possible to discriminate between the models IM2N, SC2N, and SC2N2m, and both SC2mG and SC2NG performed similarly as well as the SC2N2mG model for *B. segonzaci* (Figure 4 and SI Figure 11 & 12).

**Figure 4:**
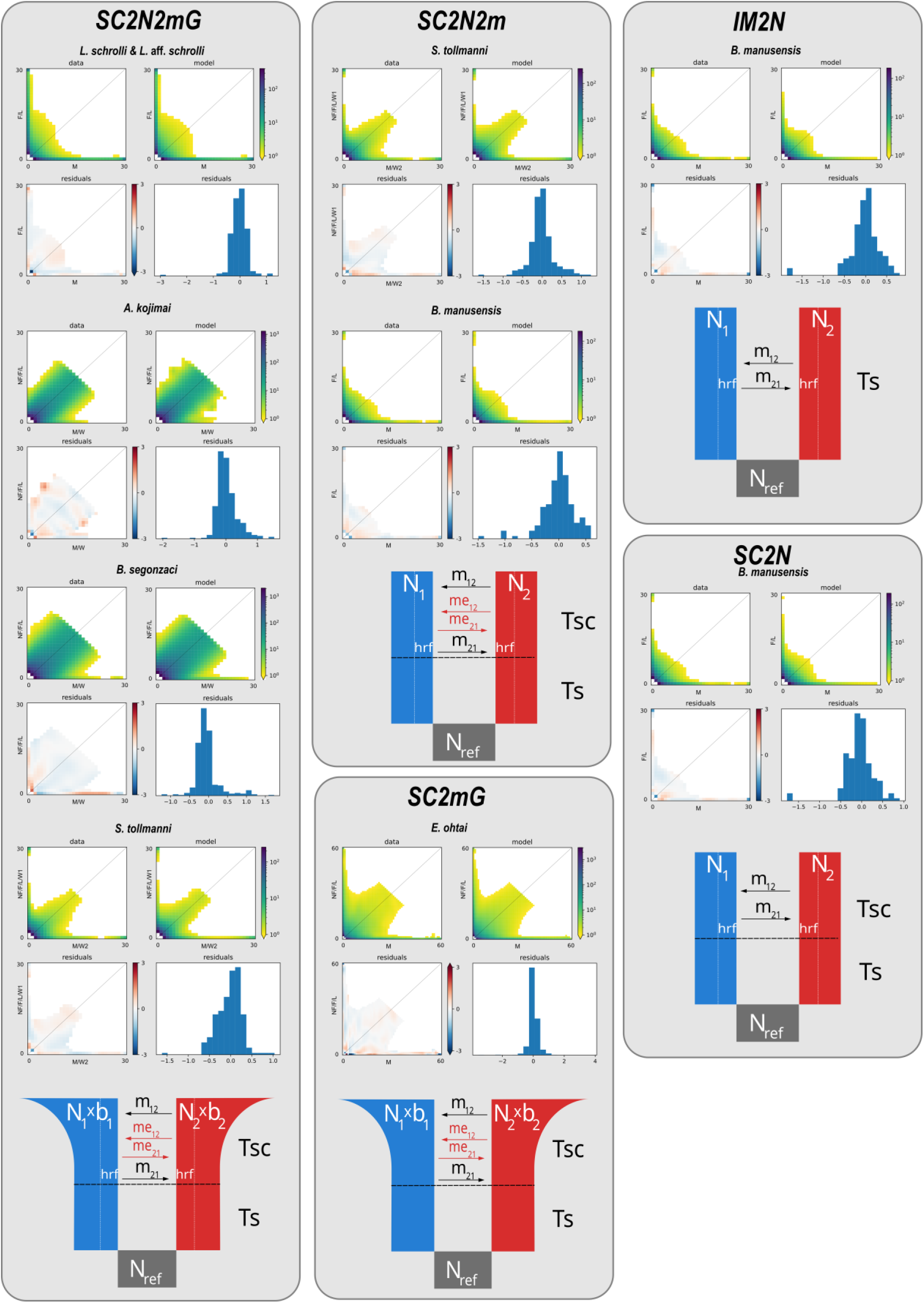
Folded Joint allele frequency spectrum (JAFS) plots for the best models (bottom) selected by dadi for all species according to the AIC as presented in SI figure 10*. All four plots represent the observed JAFS between populations (upper left), the simulated JAFS under the specified model (upper right), the log scale indicates SNP density in each frequency class. Residuals of data on the simulated JAFS (lower left) and histogram of their distribution (lower right)*

#### Timing of divergence and gene flow

Depending on the best model, we estimated divergence times: Ts (time since the population split), Tsc (time since secondary contact), and Ttotal (either Ts or Ts + Tsc), using a fixed mutation rate of 10^-8^ (Table 2). Ts ranged from 40,000 generations for *A. kojimai* to 116,000 for *L. schrolli and L.* aff. *schrolli*. Similarly, Tsc varied significantly, from 6,386 generations for *S. tollmanni* to 69,371 for *I. nautilei* and *L. schrolli*/*L.* aff. *schrolli*. The total time since the ancestral split (Ttotal) ranged from 40,892 generations for *A. kojimai* to 116,711 generations for *E. ohtai*. Notably, the divergence time for *B. manusensis* varied two-fold (43,644 to 101,718 generations) based on the model used, while *I. nautilei* and *S. tollmanni* yielded intermediate values.

**Table 2:** Estimation of divergence times in generations (estimates are made with a fixed mutation rate per generation and per site of 10^-8^ for all species) s. Ts: time since the ancestral population subdivided into two populations; Tsc: time since secondary contact. Ttotal: Ts and Ts + Tsc. For I. nautilei using the AM2N2mG model, Ts corresponds to the time of the strict split after ancient migration ended (Tsc).

| Species | Best model | *Ts* | *Tsc* | *Ttotal* |
| --- | --- | --- | --- | --- |
| *I. nautiliei* | IM2N2mG |  | 66 951 | 66 951 |
| *I. nautiliei* | SC2N2mG | 16 279 | 54 017 | 70 295 |
| *I. nautiliei* | AM2N2MG | 41 | 69 372 | 69 413 |
| *A. kojimai* | SC2N2mG | 29 157 | 11 735 | 40 892 |
| *S. tollmanni* | SC2N2m | 63 335 | 5 419 | 68 754 |
| *S. tollmanni* | SC2N2mG | 42 683 | 22 326 | 65 009 |
| *E. ohtai* | SC2mG | 37 278 | 56 793 | 94 071 |
| *B. segonzaci* | SC2N2mG | 18 194 | 15 375 | 33 569 |
| *B. manusensis* | IM2N | 101 718 |  | 101 718 |
| *B. manusensis* | SC2N2m | 40 752 | 2 892 | 43 644 |
| *B. manusensis* | SC2N | 56 394 | 37 940 | 94 335 |
| *L. schrolli/L. aff. schrolli* | SC2N2mG | 48 381 | 68 331 | 116 712 |

All species display heterogeneous gene flow (2m). Migration rate parameters estimated from *∂a∂i* show that present-day gene flow is stronger from NF/F/L to M/W (East to West) than the opposite, with the exception of *L. schrolli/L.* aff. *schrolli* (for both the two classes of gene flow “neutral” migration m and “reduced” (due to barrier loci) migration me, in relative genomic proportions P and 1-P; Table 3). For *B. manusensis*, migration rates were similar in both directions. *I. nautilei* and *E. ohtai* had about half of their genome characterized by reduced gene flow (P ∼=1-P). Only *A. kojimai* displayed high neutral gene flow (with m>me and P>>1-P) whereas *S. tollmanni, B. segonzaci, B. manusensis and L. schrolli/L.* aff. *schrolli)* exhibited a larger proportion of barrier loci (1-P >>P) which strongly reduced gene flow (me <<m) between the genome of the two genetic groups (Table 3).

**Table 3:** Gene flow parameters estimated from ∂a∂i (m and me are the “neutral” and “reduced” migration rate parameters. P the proportion of the genome characterized by migration rate m, and 1-P the proportion of the genome affected by reduced migration me). 1 : East population (NF/F/L); 2 : West population (M/W).

| Species | Best model | *m*1<-2 | *m*2<-1 | *P* | *me*1<-2 | *me*2<-1 | 1-*P* |
| --- | --- | --- | --- | --- | --- | --- | --- |
| *I. nautiliei* | IM2N2mG | 0,444 | 0,825 | 0,483 | 0,038 | 0,283 | 0,517 |
| *I. nautiliei* | SC2N2mG | 0,422 | 0,810 | 0,439 | 0,038 | 0,270 | 0,561 |
| *I. nautiliei* | AM2N2MG | 0,461 | 0,825 | 0,471 | 0,039 | 0,300 | 0,529 |
| *A. kojimai* | SC2N2mG | 4,741 | 4,389 | 0,851 | 0,384 | 0,841 | 0,149 |
| *S. tollmanni* | SC2N2m | 4,334 | 13,866 | 0,986 | 0,321 | 3,314 | 0,014 |
| *S. tollmanni* | SC2N2mG | 4.747 | 4.747 | 0,795 | 0,268 | 1,048 | 0,205 |
| *E. ohtai* | SC2mG | 0,364 | 1,386 | 0,508 | 0,013 | 0,124 | 0,492 |
| *B. segonzaci* | SC2N2mG | 2,195 | 5,412 | 1 | 0,579 | 6,489 | 0 |
| *B. manusensis* | IM2N | 0,436 | 0,566 |  |  |  |  |
| *B. manusensis* | SC2N2m | 3,988 | 4,891 | 0,323 | 0,227 | 0,646 | 0,677 |
| *B. manusensis* | SC2N | 0,520 | 0,537 |  |  |  |  |
| *L. schrolli/L. aff. schrolli* | SC2N2mG | 0,878 | 0,308 | 0,180 | 0,009 | 0,007 | 0,820 |

## Discussion

Population genomics allowed us to gain a deeper understanding of factors that have shaped genetic variation within and between populations in a context of a discontinuous ridge system in a well-delimited biogeographic region. Our study was aiming to “put the geography (and more) into comparative population genomics” (from Edwards et al. 2022) by applying such an approach to seven very different taxa strictly associated with the hydrothermal habitat. Using the same sampling scheme across all species, we show that these species share a common biogeographic break and a similar demographic history despite their different life-history traits. This pattern of differentiation was however slightly more complex for the limpets *L. schrolli & L.* aff. *schrolli* and *S. tollmanni*, and the scaleworm *B. segonzaci*, for which a further slight genetic subdivision between the Manus and Woodlark Basin was also observed.

The multispecies transition zone between the Eastern and Western metapopulations is located on the Woodlark Ridge or between it and the western part of the North Fiji BAB, isolating the Manus Basin to the West and the North Fiji and Lau Basins, as well as the Futuna volcanic arc to the East. Based on demogenetic inferences, we found that after a primary allopatric divergence, these metapopulations were secondarily connected, resulting in weak, asymmetric, and predominantly westward ongoing gene flow. Levels of divergence and differentiation, however, varied between taxa, probably because of different generation times and life-history traits. Heterogeneous gene flow was also found in all species in various proportions, suggesting that primary divergence could have led to the generation of genomic incompatibilities, as commonly found in hybrid zones (Matute et al. 2010; Bierne et al. 2011).

### A suture zone around the Woodlark back-arc Basin

All seven species share a clear genetic break into two main metapopulations across their geographical range, with the Manus/Woodlark (M/W) BABs on one hand and the North Fiji, and Lau BABs and Futuna Volcanic Arc (NF/F/L) on the other hand. Despite this common phylogeographic pattern, the strength of differentiation between these Eastern and Western genetic pools estimated by *F*_st_ varied from very high (0.387) to very low (0.020), depending on the species. A transition zone, i.e. a region where previously isolated lineages come into contact and exchange genetic material, appears to be located somewhere between North Fiji and Woodlark BAB or even at Woodlark itself, where both lineages are found in sympatry for some species. This is clearly observed for the two limpet species complexes, *S. tollmanni* and *L. schrolli & L.* aff. *schrolli*, for which the Western and Eastern lineages co-occur in Woodlark and hybrid individuals, are detected. This signature is typical of the existence of a tension zone, where selection can operate against hybrid genotypes, probably due to the existence of underdominant genetic incompatibilities (Bierne et al. 2011) or other reproductive barriers. Variable levels of admixture detected between Western and Eastern lineages in all species present at Woodlark, and, to a lesser extent at North Fiji (Tran Lu Y et al. 2022), reinforced this view. The species variability in terms of population structuring can stem from diverse origins: differences in life history traits (generation time, dispersal capability, fecundity, …) or in the genomic architecture of species (intensity of linked selection and number of barrier loci (genetic incompatibilities)). These latter factors are noticeably linked with the depth of evolutionary history of divergence and the reproductive mode. Our study suggests two possible hypotheses. First, geophysical rearrangements at a given time may have restricted, and still continue to limit the effective dispersal of several species according to their life history traits. This could have led to shared patterns of isolation, regardless of the timing of their origin (isolation with migration, IM, one of the models supported by *∂a∂i* for *Ifremeria nautilei* and *Bathymodiolus manusensis*). Alternatively, these species may have experienced one or more vicariance events, resulting in a period of primary divergence in allopatry followed (or not) by secondary contacts (SC) with some amount of gene flow. The latter scenario is the most likely, supported by the *∂a∂i* analyses for most species (Figure 4). This vicariance may have originated elsewhere in the Pacific and its very causes however remain to be identified.

While evolutionary processes unfold independently in each species, our demogenetic reconstruction indeed indicates a scenario of secondary contact (SC) for five species. For two species, the model cannot decisively distinguish between secondary contact (SC) and isolation-with-migration (IM), possibly due to a prolonged period of secondary contact. These results nevertheless strongly suggest an allopatric initial divergence for most species. Modeling also underscored the detection of a genomic heterogeneity of differentiation in all species, with both heterogeneous gene flow across the genome (2m) and linked selection (2N). This finding suggests that the initial divergence was extensive enough to generate genomic incompatibilities between populations in a genome characterized by substantial background selection and low recombination. This pattern can be a feature of hydrothermal-vent species, which may undergo strong purifying selection in this challenging environment (Chevaldonné et al. 2002; Fontanillas et al. 2017; Thomas-Bulle et al. 2022).

The contact zone between Western and Eastern lineages, presently located at Woodlark for some species and between Woodlark and the North Fiji Basin for others, may have shifted since secondary contact, likely moving along the Vanuatu or Solomon subduction arcs. Manus Basin appears more diversified in terms of species (see Poitrimol et al., 2022) and could have served as a source of biodiversity for part of the Western Pacific hydrothermal fauna, suggesting a possible “out of Manus” hypothesis. In line with this idea, recent work on taxa network analysis redescribes the Western Pacific region for hydrothermal fauna not as a single biogeographic province, as suggested by (Moalic et al. 2012), but as two distinct provinces, the North West Pacific and the South West Pacific, with the Manus Basin as a possible hub connecting both (Tunnicliffe et al. 2024).

### Gene flow between BABs

The connectivity and dispersal of hydrothermal vent species may be influenced by their life history traits, particularly larval phases. Species with planktotrophic larvae (if entrained in surface waters) are expected to have higher dispersal potential and greater connectivity across populations, whereas those with lecithotrophic larvae, which rely on yolk reserves, often show limited dispersal and connectivity even if lasting long in deep waters (Young, 1994). Our findings reveal an asymmetric gene flow, with less eastward gene flow, rejecting full isolation between Eastern and Western metapopulations regardless of contrasted dispersal traits. This aligns partly with prior research showing minimal gene flow between Manus and Lau BABs (Breusing et al., 2021; Plouviez et al., 2019; Thaler et al., 2011, 2014), though studies have also reported a lack of differentiation in *S. tollmanni* (Yahagi et al., 2019; Poitrimol et al., 2022).

While gene flow orientation corresponds partly with intermediate-depth larval dispersal models (Mitarai et al., 2016), the bidirectional flow suggests additional surface dispersal, where counter-currents flow eastward. The current limited biological knowledge, hinder assessments of larval development’s impact on population divergence and gene flow. Further study is needed, especially regarding intermediate populations in the Vanuatu and Solomon archipelagos. Varying secondary contact rates may have affected species differently; for instance, high-dispersal, long-generation species like *A. kojimai* and *B. segonzaci* show less divergence than low-dispersal, short-generation species like *L. schrolli* and *L.* aff. *schrolli*, a pattern echoed in Lau Basin vent copepods (Diaz-Recio Lorenzo et al., 2024). Nevertheless, present-day connectivity between the two metapopulations remains highly limited due to the existence of both genetic and physical barriers to dispersal.

### Species specific variation

The West/East divergence is clear across species, yet some also display basin-specific variations. *Branchinotogluma segonzaci* shows slight differentiation between Woodlark and Manus populations, not solely due to allelic introgression from other Eastern regions. Similarly, the North Fiji population of *A. kojimai* differs slightly from Lau/Futuna, despite a low regional genetic differentiation (*F*_st_= 0.018). *B. manusensis* was found east of its known range, co-occurring with *B. septemdierum*, the only mussel typically in that area, suggesting range expansion or relict population but without reduced diversity (Dupoué et al. 2021). These slight basin variations in *B. segonzaci* and *A. kojimai* imply potential limits on connectivity, likely influenced by factors like larval dispersal depth, demographic turnover, or putative effect of diversifying selection within certain populations.

While most species exhibit low genetic divergence, limpets *L. schrolli & L.* aff*. schrolli* and *S. tollmanni* show a high divergence and minimal gene flow between Western and Eastern groups. *S. tollmanni* exhibits admixture in Woodlark, where both lineages are sympatric, and one first-generation hybrid was identified. Previous studies found no mitochondrial *Cox1* differentiation (Yahagi et al. 2020; Poitrimol et al. 2022 on the same individuals used in this study), while our genomic data indicate a strong West/East lineage isolation with limited allelic exchange. This indicates a strong reduction in gene flow, but few loci can still be exchanged and captured by one of the two lineages, including the mitochondrial genome.

*Lepetodrilus schrolli & L.* aff. *schrolli* display comparable divergence but with a clearly admixed population in Woodlark between the Manus and NF/F/L lineages. This supports *Cox1* results, where half of Woodlark individuals reflect Manus haplotypes and the other half NF/F/L, consistent with previous nuclear marker studies showing low asymmetric gene flow between Manus and Lau (Plouviez et al. 2019; Poitrimol et al. 2022). These results suggest mitochondrial segregation in sympatry, partly supporting recent taxonomic revisions of *L. schrolli* and *L. fijiensis* (Chen and Sigwart 2023), however, our results show that intermediate populations exist, showing that this is in fact a complex of lineages undergoing speciation. Furthermore, this suggests that the taxonomy of *S. tollmanni* may also need to be revised.

Life-history traits may influence connectivity, though larval development type shows limited consistency. Despite their different larval modes, *B. segonzaci* (lecithotrophy) and *A. kojimai* (planktotrophy) display the lowest genetic differentiation and divergence between Eastern and Western populations.

*Branchinotogluma segonzaci* is a free-living, mobile annelid with small local populations and large mature oocytes (∼ 150 µm, SH unpublished data) suggesting lecithotrophic larval development that can persist in cold oligotrophic waters. *A. kojimai*, on the other hand, has larger, patchily-distributed populations and produces much smaller oocytes (∼ 90 µm), implying planktotrophic larvae that can reach the surface waters (Warèn and Bouchet 1993; Sommer et al. 2017; Hanson et al. 2024). Despite these differences, their low level of regional differentiation may be linked to the specific distribution of ‘hot’ vent emissions which both species likely disperse across.

*Bathymodiolus manusensis* and *E. ohtai* also show low divergence, but with moderate differentiation across metapopulations. *E. ohtai* is a common sessile hydrothermal vent cirriped forming dense populations in diffuse areas, and with large oocytes (∼ 500 µm, Yamaguchi and Newman 1997; Tyler and Young 1999) suggesting lecithotrophic development. In contrast, deep-sea bivalves such *as B. manusensis* typically have small oocytes (∼ 70 µm) and larvae are expected to be planktotrophic, as larvae of *Gigantidas childressi* (formerly known as *“Bathymodiolus*” *childressi*) that can remain in surface waters for extended periods (Arellano and Young 2009).

*Ifremeria nautilei* has lecithotrophic larvae incubated in a maternal pouch (Reynolds et al. 2010; Warèn & Bouchet 1993), likely limiting dispersal. While the larval mode in *S. tollmanni*, *L. schrolli* & *L.* aff*. schrolli* remains unknown, with large oocytes (∼ 150 µm; Poitrimol et al. 2024), other closely related *Shinkailepas* species suggest possible planktotrophy near the ocean surface (Yahagi et al. 2017), but with egg capsule incubation prior to release for *S. tollmanni* (Mouchi et al. 2024).

Though focused on the Southwest Pacific BABs, the Kermadec Basin—hundreds of miles south of Lau—harbors distinct vent fauna, *B. segonzaci* and *L.* aff. *schrolli* also appears there (SH unpublished data). This suggests long-term dispersal or greater habitat adaptability. Additional individuals from Kermadec showed no differentiation from NF/F/L *B. segonzaci* but some distinction in *L.* aff. *schrolli*, reinforcing regional population structure of this later (see SI Figures 13-15, SI Table 6 data not shown).

Our work demonstrates the challenges of linking hydrothermal species’ genetic structure solely to larval development, as other factors, such as population turnover, reproductive effort, and generation time, likely also shape patterns of population differentiation.

### Timing divergences and the hypothesis of vicariance

Our comparative study highlights shared phylogeographic patterns across the Southwest Pacific Ocean for seven vent species. This pattern likely stems from a common initial divergence event, due to either geological or climatic factors, generating two metapopulations, with the separation lying somewhere between Woodlark and North Fiji BABs, assuming the populations have remained in place during isolation. This barrier is semi permeable to gene flow, allowing some exchange of genetic material between metapopulations, through secondary contacts. This reconnection may also be modulated by species-specific life-history traits, including type of larvae, larval dispersal depth, longevity, reproduction timing, and habitat fragmentation. Net nucleotide divergence (*Da*) estimates, reveal that species have undergone different periods of divergence. Notably, the smallest species *L. schrolli & L.* aff. *schrolli* and *S. tollmanni* display higher net divergence values than other species, suggesting longer isolation but probably a greater number of generations since the split. This naturally raises question about cryptic species and speciation processes as divergence levels fall within the “grey zone” of speciation (Roux et al. 2016), where reproductive isolation between populations can vary widely. Our analyses not only reveal the gene flow intensity and heterogeneity (as previously discussed) but also raise interesting hypotheses regarding the timing of divergence.

Divergence times among species vary by a factor of three, influenced by unknown mutation rates and mean generation times, which we assumed to be uniform across species, though this is unlikely. Variations may be attributed to life-history traits, where larger species generally grow slower than smaller ones (Schöne and Giere 2005), indicating discrepancies in generation times and sexual maturity. Larger species also tend to have smaller populations, suggesting potential simultaneous demographic changes.

Estimates suggest that primary divergence began between approximately 40,892 and 101,718 generations (which may be easily explained by differences in generation times), while secondary contact ranges from 6,386 to 69,372 generations. Assuming a mutation rate of 10^-8^ and one generation per year, these events likely occurred during the Holocene amidst climatic oscillations, potentially starting around the Last Glacial Maximum (11,500–20,300 years ago for the Tongo Glaciation; 62,000 years ago, for the Komia Glaciation; 130,600– 158,000 years ago for the Mengane Glaciation) (Barrows et al. 2011). If we consider a tenfold lower mutation rate (10⁻⁹), common in molecular dating of vent fauna (Chevaldonné et al. 2002; Johnson et al. 2006; Matabos and Jollivet 2019), divergence estimates would range from 400,000 to 1,000,000 years, with secondary contact between 63,860 and 693,720 years.

Considering this later mutation rate, the primary divergence may coincide with magmatic accretion in recent and active back-arc ridges like Lau or Manus (Schellart et al. 2006). Geological structures between the two metapopulations, including the Woodlark and North Fiji basins, Vanuatu Trough, and the Solomon Islands volcanic arc, have older geotectonic histories, with accretion starting several million years ago (Woodlark: ∼6 Ma; North Fiji: ∼3 Ma; Vanuatu Trough and volcanic arc: ∼12 Ma; Solomon volcanic arc: Eocene, ∼40 Ma) (Schellart et al. 2006). While these timelines align with geological accretion, secondary contact likely occurred during the Holocene, influenced by climatic oscillations and changes in Pacific water-mass circulation.

Despite challenges in interpreting time estimates due to biological unknowns, divergence patterns suggest a significant climatic or geological event initiated a shared primary divergence followed by secondary contact for most species, with some species possibly experiencing older or prolonged secondary contact based on differing life-history traits.

### Limits

Our study primarily targeted the most common and emblematic species in hydrothermal communities of the West Pacific. Previous research on two of these species and others from these vent ecosystems has consistently supported our findings (Thaler et al. 2014; Lee et al. 2019; Plouviez et al. 2019; Poitrimol et al. 2022), revealing similar genetic differentiation patterns. However, these results may primarily reflect the evolutionary history of the most abundant and large vent species. Numerous other species with much lower densities and less reliance on vent fluids inhabit these ecosystems, potentially exhibiting more diverse and complex phylogeographic patterns. For instance, basins like Manus show higher species diversity and endemism (Boulart et al. 2022; Poitrimol et al. 2022; Diaz-Recio Lorenzo et al. 2024; Tunnicliffe et al. 2024).

Another key limitation from this sampling (mainly because of the difficulty to discover new sites) prevents us from conducting proper fine scale isolation-by-distance (IBD) analysis. IBD typically requires a quite uniform and continuous sampling distribution across the species’ range, which we were unable to achieve due to the spatial separation of hydrothermal vents and logistical constraints in sampling across such a vast and fragmented deep-sea landscape. In our case, the study area comprised a few unsampled - documented vents (e.g. Nifonea in Vanuatu), and other ‘ghost’ undiscovered vent sites possibly located on seamounts along volcanic arcs. Despite this limitation, the analysis using the current sampling scheme doesn’t display an Isolation by Distance (with the exception of *L. schrolli* & *L.* aff. *schrolli*) but rather a genetic cline (SI figure 16-23).

Additional data from these sites should not affect our main conclusions, but they would provide useful information to refine our patterns of population connectivity and the timing of contact zones. We also cannot rule out the idea that local adaptation may exacerbate the degree of genetic differentiation between the different hydrothermal populations analyzed, at least for some species. The study focused on the gastropod *Ifremeria nautilei* showed that analysis of genetic differentiation using outliers reinforces the isolation of the Woodlark Basin from Manus (Tran Lu Y et al. 2022), and depth may possibly have a filtering role on some alleles. Similarly, the East/West separation (Manus vs Lau) of hydrothermal populations in the Western Pacific is also accompanied by a change in the composition of hydrothermal fluids due to the nature of the rocks within these basins. It is therefore very difficult to tease apart the role of diversifying selection from other processes.

Distinguishing contemporary from historical gene flow remains challenging. Our demogenetic approach with ∂a∂i and *F*_st_-based metrics as used in Divmigrate, capture gene flow, through the use of allele frequencies which represent cumulative evolutionary forces effects over multiple generations rather than contemporary migration events (few generations ago). As a result, It is currently very difficult to distinguish between very recent and historical barriers and their associated factors.

### Implications for conservation and future directions

As previously shown, cases that correspond to geographically separated complex species probably need to be managed separately (e.g. *L. schrolli & L.* aff. *schrolli* or *S. tollmanni*). Other species depict much lower divergence but with some variation in population differentiation. Although sporadic and possibly rare, there is now evidence of present-day very low genetic connectivity between the Western and Eastern metapopulations with an apparent high genetic homogeneity within each of them.

Most of our knowledge on the stability of vent ecosystems through time is derived from times series established on the East Pacific Rise, a fast-spreading mid-oceanic ridge with a one-dimensional stepping-stone axis of colonization (Audzijonyte and Vrijenhoek 2010; Du Preez and Fisher 2018) and, some punctual physical barriers to dispersal (Plouviez et al. 2009; Plouviez et al. 2010; Plouviez et al. 2013). There, the fast extinction and recolonisation rates of active sites are likely to select species which can disperse far, and grow and reproduce fast. In back-arc basins, the ridge spreading rate is rather low but varies between basins (Dick 2019). Extinction and recolonisation events are likely less common, which led to concerns about the ability of the populations to recover if the metal sulfide deposits formed by the hydrothermal vent activity are mined (Du Preez and Fisher 2018). Within each of the two metapopulations, high genetic homogeneity of local populations can arise from either a substantial population size mitigating genetic drift or the presence of a sufficient number of migrants exchanged within BABs. For *I. nautilei*, *A. kojimai*, and *B. segonzaci*, introgressing alleles between metapopulations seem to reach only as far as the Woodlark and North Fiji BABs, suggesting that inter-basin dispersal alone may not compensate for population bottlenecks within each metapopulation. Consequently, dispersal may be effective between local sites at the scale of either the Western or Eastern regions, but much more limited between BABs. Because BAB zones are spatially limited with a restricted number of active vent sites, mining the already known sites should compromise any local ‘rescue’ effect.

## Conclusion

We identified a pronounced phylogeographic break across several hydrothermal species in the Southwest Pacific back-arc basins, centered between the Woodlark and North Fiji basins. While population structure patterns are shared, species show varying degrees of differentiation likely influenced by life history traits and species-specific demographic histories.

Although the timing of divergence and secondary contacts remains uncertain, connectivity between the two regions generally shows asymmetric, bidirectional gene flow favoring westward movement—except for *L. schrolli & L.* aff. *schrolli,* which may likely have originated from the Manus Basin. This study highlights genetic barriers at intermediate sites, which slow gene flow for certain species. Overall, vent species resilience seems more dependent on robust local population networks within each regional metapopulation than on long-distance dispersal. Thus, sustainable management of these communities requires conservation efforts at the biogeographic unit and at basin level, bearing in mind that the majority of current and low genetic exchange between the Eastern and Western basins are more specifically redirected towards the Manus Basin.

## Acknowledgements

The research was funded by the ANR CERBERUS project (ANR-17-CE02-0003). We would like to thank the captains and crews of the French research vessel *L’Atalante* and the ROV *Victor* team for the CHUBACARC cruise (Hourdez and Jollivet, 2019, https://doi.org/10.17600/18001111). Sampling would not have been possible without their dedication. Shiptime and scientist travels were supported by the Flotte Océanographique Française (FOF) and the Centre National de la Recherche Scientifique (CNRS). This work benefited from access to the Genomer and ABIMS platforms part of the Biogenouest network, at Station Biologique de Roscoff, an EMBRC-France and EMBRC-ERIC site. We also warmly thank Cindy L. Van Dover for sharing with us some of her Manus 2009 samples in order to perform preliminary ddRAD libraries over the different species to choose restriction enzymes and check the number of usable loci. We thank the reviewers for their contribution to improve the manuscript.

## Author contributions

Stephane Hourdez and Didier Jollivet designed the CHUBACARC and CERBERUS projects, François Bonhomme supervised the genetic work. Adrien Tran Lu Y, Stéphanie Ruault, Claire Daguin-Thiébaut, Anne-Sophie le Port, Marion Ballenghein, Sophie Arnaud-Haond, Jade Castel, Camille Poitrimol, Eric Thiébaut, François Lallier, Thomas Broquet, François Bonhomme, Didier Jollivet and Stéphane Hourdez performed laboratory work. Adrien Tran Lu Y performed bioinformatic statistical analyses with the contribution of François Bonhomme, Didier Jollivet, Pierre-Alexandre Gagnaire, Nicolas Bierne and Thomas Broquet. Adrien Tran Lu Y, François Bonhomme, Thomas Broquet, Didier Jollivet and Stephane Hourdez wrote the manuscript with feedback and inputs from all authors. All authors approved the manuscript

## Data availability statement

Raw sequence reads (Individual fastq files) are available at the European Nucleotide Archive (bioproject PRJEB47533; *I. nautilei*) and the NCBI sequence read archive (PRJNA768636 for A. kojimai; PRJNA779874 for *L. schrolli*; PRJNA772682 for *S. tollmanni*; PRJNA1044574 for *B. manusensis*; PRJNA1030156 for *E. ohtai*; PRJNA1044042 for *B. segonzaci*). Metadata relative to the samples are also available with Biosamples accessions and linked to the sequence reads accessions. Scripts used in this study (R, ∂a∂i) are available on a public Github repository: (https://github.com/Atranluy/Scripts-Ifremeria). VCFs and associated metadata will be available on public repository upon peer-review and publication (Dryad-XXXXXX).

## Benefit-sharing statement

In order to obtain the requested authorizations to work in national waters and in agreement with the Nagoya protocol, we contacted the authorities of the different countries (Papua-New Guinea, Fiji, and Tonga) and territories (Wallis and Futuna) for benefit sharing where sampling was performed. The data generated will be accessible on public databases (see above). The results obtained will also be communicated to these authorities which may have to make decisions regarding conservation of deep-sea hydrothermal vent communities in their EEZs. Observers for the different countries who took part in the on-board activities will be informed of our findings.

## Bibliography

1. Alexander DH, Lange K. 2011. Enhancements to the ADMIXTURE algorithm for individual ancestry estimation. BMC Bioinformatics 12:246. 10.1186/1471-2105-12-246

2. Arellano SM, Young CM. 2009. Spawning, Development, and the Duration of Larval Life in a deep-Sea cold-Seep mussel. The Biological Bulletin 216:149–162. https://www.journals.uchicago.edu/doi/10.1086/BBLv216n2p149

3. Audzijonyte A, Vrijenhoek RC. 2010. When Gaps Really Are Gaps: Statistical Phylogeography of hydrothermal vent invertebrates. Evolution 64:2369–2384. https://onlinelibrary.wiley.com/doi/abs/10.1111/j.1558-5646.2010.00987.x

4. Avise JC. 2000. Phylogeography: The History and Formation of Species. Harvard University Press 10.2307/j.ctv1nzfgj7

5. Avise JC. 2009. Phylogeography: retrospect and prospect. Journal of Biogeography 36:3–15. https://onlinelibrary.wiley.com/doi/abs/10.1111/j.1365-2699.2008.02032.x

6. Avise JC, Arnold J, Ball RM, Bermingham E, Lamb T, Neigel JE, Reeb CA, Saunders NC. 1987. INTRASPECIFIC PHYLOGEOGRAPHY: The Mitochondrial DNA Bridge Between Population Genetics and Systematics. Annual Review of Ecology and Systematics 18:489–522. 10.1146/annurev.es.18.110187.002421

7. Bachraty C, Legendre P, Desbruyères D. 2009. Biogeographic relationships among deep-sea hydrothermal vent faunas at global scale. Deep Sea Research Part I: Oceanographic Research Papers 56:1371–1378. https://www.sciencedirect.com/science/article/pii/S0967063709000314

8. Barrows TT, Hope GS, Prentice ML, Fifield LK, Tims SG. 2011. Late Pleistocene glaciation of the Mt Giluwe volcano, Papua New Guinea. Quaternary Science Reviews 30:2676–2689. https://linkinghub.elsevier.com/retrieve/pii/S0277379111001685

9. Barton N, Bengtsson BO. 1986. The barrier to genetic exchange between hybridising populations. Heredity 57:357–376. https://www.nature.com/articles/hdy1986135

10. Bierne N, Welch J, Loire E, Bonhomme F, David P. 2011. The coupling hypothesis: why genome scans may fail to map local adaptation genes. Molecular Ecology 20:2044–2072. https://onlinelibrary.wiley.com/doi/abs/10.1111/j.1365-294X.2011.05080.x

11. Boulart C, Rouxel O, Scalabrin C, Le Meur P, Pelleter E, Poitrimol C, Thiébaut E, Matabos M, Castel J, Tran Lu Y A, et al. 2022. Active hydrothermal vents in the Woodlark Basin may act as dispersing centres for hydrothermal fauna. Commun Earth Environ 3:1–16. https://www.nature.com/articles/s43247-022-00387-9

12. Breusing C, Johnson SB, Mitarai S, Beinart RA, Tunnicliffe V. 2021. Differential patterns of connectivity in Western Pacific hydrothermal vent metapopulations: A comparison of biophysical and genetic models. Evolutionary Applications. https://onlinelibrary.wiley.com/doi/abs/10.1111/eva.13326

13. Carver R, Childs J, Steinberg P, Mabon L, Matsuda H, Squire R, McLellan B, Esteban M. 2020. A critical social perspective on deep sea mining: Lessons from the emergent industry in Japan. Ocean & Coastal Management 193:105242. https://www.sciencedirect.com/science/article/pii/S0964569120301526

14. Castel J, Hourdez S, Pradillon F, Daguin-Thiébaut C, Ballenghien M, Ruault S, Corre E, Tran Lu Y A, Mary J, Gagnaire P-A, et al. 2022. Inter-Specific Genetic Exchange Despite Strong Divergence in Deep-Sea Hydrothermal Vent Gastropods of the Genus *Alviniconcha*. Genes 13:985. https://www.mdpi.com/2073-4425/13/6/985

15. Chen C, Sigwart JD. 2023. The lost vent gastropod species of Lothar A. Beck. Zootaxa 5270:401–436. https://www.mapress.com/zt/article/view/zootaxa.5270.3.2

16. Chevaldonné P, Jollivet D, Desbruyères D, Lutz R, Vrijenhoek R. 2002. Sister-species of eastern Pacific hydrothermal vent worms (*Ampharetidae, Alvinellidae, Vestimentifera*) provide new mitochondrial COI clock calibration. CBM - Cahiers de Biologie Marine 43:367–370. https://archimer.ifremer.fr/doc/00000/895/

17. De Jode A, Le Moan A, Johannesson K, Faria R, Stankowski S, Westram AM, Butlin RK, Rafajlović M, Fraïsse C. 2023. Ten years of demographic modelling of divergence and speciation in the sea. Evolutionary Applications 16:542–559. https://onlinelibrary.wiley.com/doi/abs/10.1111/eva.13428

18. Desbruyères D, Hashimoto J, Fabri M-C. 2006. Composition and Biogeography of Hydrothermal Vent Communities in Western Pacific Back-Arc Basins. In: Back-Arc Spreading Systems: Geological, Biological, Chemical, and Physical Interactions. American Geophysical Union (AGU). p. 215–234. https://onlinelibrary.wiley.com/doi/abs/10.1029/166GM11

19. Diaz-Recio Lorenzo C, Tran Lu Y A, Brunner O, Arbizu PM, Jollivet D, Laurent S, Gollner S. 2024. Highly structured populations of copepods at risk to deep-sea mining: Integration of genomic data with demogenetic and biophysical modelling. Molecular Ecology 33:e17340. https://onlinelibrary.wiley.com/doi/abs/10.1111/mec.17340

20. Dick GJ. 2019. The microbiomes of deep-sea hydrothermal vents: distributed globally, shaped locally. Nat Rev Microbiol 17:271–283. https://www.nature.com/articles/s41579-019-0160-2

21. Du Preez C, Fisher CR. 2018. Long-Term Stability of Back-Arc Basin Hydrothermal Vents. Front. Mar. Sci. 5. https://www.frontiersin.org/articles/10.3389/fmars.2018.00054

22. Dupoué A, Trochet A, Richard M, Sorlin M, Guillon M, Teulieres-Quillet J, Vallé C, Rault C, Berroneau Maud, Berroneau Matthieu, et al. 2021. Genetic and demographic trends from rear to leading edge are explained by climate and forest cover in a cold-adapted ectotherm. Diversity and Distributions 27:267–281. https://onlinelibrary.wiley.com/doi/abs/10.1111/ddi.13202

23. Edwards SV, Robin VV, Ferrand N, Moritz C. 2022. The Evolution of Comparative Phylogeography: Putting the Geography (and More) into Comparative Population Genomics. Genome Biology and Evolution 14:evab176. 10.1093/gbe/evab176

24. Ewing GB, Jensen JD. 2016. The consequences of not accounting for background selection in demographic inference. Molecular Ecology 25:135–141. https://onlinelibrary.wiley.com/doi/abs/10.1111/mec.13390

25. Excoffier L, Lischer HEL. 2010. Arlequin suite ver 3.5: a new series of programs to perform population genetics analyses under Linux and Windows. Molecular Ecology Resources 10:564–567. https://onlinelibrary.wiley.com/doi/abs/10.1111/j.1755-0998.2010.02847.x

26. Faure B, Jollivet D, Tanguy A, Bonhomme F, Bierne N. 2009. Speciation in the Deep Sea: Multi-Locus Analysis of Divergence and Gene Flow between Two Hybridizing Species of Hydrothermal Vent Mussels. PLOS ONE 4:e6485. https://journals.plos.org/plosone/article?id=10.1371/journal.pone.0006485

27. Fontanillas E, Galzitskaya OV, Lecompte O, Lobanov MY, Tanguy A, Mary J, Girguis PR, Hourdez S, Jollivet D. 2017. Proteome Evolution of Deep-Sea Hydrothermal Vent Alvinellid Polychaetes Supports the Ancestry of Thermophily and Subsequent Adaptation to Cold in Some Lineages. Genome Biology and Evolution 9:279–296. 10.1093/gbe/evw298

28. Gagnaire P-A. 2020. Comparative genomics approach to evolutionary process connectivity. Evolutionary Applications 13:1320–1334. https://onlinelibrary.wiley.com/doi/abs/10.1111/eva.12978

29. Gena K. 2013. Deep Sea Mining of Submarine Hydrothermal Deposits and its Possible Environmental Impact in Manus Basin, Papua New Guinea. Procedia Earth and Planetary Science 6:226–233. https://www.sciencedirect.com/science/article/pii/S1878522013000325

30. Gutenkunst RN, Hernandez RD, Williamson SH, Bustamante CD. 2009. Inferring the Joint Demographic History of Multiple Populations from Multidimensional SNP Frequency Data. PLOS Genetics 5:e1000695. https://journals.plos.org/plosgenetics/article?id=10.1371/journal.pgen.1000695

31. Hamilton WD, May RM. 1977. Dispersal in stable habitats. Nature 269:578–581. https://www.nature.com/articles/269578a0

32. Hanson N, Bates A, Dufour S. 2024. Reproductive biology of the original hydrothermal hairy snail, *Alviniconcha hessleri*, from the Mariana back-arc. https://www.researchsquare.com/article/rs-4883307/v1

33. Hickerson MJ, Carstens BC, Cavender-Bares J, Crandall KA, Graham CH, Johnson JB, Rissler L, Victoriano PF, Yoder AD. 2010. Phylogeography’s past, present, and future: 10 years after Avise, 2000. Molecular Phylogenetics and Evolution 54:291–301. https://linkinghub.elsevier.com/retrieve/pii/S105579030900373X

34. Hourdez S, Jollivet D. 2019. CHUBACARC Cruise Report. https://hal.sorbonne-universite.fr/hal-04100305v1

35. Hourdez S, Jollivet D. 2020. Chapter 2. In: Metazoan adaptation to deep-sea hydrothermal vents. G. di Prisco, A. Huiskes and C. Ellis-Evanz (eds.). Ecological reviews, Cambridge Univ. Press. p. 42–67.

36. Hurtado LA, Lutz RA, Vrijenhoek RC. 2004. Distinct patterns of genetic differentiation among annelids of eastern Pacific hydrothermal vents. Molecular Ecology 13:2603–2615. https://onlinelibrary.wiley.com/doi/abs/10.1111/j.1365-294X.2004.02287.x

37. Johnson SB, Won Y-J, Harvey JB, Vrijenhoek RC. 2013. A hybrid zone between *Bathymodiolus* mussel lineages from eastern Pacific hydrothermal vents. BMC Evol Biol 13:21. 10.1186/1471-2148-13-21

38. Johnson SB, Young CR, Jones WJ, Warén A, Vrijenhoek RC. 2006. Migration, Isolation, and Speciation of Hydrothermal Vent Limpets (*Gastropoda; Lepetodrilidae*) Across the Blanco Transform Fault. The Biological Bulletin 210:140–157. https://www.journals.uchicago.edu/doi/10.2307/4134603

39. Jolly M, Viard F, Weinmayr G, Gentil F, Thiébaut E, Jollivet D. 2003. Does the genetic structure of *Pectinaria koreni (Polychaeta: Pectinariidae*) conform to a source–sink metapopulation model at the scale of the Baie de Seine? Helgol Mar Res 56:238–246. https://hmr.biomedcentral.com/articles/10.1007/s10152-002-0123-1

40. Keenan K, McGinnity P, Cross TF, Crozier WW, Prodöhl PA. 2013. diveRsity: An R package for the estimation and exploration of population genetics parameters and their associated errors. Methods in Ecology and Evolution 4:782–788. https://onlinelibrary.wiley.com/doi/abs/10.1111/2041-210X.12067

41. Lee W-K, Kim S-J, Hou BK, Dover CLV, Ju S-J. 2019. Population genetic differentiation of the hydrothermal vent crab *Austinograea alayseae (Crustacea: Bythograeidae*) in the Southwest Pacific Ocean. PLOS ONE 14:e0215829. https://journals.plos.org/plosone/article?id=10.1371/journal.pone.0215829

42. Levin LA. 1990. A review of methods for labeling and tracking marine invertebrate larvae. Ophelia 32:115–144. 10.1080/00785236.1990.10422028

43. Lynch M. 2010. Evolution of the mutation rate. Trends in Genetics 26:345–352. https://www.sciencedirect.com/science/article/pii/S0168952510001034

44. Mastretta-Yanes A, Arrigo N, Alvarez N, Jorgensen TH, Piñero D, Emerson BC. 2015. Restriction site-associated DNA sequencing, genotyping error estimation and de novo assembly optimization for population genetic inference. Molecular Ecology Resources 15:28–41. https://onlinelibrary.wiley.com/doi/abs/10.1111/1755-0998.12291

45. Matabos M, Jollivet D. 2019. Revisiting the *Lepetodrilus elevatus* species complex (*Vetigastropoda: Lepetodrilidae*), using samples from the Galápagos and Guaymas hydrothermal vent systems. Journal of Molluscan Studies 85:154–165. 10.1093/mollus/eyy061

46. Matabos M, Plouviez S, Hourdez S, Desbruyères D, Legendre P, Warén A, Jollivet D, Thiébaut E. 2011. Faunal changes and geographic crypticism indicate the occurrence of a biogeographic transition zone along the southern East Pacific Rise. Journal of Biogeography 38:575–594. https://onlinelibrary.wiley.com/doi/abs/10.1111/j.1365-2699.2010.02418.x

47. Matute DR, Butler IA, Turissini DA, Coyne JA. 2010. A Test of the Snowball Theory for the Rate of Evolution of Hybrid Incompatibilities. Science 329:1518–1521. https://www.science.org/doi/full/10.1126/science.1193440

48. McPeek MA, Holt RD. 1992. The Evolution of Dispersal in Spatially and Temporally Varying Environments. The American Naturalist 140:1010–1027. https://www.journals.uchicago.edu/doi/abs/10.1086/285453

49. Mitarai S, Watanabe H, Nakajima Y, Shchepetkin AF, McWilliams JC. 2016. Quantifying dispersal from hydrothermal vent fields in the western Pacific Ocean. Proceedings of the National Academy of Sciences 113:2976–2981. http://www.pnas.org/lookup/doi/10.1073/pnas.1518395113

50. Moalic Y, Desbruyères D, Duarte CM, Rozenfeld AF, Bachraty C, Arnaud-Haond S. 2012. Biogeography Revisited with Network Theory: Retracing the History of Hydrothermal Vent Communities. *Syst Biol* 61:127–127. https://academic.oup.com/sysbio/article/61/1/127/1677959

51. Mouchi V, Pecheyran C, Claverie F, Cathalot C, Matabos M, Germain Y, Rouxel O, Jollivet D, Broquet T, Comtet T. 2024. A step towards measuring connectivity in the deep sea: elemental fingerprints of mollusk larval shells discriminate hydrothermal vent sites. Biogeosciences 21:145–160. https://bg.copernicus.org/articles/21/145/2024/

52. Nei M, Li WH. 1979. Mathematical model for studying genetic variation in terms of restriction endonucleases. PNAS 76:5269–5273. https://www.pnas.org/content/76/10/5269

53. Niner HJ, Ardron JA, Escobar EG, Gianni M, Jaeckel A, Jones DOB, Levin LA, Smith CR, Thiele T, Turner PJ, et al. 2018. Deep-Sea Mining With No Net Loss of Biodiversity—An Impossible Aim. Frontiers in Marine Science 5. https://www.frontiersin.org/article/10.3389/fmars.2018.00053

54. Papadopoulou A, Knowles LL. 2016. Toward a paradigm shift in comparative phylogeography driven by trait-based hypotheses. Proceedings of the National Academy of Sciences 113:8018– 8024. https://www.pnas.org/doi/abs/10.1073/pnas.1601069113

55. Paris JR, Stevens JR, Catchen JM. 2017. Lost in parameter space: a road map for stacks. Methods in Ecology and Evolution 8:1360–1373. https://besjournals.onlinelibrary.wiley.com/doi/abs/10.1111/2041-210X.12775

56. Pickrell JK, Pritchard JK. 2012. Inference of Population Splits and Mixtures from Genome-Wide Allele Frequency Data. PLOS Genetics 8:e1002967. https://journals.plos.org/plosgenetics/article?id=10.1371/journal.pgen.1002967

57. Plouviez S, LaBella AL, Weisrock DW, Meijenfeldt FAB von, Ball B, Neigel JE, Dover CLV. 2019. Amplicon sequencing of 42 nuclear loci supports directional gene flow between South Pacific populations of a hydrothermal vent limpet. Ecology and Evolution 9:6568–6580. https://onlinelibrary.wiley.com/doi/abs/10.1002/ece3.5235

58. Plouviez S, Le Guen D, Lecompte O, Lallier FH, Jollivet D. 2010. Determining gene flow and the influence of selection across the equatorial barrier of the East Pacific Rise in the tube-dwelling polychaete *Alvinella pompejana*. BMC Evolutionary Biology 10:220. 10.1186/1471-2148-10-220

59. Plouviez S, Schultz TF, McGinnis G, Minshall H, Rudder M, Van Dover CL. 2013. Genetic diversity of hydrothermal-vent barnacles in Manus Basin. Deep Sea Research Part I: Oceanographic Research Papers 82:73–79. https://www.sciencedirect.com/science/article/pii/S0967063713001805

60. Plouviez S, Shank TM, Faure B, Daguin-Thiebaut C, Viard F, Lallier FH, Jollivet D. 2009. Comparative phylogeography among hydrothermal vent species along the East Pacific Rise reveals vicariant processes and population expansion in the South. Molecular Ecology 18:3903–3917. https://onlinelibrary.wiley.com/doi/abs/10.1111/j.1365-294X.2009.04325.x

61. Poitrimol C, Matabos M, Veuillot A, Ramière A, Comtet T, Boulart C, Cathalot C, Thiébaut É. 2024. Reproductive biology and population structure of three hydrothermal gastropods *(Lepetodrilus schrolli*, L. fijiensis and Shinkailepas tollmanni) from the South West Pacific back-arc basins. Mar Biol 171:31. 10.1007/s00227-023-04348-4

62. Poitrimol C, Thiébaut É, Daguin-Thiébaut C, Port A-SL, Ballenghien M, Y ATL, Jollivet D, Hourdez S, Matabos M. 2022. Contrasted phylogeographic patterns of hydrothermal vent gastropods along South West Pacific: Woodlark Basin, a possible contact zone and/or stepping-stone. PLOS ONE 17:e0275638. https://journals.plos.org/plosone/article?id=10.1371/journal.pone.0275638

63. Popovic I, Bergeron LA, Bozec Y-M, Waldvogel A-M, Howitt SM, Damjanovic K, Patel F, Cabrera MG, Wörheide G, Uthicke S, et al. 2023. High germline mutation rates but not extreme population size outbreaks influence genetic diversity in crown-of-thorns sea stars. PLOS Genetics 20(2): e1011129. 10.1371/journal.pgen.1011129

64. Ravinet M, Faria R, Butlin RK, Galindo J, Bierne N, Rafajlović M, Noor M a. F, Mehlig B, Westram AM. 2017. Interpreting the genomic landscape of speciation: a road map for finding barriers to gene flow. Journal of Evolutionary Biology 30:1450–1477. https://onlinelibrary.wiley.com/doi/abs/10.1111/jeb.13047

65. Rochette NC, Rivera-Colón AG, Catchen JM. 2019. Stacks 2: Analytical methods for paired-end sequencing improve RADseq-based population genomics. Molecular Ecology 28:4737–4754. https://onlinelibrary.wiley.com/doi/abs/10.1111/mec.15253

66. Rougeux C, Bernatchez L, Gagnaire P-A. 2017. Modeling the Multiple Facets of Speciation-with-Gene-Flow toward Inferring the Divergence History of Lake Whitefish Species Pairs (*Coregonus clupeaformis*). Genome Biol Evol 9:2057–2074. https://academic.oup.com/gbe/article/9/8/2057/4060520

67. Roux C, Fraïsse C, Romiguier J, Anciaux Y, Galtier N, Bierne N. 2016. Shedding Light on the Grey Zone of Speciation along a Continuum of Genomic Divergence. PLOS Biology 14:e2000234. https://journals.plos.org/plosbiology/article?id=10.1371/journal.pbio.2000234

68. Schellart WP, Lister GS, Toy VG. 2006. A Late Cretaceous and Cenozoic reconstruction of the Southwest Pacific region: Tectonics controlled by subduction and slab rollback processes. Earth-Science Reviews 76:191–233. https://www.sciencedirect.com/science/article/pii/S001282520600016X

69. Schöne BR, Giere O. 2005. Growth increments and stable isotope variation in shells of the deep-sea hydrothermal vent bivalve mollusk *Bathymodiolus brevior* from the North Fiji Basin, Pacific Ocean. Deep Sea Research Part I: Oceanographic Research Papers 52:1896–1910. https://www.sciencedirect.com/science/article/pii/S0967063705001330

70. Sommer SA, Van Woudenberg L, Lenz PH, Cepeda G, Goetze E. 2017. Vertical gradients in species richness and community composition across the twilight zone in the North Pacific Subtropical Gyre. Molecular Ecology 26:6136–6156. https://onlinelibrary.wiley.com/doi/abs/10.1111/mec.14286

71. Sundqvist L, Keenan K, Zackrisson M, Prodöhl P, Kleinhans D. 2016. Directional genetic differentiation and relative migration. Ecology and Evolution 6:3461–3475. https://onlinelibrary.wiley.com/doi/abs/10.1002/ece3.2096

72. Teixeira S, Serrão EA, Arnaud-Haond S. 2012. Panmixia in a Fragmented and Unstable Environment: The Hydrothermal Shrimp *Rimicaris exoculata* Disperses Extensively along the Mid-Atlantic Ridge. PLOS ONE 7:e38521. https://journals.plos.org/plosone/article?id=10.1371/journal.pone.0038521

73. Thaler AD, Plouviez S, Saleu W, Alei F, Jacobson A, Boyle EA, Schultz TF, Carlsson J, Dover CLV. 2014. Comparative Population Structure of Two Deep-Sea Hydrothermal-Vent-Associated Decapods (*Chorocaris sp. 2* and *Munidopsis lauensis*) from Southwestern Pacific Back-Arc Basins. PLOS ONE 9:e101345. https://journals.plos.org/plosone/article?id=10.1371/journal.pone.0101345

74. Thaler AD, Zelnio K, Saleu W, Schultz TF, Carlsson J, Cunningham C, Vrijenhoek RC, Van Dover CL. 2011. The spatial scale of genetic subdivision in populations of *Ifremeria nautilei*, a hydrothermal-vent gastropod from the southwest Pacific. BMC Evolutionary Biology 11:372. http://bmcevolbiol.biomedcentral.com/articles/10.1186/1471-2148-11-372

75. Thomas-Bulle C, Bertrand D, Nagarajan N, Copley RR, Corre E, Hourdez S, Bonnivard É, Claridge-Chang A, Jollivet D. 2022. Genomic patterns of divergence in the early and late steps of speciation of the deep-sea vent thermophilic worms of the genus *Alvinella*. BMC Ecol Evo 22:106. 10.1186/s12862-022-02057-y

76. Tran Lu Y A, Ruault S, Daguin-Thiébaut C, Castel J, Bierne N, Broquet T, Wincker P, Perdereau A, Arnaud-Haond S, Gagnaire P-A, et al. 2022. Subtle limits to connectivity revealed by outlier loci within two divergent metapopulations of the deep-sea hydrothermal gastropod *Ifremeria nautilei*. Molecular Ecology 31:2796–2813. https://onlinelibrary.wiley.com/doi/abs/10.1111/mec.16430

77. Tunnicliffe V. 1992. The Nature and Origin of the Modern Hydrothermal Vent Fauna. PALAIOS 7:338–350. https://www.jstor.org/stable/3514820

78. Tunnicliffe V, Chen C, Giguère T, Rowden AA, Watanabe HK, Brunner O. 2024. Hydrothermal vent fauna of the western Pacific Ocean: Distribution patterns and biogeographic networks. Diversity and Distributions 30:e13794. https://onlinelibrary.wiley.com/doi/abs/10.1111/ddi.13794

79. Tyler PA, Young CM. 1999. Reproduction and dispersal at vents and cold seeps. Journal of the Marine Biological Association of the United Kingdom 79:193–208. https://www.cambridge.org/core/journals/journal-of-the-marine-biological-association-of-the-united-kingdom/article/abs/reproduction-and-dispersal-at-vents-and-cold-seeps/FACD4AC999965AC706C719FC849BAF8F

80. Van Dover CL. 2011. Mining seafloor massive sulphides and biodiversity: what is at risk? ICES Journal of Marine Science 68:341–348. 10.1093/icesjms/fsq086

81. Van Dover CL, Ardron JA, Escobar E, Gianni M, Gjerde KM, Jaeckel A, Jones DOB, Levin LA, Niner HJ, Pendleton L, et al. 2017. Biodiversity loss from deep-sea mining. Nature Geosci 10:464–465. https://www.nature.com/articles/ngeo2983

82. Vrijenhoek RC. 2010. Genetic diversity and connectivity of deep-sea hydrothermal vent metapopulations. Molecular Ecology 19:4391–4411. https://onlinelibrary.wiley.com/doi/abs/10.1111/j.1365-294X.2010.04789.x

83. Warèn A, Bouchet P. 1993. New records, species, genera, and a new family of gastropods from hydrothermal vents and hydrocarbon seeps*. Zoologica Scripta 22:1–90. https://onlinelibrary.wiley.com/doi/abs/10.1111/j.1463-6409.1993.tb00342.x

84. Washburn TW, Turner PJ, Durden JM, Jones DOB, Weaver P, Van Dover CL. 2019. Ecological risk assessment for deep-sea mining. Ocean & Coastal Management 176:24–39. https://www.sciencedirect.com/science/article/pii/S096456911830838X

85. Yahagi T, Thaler AD, Dover CLV, Kano Y. 2020. Population connectivity of the hydrothermal-vent limpet *Shinkailepas tollmanni* in the Southwest Pacific (*Gastropoda: Neritimorpha: Phenacolepadidae*). PLOS ONE 15:e0239784. https://journals.plos.org/plosone/article?id=10.1371/journal.pone.0239784

86. Yamaguchi T, Newman WA. 1997. The hydrothermal vent barnacle Eochionelasmus (Cirripedia, Balanomorpha) from the North Fiji, Lau and Manus Basins, South-West Pacific. Zoosystema 19 (4) : 623–649 https://sciencepress.mnhn.fr/en/periodiques/zoosystema/19/4/hydrothermal-vent-barnacles-eochionelasmus-cirripedia-balanomorpha-north-fiji-lau-and-manus-basins-south-west-pacific

87. Young CM, Eckelbarger KJ, Eckelbarger K. 1994. Reproduction, Larval Biology, and Recruitment of the Deep-sea Benthos. Columbia University Press

88. Zheng X, Levine D, Shen J, Gogarten SM, Laurie C, Weir BS. 2012. A high-performance computing toolset for relatedness and principal component analysis of SNP data. Bioinformatics 28:3326–3328. 10.1093/bioinformatics/bts606

