## Supplementary_Information for "Comparative population genomics unveils congruent secondary suture zone in Southwest Pacific Hydrothermal Vents"

### Tables

*Table 1: Sample details for all species used in this study, including the number of replicates. All samples have been collected during the Chubacarc 2019 campaign and a few samples from S.Hourdez (SH) and C.L Van Dover's (CDV) collections from 2009. I. nautilei population information can be found in Tran Lu Y et al., 2022. A hierarchical sampling has been followed with 3 levels of hierarchy: sites, localities and basins. N° samples is the number of individuals processed per site. N° libraries represent the total number of RAD libraries (including replicates) per site. Depth is in meters. Species short names: A_koj = Alviniconcha kojimai; B_m = Bathymodiolus manusensis; B_seg = Branchinotogluma segonzaci; E_ohtai = Eochionelasmus ohtai; L_affschro = Lepetodrilus aff. schrolli; L_schro = Lepetodrilus schrolli; S_toll = Shinkailepas tollmanni.*

| Basin | Locality | ID_Site | N° sample | N° libraries | Species | Longitude | Latitude | Depth |
|---|---|---|---|---|---|---|---|---|
| **Futuna** | Fati_Ufu | 12 | 19 | 19 | A_koj | W 177 11.082 | S 14 45.589 | 1519 |
| **Futuna** | Fati_Ufu | 14 | 3 | 3 | A_koj | W 177 11.098 | S 14 45.587 | 1518 |
| **Futuna** | Fati_Ufu | 18 | 20 | 23 | A_koj | W 177 11.116 | S 14 45.597 | 1519 |
| **Futuna** | Fatu_Kapa | 25 | 4 | 4 | A_koj | W 177 09.132 | S 14 45.109 | 1562 |
| **Futuna** | Fatu_Kapa | 30 | 14 | 14 | A_koj | W 177 09.960 | S 14 44.245 | 1547 |
| **Lau** | Mangatolo | 46 | 3 | 3 | A_koj | W 174 39.210 | S 15 24.875 | 2031 |
| **Lau** | Mangatolo | 51 | 18 | 18 | A_koj | W 174 39.331 | S 15 24.961 | 2039 |
| **Lau** | Abe | 6 | 2 | 2 | A_koj | W 176 11.481 | S 20 45.785 | 2149 |
| **Lau** | Tow_Cam | 74 | 21 | 23 | A_koj | W 176 08.257 | S 20 19.074 | 2716 |
| **Lau** | Tow_Cam | 77 | 19 | 21 | A_koj | W 176 08.264 | S 20 19.085 | 2711 |
| **Lau** | Tui_Malila | 78 | 20 | 21 | A_koj | W 176 34.070 | S 21 59.254 | 1899 |
| **Lau** | Tui_Malila | 83 | 12 | 12 | A_koj | W 176 34.091 | S 21 59.356 | 1884 |
| **Lau** | Tui_Malila | 86 | 16 | 18 | A_koj | W 176 34.099 | S 21 59.354 | 1886 |
| **Manus** | Suzette | 139 | 18 | 18 | A_koj | E 152 05.783 | S 03 47.368 | 1505 |
| **Manus** | North_Su | 144 | 13 | 15 | A_koj | E 152 06.046 | S 03 47.933 | 1218 |
| **Manus** | South_Su | 158 | 16 | 16 | A_koj | E 152 06.299 | S 03 48.530 | 1300 |
| **North-Fiji** | Phoenix | 53 | 16 | 16 | A_koj | E 173 55.078 | S 16 56.963 | 1973 |
| **North-Fiji** | Phoenix | 59 | 18 | 22 | A_koj | E 173 55.127 | S 16 57.000 | 1961 |
| **Woodlark** | Scala | 164 | 1 | 3 | A_koj | E 155 03.117 | S 09 47.939 | 3344 |
| **Woodlark** | Scala | 171 | 23 | 23 | A_koj | E 155 03.160 | S 09 47.945 | 3388 |
| **Manus** | Fenway | 102 | 24 | 24 | B_m | E 151 40.360 | S 03 43.675 | 1696 |
| **Manus** | Fenway | 105 | 15 | 16 | B_m | E 151 40.370 | S 03 43.681 | 1698 |
| **Manus** | Solwara_6 | 126 | 13 | 16 | B_m | E 151 40.867 | S 03 43.649 | 1725 |
| **Manus** | Desmos | 127 | 5 | 5 | B_m | E 151 51.933 | S 03 41.528 | 1896 |
| **Manus** | North_Su | 146 | 16 | 16 | B_m | E 152 06.060 | S 03 47.942 | 1210 |
| **Manus** | North_Su | 148 | 16 | 16 | B_m | E 152 06.089 | S 03 47.957 | 1195 |
| **Manus** | South_Su | 160 | 19 | 19 | B_m | E 152 06.300 | S 03 48.482 | 1360 |
| **Manus** | South_Su | 162 | 13 | 16 | B_m | E 152 06.310 | S 03 48.582 | 1353 |

| | | | | | | | | |
|---|---|---|---|---|---|---|---|---|
| **Futuna** | Fati_Ufu | 21 | 2 | 2 | B_m | W 177 11.301 | S 14 45.921 | 1520 |
| **Futuna** | Fati_Ufu | 22 | 5 | 5 | B_m | W 177 11.303 | S 14 45.920 | 1500 |
| **Futuna** | Kulo_Lasi | 36 | 12 | 12 | B_m | W 177 15.004 | S 14 56.537 | 1412 |
| **Futuna** | Kulo_Lasi | 37 | 3 | 3 | B_m | W 177 15.007 | S 14 56.539 | 1414 |
| **Futuna** | Kulo_Lasi | 38 | 1 | 1 | B_m | W 177 15.551 | S 14 56.468 | 1371 |
| **Futuna** | Kulo_Lasi | 41 | 13 | 13 | B_m | W 177 15.565 | S 14 56.384 | 1407 |
| **Lau** | Mangatolo | 45 | 7 | 7 | B_m | W 174 39.209 | S 15 24.878 | 2031 |
| **Manus** | Snowcap | 88 | 22 | 24 | B_m | E 151 40.213 | S 03 43.691 | 1640 |
| **Futuna** | Fati_Ufu | 11 | 3 | 5 | B_seg | W 177 11.082 | S 14 45.587 | 1520 |
| **Manus** | Solwara_7 | 110 | 3 | 3 | B_seg | E 151 40.377 | S 03 43.033 | 1765 |
| **Manus** | Solwara_8 | 114 | 1 | 1 | B_seg | E 151 40.452 | S 03 43.820 | 1737 |
| **Manus** | Romans_Ruins | 118 | 3 | 3 | B_seg | E 151 40.463 | S 03 43.287 | 1666 |
| **Futuna** | Fati_Ufu | 13 | 24 | 30 | B_seg | W 177 11.093 | S 14 45.590 | 1520 |
| **Manus** | Suzette | 137 | 17 | 17 | B_seg | E 152 05.733 | S 03 47.348 | 1497 |
| **Manus** | Suzette | 141 | 2 | 2 | B_seg | E 152 05.794 | S 03 47.345 | 1503 |
| **Woodlark** | Scala | 164 | 3 | 3 | B_seg | E 155 03.117 | S 09 47.939 | 3344 |
| **Woodlark** | Scala | 170 | 17 | 20 | B_seg | E 155 03.160 | S 09 47.935 | 3374 |
| **Manus** | South_Su | 174 | 1 | 3 | B_seg | E 152 06.314 | S 03 48.556 | 1331 |
| **Futuna** | Fati_Ufu | 19 | 2 | 2 | B_seg | W 177 11.149 | S 14 45.576 | 1519 |
| **Futuna** | Fati_Ufu | 20 | 7 | 7 | B_seg | W 177 11.180 | S 14 45.486 | 1527 |
| **Futuna** | Fatu_Kapa | 28 | 5 | 5 | B_seg | W 177 09.257 | S 14 45.165 | 1545 |
| **Futuna** | Fatu_Kapa | 32 | 20 | 20 | B_seg | W 177 09.964 | S 14 44.236 | 1547 |
| **Futuna** | Kulo_Lasi | 39 | 2 | 2 | B_seg | W 177 15.554 | S 14 56.468 | 1371 |
| **Lau** | Mangatolo | 43 | 11 | 18 | B_seg | W 174 39.205 | S 15 24.867 | 2027 |
| **North-Fiji** | Phoenix | 54 | 6 | 9 | B_seg | E 173 55.080 | S 16 56.951 | 1973 |
| **North-Fiji** | Phoenix | 60 | 4 | 6 | B_seg | E 173 55.127 | S 16 57.001 | 1961 |
| **Lau** | Tow_Cam | 67 | 10 | 10 | B_seg | W 176 08.205 | S 20 18.981 | 2716 |
| **Lau** | Tow_Cam | 68 | 9 | 9 | B_seg | W 176 08.207 | S 20 18.982 | 2716 |
| **Lau** | Tow_Cam | 72 | 5 | 6 | B_seg | W 176 08.245 | S 20 19.074 | 2717 |
| **Lau** | Abe | 8 | 11 | 11 | B_seg | W 176 11.526 | S 20 45.762 | 2127 |
| **Lau** | Tui_Malila | 82 | 14 | 23 | B_seg | W 176 34.089 | S 21 59.375 | 1873 |
| **Manus** | Suzette | 87 | 1 | 1 | B_seg | E 152 05.412 | S 03 47.412 | 1579 |
| **Lau** | Abe | 9 | 11 | 11 | B_seg | W 176 11.530 | S 20 45.766 | 2129 |
| **Manus** | Big_Papi | 95 | 17 | 26 | B_seg | E 151 40.334 | S 03 43.729 | 1709 |
| **Manus** | Big_Papi | 96 | 9 | 9 | B_seg | E 151 40.335 | S 03 43.731 | 1708 |
| **Kermadec** | Haungaroa | SH | 20 | 20 | B_seg | W 179°37.12 | S 32°37.264 | NA |
| **Manus** | Solwara_8 | 113 | 24 | 28 | E_ohtai | E 151 40.451 | S 03 43.825 | 1736 |
| **Manus** | North_Su | 143 | 24 | 24 | E_ohtai | E 152 06.045 | S 03 47.936 | 1215 |
| **Futuna** | Fatu_Kapa | 23 | 16 | 17 | E_ohtai | W 177 09.096 | S 14 45.110 | 1567 |
| **Futuna** | Fatu_Kapa | 29 | 7 | 7 | E_ohtai | W 177 09.960 | S 14 44.243 | 1548 |
| **Lau** | Abe | 3 | 24 | 27 | E_ohtai | W 176 11.479 | S 20 45.786 | 2151 |
| **Futuna** | Fatu_Kapa | 31 | 24 | 26 | E_ohtai | W 177 09.961 | S 14 44.243 | 1548 |
| **Futuna** | Kulo_Lasi | 37 | 24 | 26 | E_ohtai | W 177 15.007 | S 14 56.539 | 1414 |

| | | | | | | | | |
|---|---|---|---|---|---|---|---|---|
| **Lau** | Mangatolo | 45 | 24 | 26 | E_ohtai | W 174 39.209 | S 15 24.878 | 2031 |
| **Lau** | Mangatolo | 52 | 24 | 24 | E_ohtai | W 174 39.335 | S 15 24.963 | 2039 |
| **North-Fiji** | Phoenix | 61 | 24 | 29 | E_ohtai | E 173 55.127 | S 16 57.002 | 1961 |
| **Lau** | Tow_Cam | 70 | 24 | 24 | E_ohtai | W 176 08.212 | S 20 19.051 | 2965 |
| **Lau** | Tui_Malila | 80 | 24 | 24 | E_ohtai | W 176 34.077 | S 21 59.280 | 1892 |
| **Futuna** | Fati_Ufu | 16 | 24 | 24 | L_aff_schro | W 177 11.113 | S 14 45.601 | 1519 |
| **Lau** | Abe | 2 | 24 | 26 | L_aff_schro | W 176 11.479 | S 20 45.784 | 2153 |
| **Futuna** | Fatu_Kapa | 26 | 4 | 4 | L_aff_schro | W 177 09.133 | S 14 45.110 | 1562 |
| **Futuna** | Fatu_Kapa | 29 | 20 | 20 | L_aff_schro | W 177 09.960 | S 14 44.243 | 1548 |
| **Futuna** | Kulo_Lasi | 37 | 10 | 12 | L_aff_schro | W 177 15.007 | S 14 56.539 | 1414 |
| **Futuna** | Kulo_Lasi | 38 | 24 | 26 | L_aff_schro | W 177 15.551 | S 14 56.468 | 1371 |
| **Lau** | Mangatolo | 45 | 12 | 12 | L_aff_schro | W 174 39.209 | S 15 24.878 | 2031 |
| **Lau** | Mangatolo | 52 | 12 | 12 | L_aff_schro | W 174 39.335 | S 15 24.963 | 2039 |
| **North-Fiji** | Phoenix | 56 | 18 | 18 | L_aff_schro | E 173 55.111 | S 16 56.936 | 1974 |
| **North-Fiji** | Phoenix | 62 | 18 | 20 | L_aff_schro | E 173 55.133 | S 16 57.005 | 1961 |
| **Lau** | Tow_Cam | 73 | 16 | 24 | L_aff_schro | W 176 08.250 | S 20 19.074 | 2711 |
| **Lau** | Tow_Cam | 76 | 16 | 16 | L_aff_schro | W 176 08.263 | S 20 19.084 | 2711 |
| **Lau** | Tui_Malila | 80 | 16 | 16 | L_aff_schro | W 176 34.077 | S 21 59.280 | 1892 |
| **Lau** | Tui_Malila | 84 | 16 | 16 | L_aff_schro | W 176 34.094 | S 21 59.354 | 1877 |
| **Manus** | Fenway | 103 | 8 | 8 | L_schro | E 151 40.367 | S 03 43.665 | 1699 |
| **Manus** | Fenway | 105 | 8 | 8 | L_schro | E 151 40.370 | S 03 43.681 | 1698 |
| **Manus** | Solwara_7 | 107 | 16 | 18 | L_schro | E 151 40.374 | S 03 43.042 | 1769 |
| **Manus** | Solwara_8 | 111 | 23 | 25 | L_schro | E 151 40.441 | S 03 43.825 | 1739 |
| **Manus** | Solwara_6 | 123 | 15 | 15 | L_schro | E 151 40.854 | S 03 43.654 | 1729 |
| **Manus** | Desmos | 132 | 24 | 44 | L_schro | E 151 51.985 | S 03 41.542 | 1912 |
| **Manus** | South_Su | 135 | 24 | 24 | L_schro | E 152 06.310 | S 03 48.582 | 1353 |
| **Manus** | Suzette | 140 | 15 | 15 | L_schro | E 152 05.783 | S 03 47.368 | 1506 |
| **Manus** | North_Su | 145 | 16 | 16 | L_schro | E 152 06.046 | S 03 47.935 | 1216 |
| **Manus** | North_Su | 146 | 8 | 10 | L_schro | E 152 06.060 | S 03 47.942 | 1210 |
| **Manus** | North_Su | 147 | 12 | 12 | L_schro | E 152 06.084 | S 03 47.957 | 1194 |
| **Manus** | North_Su | 148 | 8 | 16 | L_schro | E 152 06.089 | S 03 47.957 | 1195 |
| **Woodlark** | Scala | 172 | 19 | 31 | L_schro | E 155 03.161 | S 09 47.945 | 3388 |
| **Manus** | Fenway | 98 | 8 | 8 | L_schro | E 151 40.342 | S 03 43.707 | 1703 |
| **Manus** | Fenway | 99 | 6 | 6 | L_schro | E 151 40.344 | S 03 43.704 | 1701 |
| **Manus** | Solwara_1 | CDV | 12 | 12 | L_schro | NA | NA | NA |
| **Manus** | Solwara_8 | CDV | 8 | 8 | L_schro | NA | NA | NA |
| **Kermadec** | Haungaroa | SH | 24 | 24 | L_schro | W 179°37.12 | S 32°37.264 | NA |
| **Lau** | Abe | 2 | 20 | 28 | S_toll | W 176 11.479 | S 20 45.784 | 2153 |
| **Futuna** | Fati_Ufu | 10 | 6 | 18 | S_toll | W 177 10.968 | S 14 45.325 | 1516 |
| **Futuna** | Fati_Ufu | 16 | 6 | 21 | S_toll | W 177 11.113 | S 14 45.601 | 1519 |
| **Futuna** | Fatu_Kapa | 24 | 11 | 13 | S_toll | W 177 09.103 | S 14 45.113 | 1564 |
| **Futuna** | Fatu_Kapa | 29 | 14 | 14 | S_toll | W 177 09.960 | S 14 44.243 | 1548 |
| **Futuna** | Kulo_Lasi | 38 | 23 | 25 | S_toll | W 177 15.551 | S 14 56.468 | 1371 |

| | | | | | | | | |
|---|---|---|---|---|---|---|---|---|
| **Lau** | Mangatolo | 46 | 12 | 12 | S_toll | W 174 39.210 | S 15 24.875 | 2031 |
| **Lau** | Mangatolo | 52 | 12 | 12 | S_toll | W 174 39.335 | S 15 24.963 | 2039 |
| **Fidjien** | Phoenix | 56 | 23 | 25 | S_toll | E 173 55.111 | S 16 56.936 | 1974 |
| **Fidjien** | Phoenix | 62 | 20 | 28 | S_toll | E 173 55.133 | S 16 57.005 | 1961 |
| **Lau** | Tow_Cam | 73 | 12 | 12 | S_toll | W 176 08.250 | S 20 19.074 | 2711 |
| **Lau** | Tow_Cam | 76 | 11 | 13 | S_toll | W 176 08.263 | S 20 19.084 | 2711 |
| **Lau** | Tui_Malila | 80 | 10 | 16 | S_toll | W 176 34.077 | S 21 59.280 | 1892 |
| **Lau** | Tui_Malila | 84 | 12 | 12 | S_toll | W 176 34.094 | S 21 59.354 | 1877 |
| **Manus** | Fenway | 98 | 11 | 17 | S_toll | E 151 40.342 | S 03 43.707 | 1703 |
| **Manus** | Solwara_7 | 107 | 12 | 12 | S_toll | E 151 40.374 | S 03 43.042 | 1769 |
| **Manus** | Solwara_8 | 116 | 12 | 12 | S_toll | E 151 40.460 | S 03 43.820 | 1737 |
| **Manus** | Romans_Ruins | 119 | 3 | 21 | S_toll | E 151 40.469 | S 03 43.287 | 1659 |
| **Manus** | Solwara_6 | 123 | 13 | 15 | S_toll | E 151 40.854 | S 03 43.654 | 1729 |
| **Manus** | South_Su | 135 | 12 | 12 | S_toll | E 152 06.310 | S 03 48.582 | 1353 |
| **Manus** | Suzette | 140 | 14 | 14 | S_toll | E 152 05.783 | S 03 47.368 | 1506 |
| **Manus** | North_Su | 145 | 14 | 14 | S_toll | E 152 06.046 | S 03 47.935 | 1216 |
| **Woodlark** | Scala | 172 | 10 | 10 | S_toll | E 155 03.161 | S 09 47.945 | 3388 |
| **Manus_CDV** | CDV | NA | 41 | 41 | S_toll | 0 | NA | NA |
| **Lau_JUV** | Tui_Malila_JUV | NA | 50 | 52 | S_toll | NA | NA | NA |

*Table 2: Summary table of table S1 per basin, locality and species. The first number represents the number of sites sampled and the second the total number of samples sampled per locality. Species short names: A_koj = Alviniconcha kojimai; B_m = Bathymodiolus manusensis; B_seg = Branchinotogluma segonzaci; E_ohtai = Eochionelasmus ohtai; L_affschro = Lepetodrilus aff. schrolli; L_schro = Lepetodrilus schrolli; S_toll = Shinkailepas tollmann. SUM= SUM for all species of total sites used (ID_sites) and total number of samples used.*

| **Basin** | **Locality** | ***S_toll*** | ***A_koj*** | ***B_m*** | ***B_seg*** | ***E_ohtai*** | ***L_schro & aff schro*** |
|---|---|---|---|---|---|---|---|
| **North-Fiji** | Phoenix | 2 / 53 | 2 / 38 | 0 / 0 | 2 / 15 | 1 / 29 | 2 / 38 |
| **Futuna** | Fati_Ufu | 2 / 39 | 3 / 45 | 2 / 7 | 4 / 44 | 0 / 0 | 1 / 24 |
| **Futuna** | Fatu_Kapa | 2 / 27 | 2 / 18 | 0 / 0 | 2 / 25 | 3 / 50 | 2 / 24 |
| **Futuna** | Kulo_Lasi | 1 / 25 | 0 / 0 | 4 / 29 | 1 / 2 | 1 / 26 | 2 / 38 |
| **Kermadec** | Haungaroa | 0 / 0 | 0 / 0 | 0 / 0 | 1 / 20 | 0 / 0 | 1 / 24 |
| **Lau** | Abe | 1 / 28 | 1 / 2 | 0 / 0 | 2 / 22 | 1 / 27 | 1 / 26 |
| **Lau** | Mangatolo | 2 / 24 | 2 / 21 | 1 / 7 | 1 / 18 | 2 / 50 | 2 / 24 |
| **Lau** | Tow_Cam | 2 / 25 | 2 / 44 | 0 / 0 | 3 / 25 | 1 / 24 | 2 / 40 |
| **Lau** | Tui_Malila | 2 / 28 | 3 / 51 | 0 / 0 | 1 / 23 | 1 / 24 | 2 / 32 |
| **Lau_JUV** | Tui_Malila_JUV | 1 / 52 | 0 / 0 | 0 / 0 | 0 / 0 | 0 / 0 | 0 / 0 |
| **Manus** | Big_Papi | 0 / 0 | 0 / 0 | 0 / 0 | 2 / 35 | 0 / 0 | 0 / 0 |

| | | | | | | | |
|---|---|---|---|---|---|---|---|
| **Manus** | Desmos | 0 / 0 | 0 / 0 | 1 / 5 | 0 / 0 | 0 / 0 | 1 / 44 |
| **Manus** | Fenway | 1 / 17 | 0 / 0 | 2 / 40 | 0 / 0 | 0 / 0 | 4 / 30 |
| **Manus** | North_Su | 1 / 14 | 1 / 15 | 2 / 32 | 0 / 0 | 1 / 24 | 4 / 54 |
| **Manus** | Romans_Ruins | 1 / 21 | 0 / 0 | 0 / 0 | 1 / 3 | 0 / 0 | 0 / 0 |
| **Manus** | Snowcap | 0 / 0 | 0 / 0 | 1 / 24 | 0 / 0 | 0 / 0 | 0 / 0 |
| **Manus** | Solwara_1 | 0 / 0 | 0 / 0 | 0 / 0 | 0 / 0 | 0 / 0 | 1 / 12 |
| **Manus** | Solwara_6 | 1 / 15 | 0 / 0 | 1 / 16 | 0 / 0 | 0 / 0 | 1 / 15 |
| **Manus** | Solwara_7 | 1 / 12 | 0 / 0 | 0 / 0 | 1 / 3 | 0 / 0 | 1 / 18 |
| **Manus** | Solwara_8 | 1 / 12 | 0 / 0 | 0 / 0 | 1 / 1 | 1 / 28 | 2 / 33 |
| **Manus** | South_Su | 1 / 12 | 1 / 16 | 2 / 35 | 1 / 3 | 0 / 0 | 1 / 24 |
| **Manus** | Suzette | 1 / 14 | 1 / 18 | 0 / 0 | 3 / 20 | 0 / 0 | 1 / 15 |
| **Manus_CDV** | CDV | 1 / 41 | 0 / 0 | 0 / 0 | 0 / 0 | 0 / 0 | 0 / 0 |
| **Woodlark** | Scala | 1 / 10 | 2 / 26 | 0 / 0 | 2 / 23 | 0 / 0 | 1 / 31 |
| **SUM** | | 25 / 469 | 20 / 294 | 16 / 195 | 28 / 282 | 12 / 282 | 18 / 546 |

*Table 3: Description of all filtering steps to obtain the VCF files. HWE (step10) test ( performed with snpgdsHWE function from SNPRelate R package* V.1.21.7*), was performed independently within each genetic unit depicted by the PCA and loci were excluded if they deviated from HWE in any populations (genetic cluster) with a p-value threshold of 0.05.*

| **Filtering criteria** | STEP |
|---|---|
| Assembled data | 1 |
| Replicate and samples with more 20 % miss data | 2 |
| SNPs with heterozygosity > 0.6 | 3 |
| max missing data per SNPs 10% | 4 |
| max missing data per sample 10 % (15 % S. tollmanni) | 5 |
| Max Read Depth (DP) > 80 | 7 |
| Minor allele frequency > 0.05 | 8 |
| Max missing data par sample 10 % (15 % S. tollmanni) | 9 |
| Out of Hardy-Weinberg equibrium *p*-value < 0.05 | 10 |
| 1 SNPs per RADtag | 11 |

*Table 4: Filtering steps for each species with their number of variants (SNPs), radtags and samples.*

| | ***A. kojimai*** | | | ***S. tollmanni*** | | |
|---|---|---|---|---|---|---|
| **Filtering step** | **N° SNPs** | **N° radtag** | **N° samples** | **N° SNPs** | **N° radtag** | **N° samples** |
| 1 | 286 975 | 39 931 | 294 | 1 142 237 | 21 424 | 384 |
| 2 | 286 975 | 39 931 | 259 | 1 142 237 | 21 424 | 347 |
| 3 | 286 975 | 39 931 | 259 | 1 142 237 | 21 424 | 347 |
| 4 | 164 171 | 24 375 | 259 | 174 914 | 4 920 | 347 |
| 5 | 164 144 | 24 363 | 259 | 174 914 | 4 920 | 347 |
| 6 | 164 077 | 24 349 | 212 | 174 853 | 4 907 | 356 |
| 7 | 32 031 | 13 728 | 212 | 15 413 | 3 395 | 347 |
| 8 | 32 031 | 13 728 | 212 | 15 413 | 3 395 | 347 |
| 9 | 28 330 | 12 957 | 212 | 11 059 | 3 074 | 347 |
| 10 | 12 957 | 12 957 | 212 | 3 074 | 3 074 | 347 |

| | ***E. ohtai*** | | | ***B. manusensis*** | | |
|---|---|---|---|---|---|---|
| **Filtering step** | **N° SNPs** | **N° radtag** | **N° samples** | **N° SNPs** | **N° radtag** | **N° samples** |
| 1 | 2 144 137 | 103 097 | 282 | 84281 | 11665 | 186 |
| 2 | 2 144 137 | 103 097 | 256 | 84281 | 11665 | 168 |
| 3 | 2 143 451 | 103 077 | 256 | 84281 | 11665 | 168 |
| 4 | 1 407 227 | 74 794 | 256 | 43222 | 6393 | 168 |
| 5 | 1 407 227 | 74 794 | 240 | 43222 | 6393 | 161 |
| 6 | 1 407 110 | 74 773 | 240 | 43222 | 6393 | 161 |
| 7 | 118 173 | 48 486 | 240 | 6783 | 3356 | 161 |
| 8 | 118 173 | 48 486 | 224 | 6783 | 3356 | 159 |
| 9 | 104 892 | 45 984 | 224 | 5358 | 2904 | 159 |
| 10 | 45 984 | 45 984 | 224 | 2904 | 2904 | 159 |

| | ***B. segonzaci*** | | | ***L. schrolli & L. aff schrolli*** | | |
|---|---|---|---|---|---|---|
| **Filtering step** | **N° SNPs** | **N° radtag** | **N° samples** | **N° SNPs** | **N° radtag** | **N° samples** |
| 1 | 899 488 | 93 006 | 262 | 1 463 969 | 35 025 | 522 |
| 2 | 899 488 | 93 006 | 233 | 1 463 969 | 35 025 | 451 |
| 3 | 899 488 | 93 006 | 233 | 1 463 969 | 35 025 | 451 |
| 4 | 640691 | 69106 | 233 | 1 161 506 | 27 853 | 451 |
| 5 | 640691 | 69106 | 233 | 1 161 506 | 27 853 | 423 |
| 6 | 640673 | 69101 | 233 | 1 107 329 | 26 695 | 423 |
| 7 | 147128 | 50221 | 233 | 87 258 | 22 147 | 423 |
| 8 | 147128 | 50221 | 204 | 87 258 | 22 147 | 423 |
| 9 | 127888 | 47547 | 204 | 48 979 | 18 187 | 423 |
| 10 | 47547 | 47547 | 204 | 18 187 | 18 187 | 423 |

*NB : In addition, a filter specific to L. schrolli (& L. affschro) was applied before the last step 10, to remove radtags and variants potentially linked to some genomic architecture or sex determism. To achieve this, markers contributing more than 50% to the PC1 of the separation in Manus BAB were removed from the total dataset.*

*Table 5: Stacks parameters for de novo assembly of bi-allelic loci between individuals and number of SNPs and individuals for each dataset at 1 SNP per ddRADtag/loci. n.SNPs : number of SNPs structure analysis; n.SNPs dadi dataset : number of SNPs dadi analysis; n.Ind: total number of individuals. (in bold data from Tran Lu Y et al. 2022). m is the Minimum stack depth, M the Distance allowed between stacks and n the Distance allowed between catalog loci.*

| **Species** | ***m*** | ***M*** | ***n*** | ***n.SNPs*** | ***n.SNPs dadi dataset*** | ***n.Ind*** |
|---|---|---|---|---|---|---|
| ***Ifremeria nautilei*** | **5** | **6** | **6** | **10 570** | **17 365** | **362** |
| *Alviniconcha kojimai* | 6 | 7 | 7 | 12 957 | 39 669 | 212 |
| *Shinkailepas tollmanni* | 5 | 11 | 11 | 3 074 | 12 953 | 347 |
| *Eochionelasmus ohtai* | 4 | 7 | 7 | 45 984 | 74 773 | 224 |
| *Bathymodiolus manusensis* | 5 | 5 | 5 | 2 904 | 6 340 | 159 |
| *Branchinotogluma segonzaci* | 4 | 4 | 4 | 47 547 | 69 101 | 195 |
| *Lepetodrilus schrolli & L. affschro* | 4 | 5 | 5 | 18 187 | 26 627 | 414 |

*Table 6: Pairwise $F_{st}$ matrix between BABs for each species calculated with arlequin (V.3.5.2.2). Significativity threshold for 10000 permutations (*:0.05, **:0.01, ***:<0.001). For the species L. schrolli/aff. schrolli and B. segonzaci, one additional population has been sampled and processed from the Kermadec BAB (display in red). For coherence between species and readability, these results from Kermadec have not been reported and discussed in the main part of the paper*

| *A. kojimai* | | | | | |
|---|---|---|---|---|---|
| | Manus | Woodlark | North Fiji | Lau | Futuna |
| Manus | 0.00000 | | | | |
| Woodlark | -0.00079 | 0.00000 | | | |
| North Fiji | **0.01651*** | **0.01668*** | 0.00000 | | |
| Lau | **0.01888*** | **0.01911*** | **0.00317*** | 0.00000 | |
| Futuna | **0.01868*** | **0.01881*** | **0.00324*** | -0.00044 | 0.00000 |

| *E. ohtai* | | | | |
|---|---|---|---|---|
| | Manus | Lau | North Fiji | Futuna |
| Manus | 0.00000 | | | |
| Lau | **0.20507*** | 0.00000 | | |
| North Fiji | **0.21365*** | **-0.00008** | 0.00000 | |
| Futuna | **0.20765*** | -0.00022 | **0.00008** | 0.00000 |

| *B. manusensis* | | | |
|---|---|---|---|
| | Manus | Lau | Futuna |
| Manus | 0.00000 | | |
| Lau | **0.19262*** | 0.00000 | |
| Futuna | **0.20663*** | -0.00315 | 0.00000 |

| *S. tollmanni* | | | | | | |
|---|---|---|---|---|---|---|
| | Manus | Woodlark (M) | Woodlark (NFFL) | North Fiji | Lau | Futuna |
| Manus | 0.00000 | | | | | |
| Woodlark_M | **0.00695** | 0.00000 | | | | |
| Woodlark_NFFL | **0.26753*** | **0.21692** | 0.00000 | | | |
| North Fiji | **0.27114*** | **0.23451*** | 0.00146* | 0.00000 | | |
| Lau | **0.27280*** | **0.23649*** | -0.00248 | -0.00103 | 0.00000 | |
| Futuna | **0.27256*** | **0.23492*** | -0.00212 | -0.00049 | -0.00057 | 0.00000 |

| *B. segonzaci* | | | | | | |
|---|---|---|---|---|---|---|
| | Manus | Woodlark | North Fiji | Futuna | Lau | Kermadec |
| Manus | 0.00000 | | | | | |
| Woodlark | **0.00921*** ** | 0.00000 | | | | |
| North Fiji | **0.04145*** ** | **0.03551*** ** | 0.00000 | | | |
| Futuna | **0.04132*** ** | **0.03658*** ** | **0.00099*** ** | 0.00000 | | |
| Lau | **0.04087*** ** | **0.03640*** ** | **0.00076*** ** | **-0.00021* ** | 0.00000 | |
| Kermadec | 0.03695*** | 0.03071*** | -0.00318 | -0.00334 | -0.00332 | 0.00000 |

| *L. schrolli & L. aff. schrolli* | | | | | | |
|---|---|---|---|---|---|---|
| | Manus | Woodlark | Lau | Futuna | North Fiji | Kermadec |
| Manus | 0.00000 | | | | | |
| Woodlark | **0.18480*** ** | 0.00000 | | | | |
| Lau | **0.36353*** ** | **0.15474*** ** | 0.00000 | | | |
| Futuna | **0.36170*** ** | **0.15610*** ** | **0.00992*** ** | 0.00000 | | |
| North Fiji | **0.35485*** ** | **0.14153*** ** | **0.02855*** ** | **0.02557*** ** | 0.00000 | |
| Kermadec | **0.41578*** ** | **0.30053*** ** | **0.18267*** ** | **0.18120*** ** | 0.27850** | 0.00000 |

*Table 7: All dadi parameters for the best model and each species. species are by column and row is the parameter value under the best model. all parameters are scaled by Nref which is scale by theta. Nu1 and Nu2 represent population size. b1 and b2 represent the growth factor over time. hrf represents the hill-robertson factor. Ts represents the divergence time without migration for AM, SC and SI and with migration for IM model. Tsc represents the time of secondary contact between the two populations. m12, represents the unrestricted migration rate from population 2 towards population 1; me12, the restricted migration rate (e.g., barrier loci) from population 2 towards population 1. Q represents the proportion of loci affected by the linked selection (hill-robertson effect) and P the proportion of loci that show unconstrained gene flow (m12 and m21).*

| | ***A. kojimai*** | ***S. tollmanni*** | ***S. tollmanni*** | ***E. ohtai*** | ***B. segonzaci*** | ***B. manusensis*** | ***B. manusensis*** | ***B. manusensis*** | ***L. schrolli & L. aff schrolli*** |
|---|---|---|---|---|---|---|---|---|---|
| Model | SC2N2mG | SC2N2m | SC2N2mG | SC2mG | SC2N2mG | IM2N | SC2N2m | SC2N | SC2N2mG |
| **Nu1 (NFFL)** | 0.694 | 16.057 | 1.752 | 1.343 | 2.757 | 10.896 | 6.052 | 9.524 | 1.077 |
| **Nu2 (M/W)** | 2.750 | 1.527 | 3.568 | 1.203 | 0.605 | 10.691 | 5.282 | 10.809 | 1.018 |
| **b1** | 57.949 | | 4.378 | 31.559 | 2.191 | | | | 5.581 |
| **b2** | 7.981 | | 0.524 | 4.080 | 4.082 | | | | 39.899 |
| **hrf** | 0.381 | 0.042 | 0.039 | 0.656 | 0.064 | 0.075 | 0.013 | 0.080 | 0.089 |
| **Ts** | 0.627 | 1.037 | 1.004 | 1.000 | 0.467 | 2.156 | 1.486 | 1.216 | 1.041 |
| **Tsc** | 0.252 | 0.089 | 0.103 | | 0.394 | | 0.105 | 0.818 | 1.470 |
| **m12** | 4.741 | 0.321 | 0.445 | 0.364 | 0.579 | 0.436 | 3.989 | 0.520 | 0.878 |
| **m21** | 4.389 | 3.314 | 2.887 | 1.386 | 6.489 | 0.566 | 4.891 | 0.537 | 0.308 |
| **me12** | 0.384 | 4.334 | 4.747 | 0.013 | 2.195 | | 0.227 | | 0.009 |
| **me21** | 0.841 | 13.866 | 14.381 | 0.124 | 5.412 | | 0.646 | | 0.007 |
| **P** | 0.851 | 0.014 | 0.039 | 0.508 | 0.000 | | 0.323 | | 0.180 |
| **Q** | 0.005 | 0.434 | 0.416 | | 0.232 | 0.580 | 0.699 | 0.531 | 0.569 |
| **Theta** | 860.782 | 107.227 | 109.056 | 1 268.530 | 1 714.163 | 140.211 | 81.495 | 137.803 | 136.458 |
| **Nref** | 23 243.21 | 30 529.04 | 31 049.58 | 28 403.58 | 19 491.165 | 23 594.133 | 13 713.680 | 23 188.985 | 23 243.207 |

*NB : in main text table 3, for S. tollmanni and B. segonzaci, the purpose of understanding and displaying the main gene flow value, m and me have been switch, as well for P (switch to 1-P).*

*Table S8: all Standard deviation (SD) parameters for the best model and each species, estimated with Fisher Information Matrix (FIM) from dadi and from SI Table 7 value.*

| | ***A. kojimai*** | ***S. tollmanni*** | ***S. tollmanni*** | ***E. ohtai*** | ***B. segonzaci*** | ***B. manusensis*** | ***B. manusensis*** | ***B. manusensis*** | ***L. schrolli & L. aff schrolli*** |
|---|---|---|---|---|---|---|---|---|---|
| | SC2N2mG | SC2N2m | SC2N2mG | SC2mG | SC2N2mG | IM2N | SC2N2m | SC2N | SC2N2mG |
| **Nu1 (NFFL)** | 0.091 | 3.519 | 3.493 | 0.160 | 1.755 | 2.792 | 0.855 | 1.957 | 0.527 |
| **Nu2 (M/W)** | 0.181 | 0.322 | 0.342 | 0.157 | 0.287 | 3.253 | 1.115 | 2.881 | 0.054 |
| **b1** | 15.152 | | 0.380 | 4.124 | 4.401 | | | | 13.586 |
| **b2** | 1.707 | | 2.349 | 0.438 | 2.324 | | | | 0.526 |
| **hrf** | 1.883 | 0.009 | 0.007 | | 0.024 | 0.008 | 0.006 | 0.009 | 0.018 |
| **Ts** | 0.073 | 0.135 | 0.122 | 0.103 | 0.533 | 0.481 | 0.239 | 0.302 | 0.410 |
| **Tsc** | 0.016 | 0.022 | 0.074 | 0.051 | 0.064 | | 0.047 | 0.202 | 0.085 |
| **m12** | 0.502 | 0.113 | 0.090 | 0.057 | 0.376 | 0.130 | 1.911 | 0.105 | 0.197 |
| **m21** | 0.540 | 0.973 | 0.227 | 0.142 | 5.379 | 0.171 | 2.419 | 0.134 | 0.112 |
| **me12** | 0.082 | 1.492 | 3.020 | 0.002 | 0.858 | | 0.156 | | 0.004 |
| **me21** | 0.011 | 4.009 | 3.968 | 0.011 | 4.475 | | 0.308 | | 0.008 |
| **P** | 0.014 | 0.147 | 0.086 | 0.013 | 0.530 | | 0.085 | | 0.123 |
| **Q** | 0.078 | 0.090 | 0.068 | | 0.129 | 0.042 | 0.071 | 0.044 | 0.253 |
| **Theta** | 35.588 | 7.896 | 8.253 | 71.235 | 341.493 | 21.664 | 8.309 | 17.119 | 23.705 |

### Figures

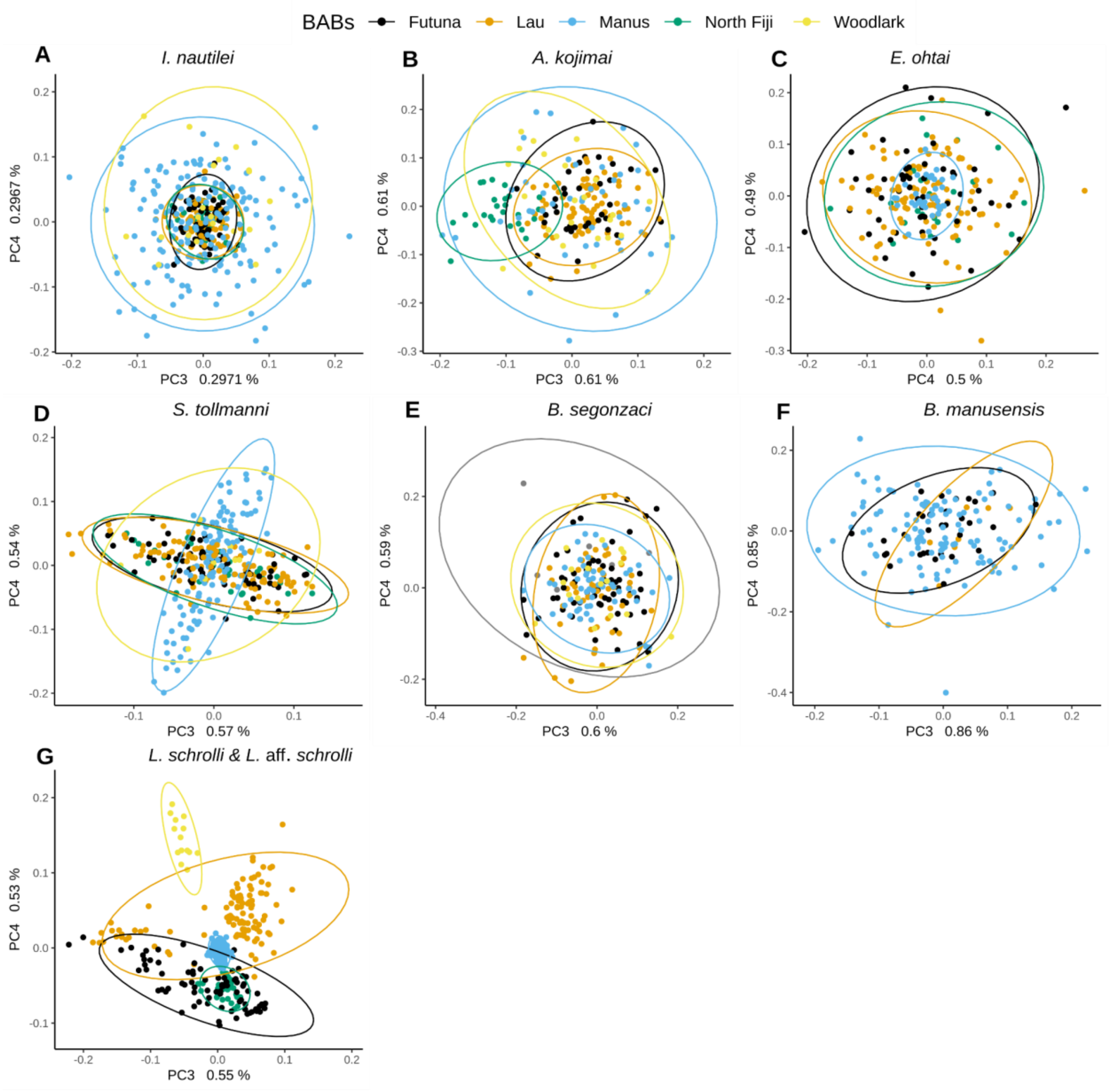


*Figure 1: All PCA plots for Principal Component 3 and 4 for each species. Open ellipses represent the multivariate normal distribution of each group at 95%, here BABs. I. nautilei data are directly taken from Tran Lu Y et al., 2022*

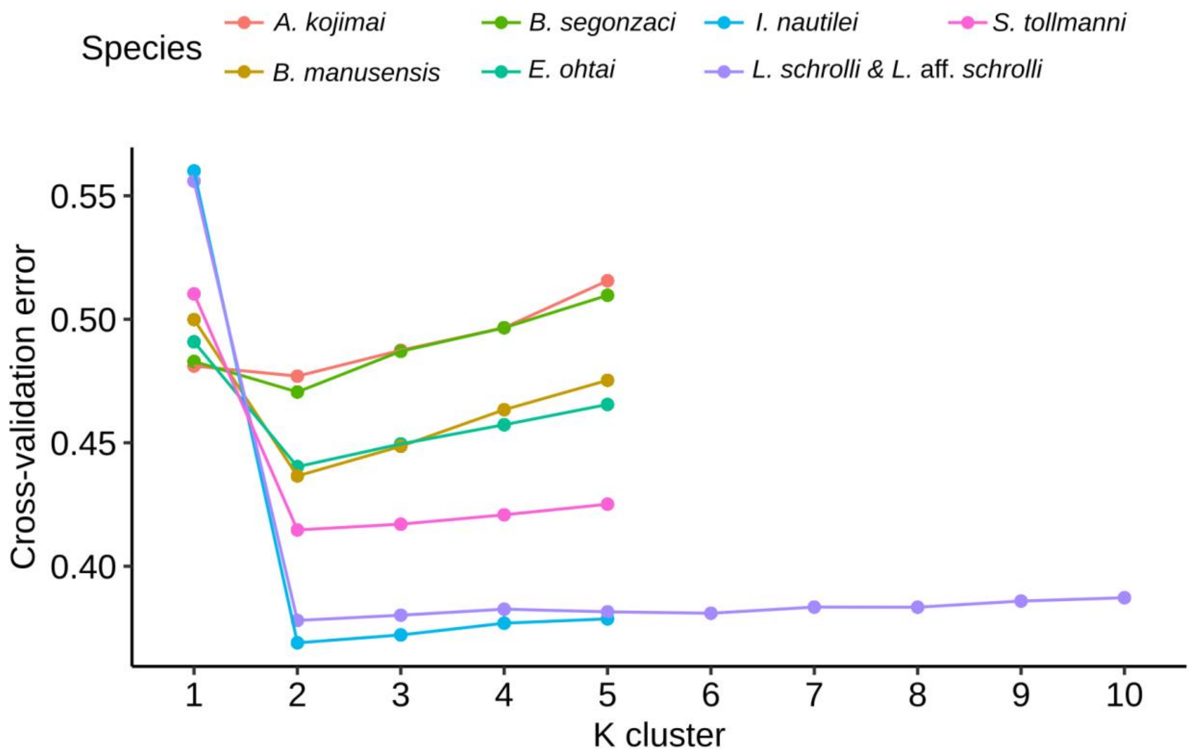


*Figure 2 : Cross-validation error for each K value for all species and runs of Admixture analysis. Color represents species. I. nautilei data are directly taken from Tran Lu Y et al., 2022*

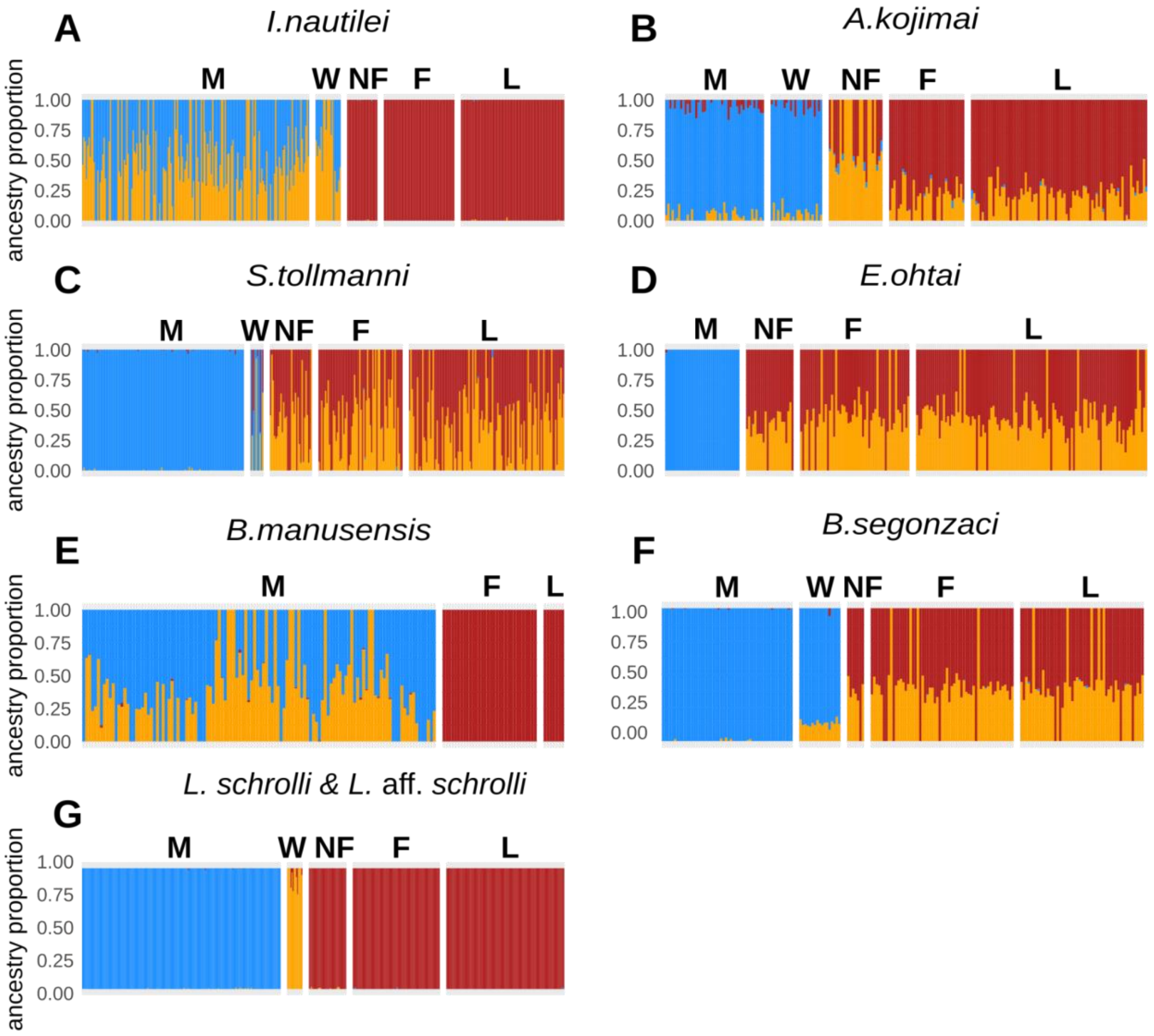


*Figure 3 : Bar Plot of K=3 Admxiture analysis for each species. Each color represents a genetic cluster and each bar represents an individual. All individuals are grouped by geographic zone. I. nautilei data are directly taken from Tran Lu Y et al., 2022*

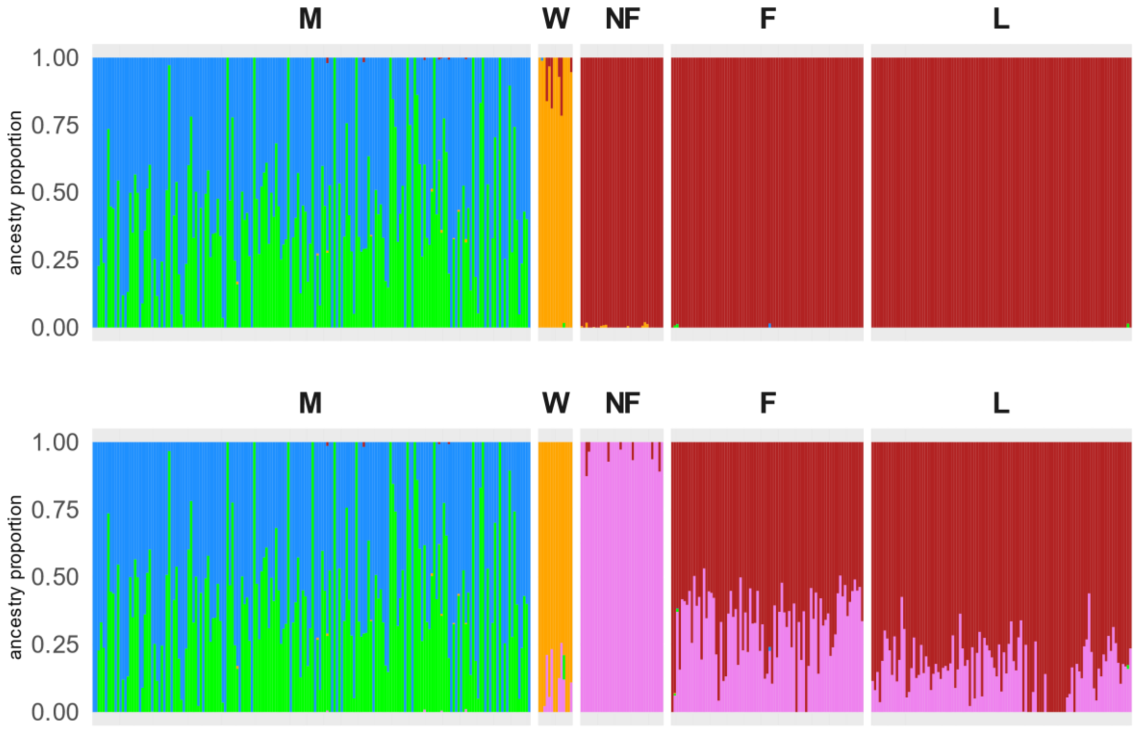


*Figure 4 : Barplot of K=4 and K=5 Admture analysis for L. schrolli & aff schrolli. Each color represents a genetic cluster and each bar represents an individual. All individuals are grouped by geographic zone.*

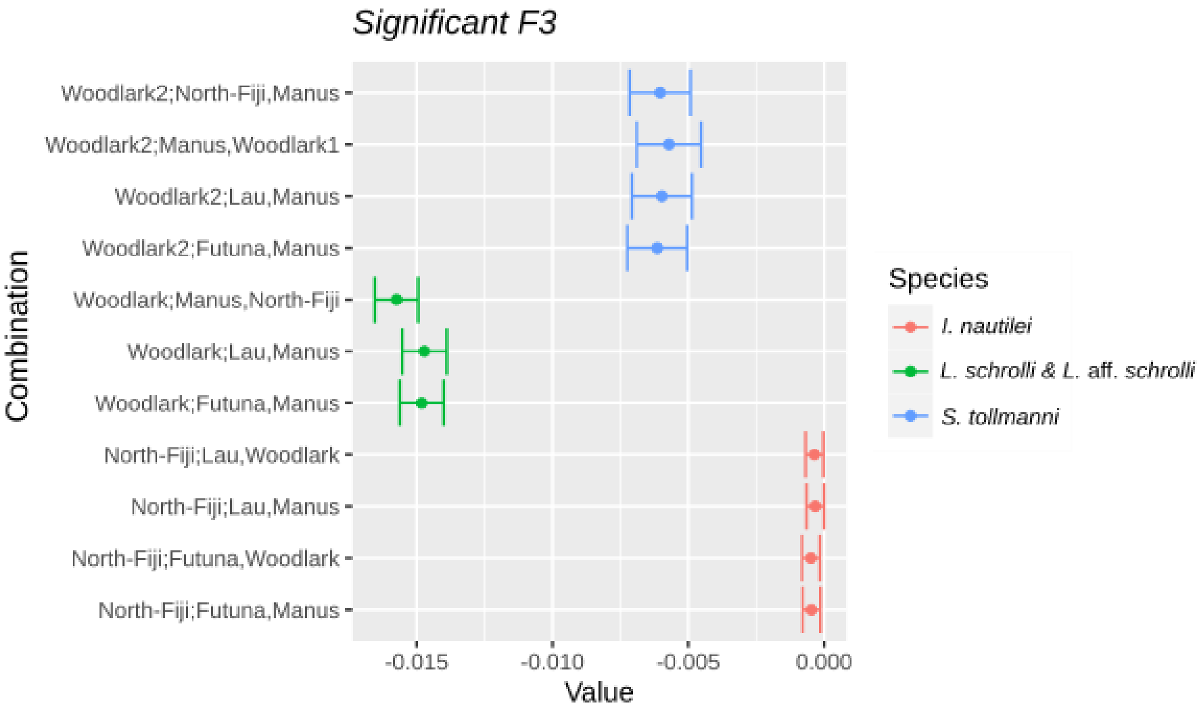


*Figure 5 : Plot of f3 statistics and the confidence interval at 95%. Combinaison represent, first the focal population and then the two source populations. I. nautilei data are directly taken from Tran Lu Y et al., 2022*

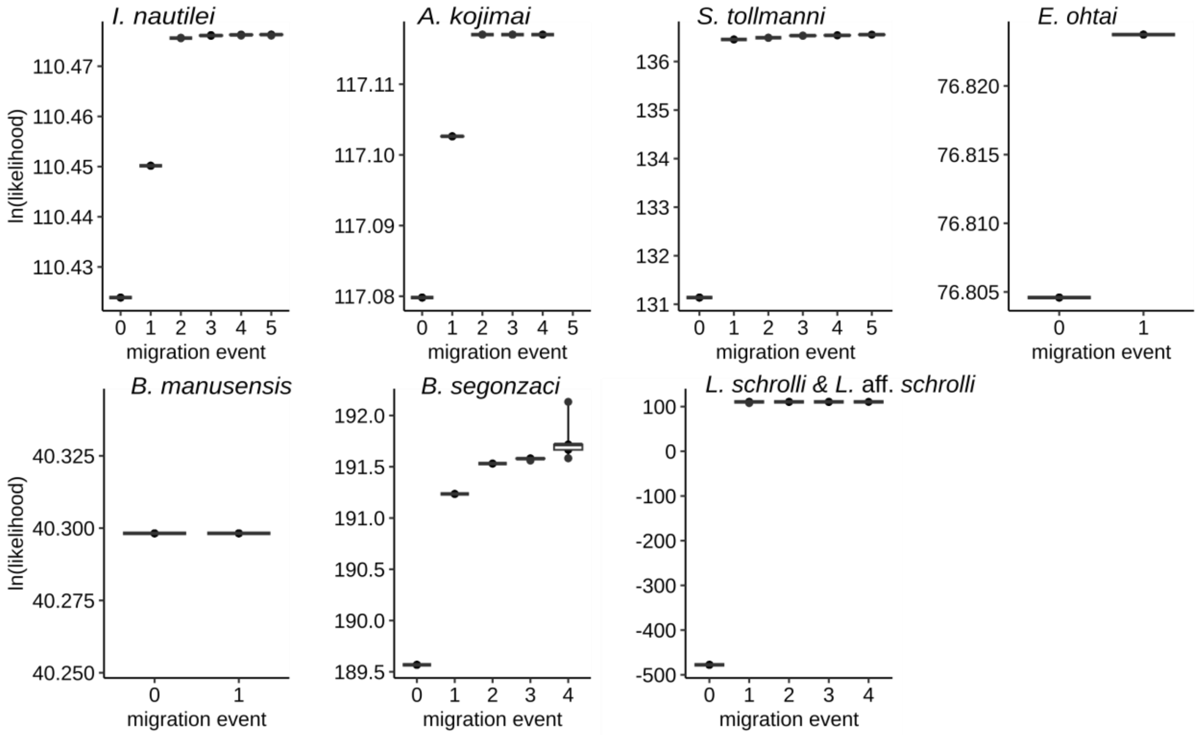


*Figure 6 : Log likelihood for each species and each migration event for 10 independent runs . I. nautilei data are directly taken from Tran Lu Y et al., 2022*

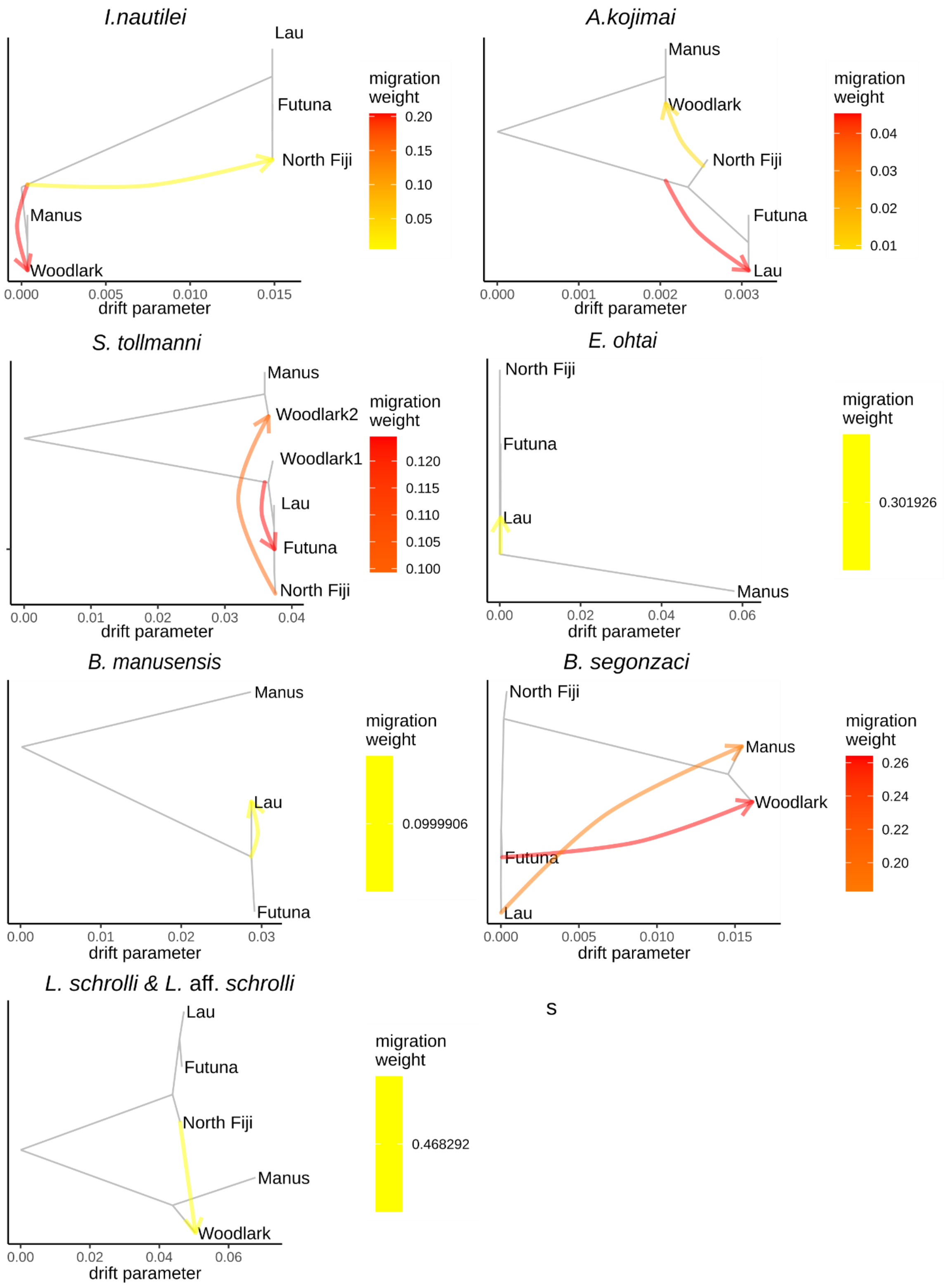


*Figure 7 : Treemix plot for each species and the optimal number of migration events. I. nautilei data are directly taken from Tran Lu Y et al., 2022*

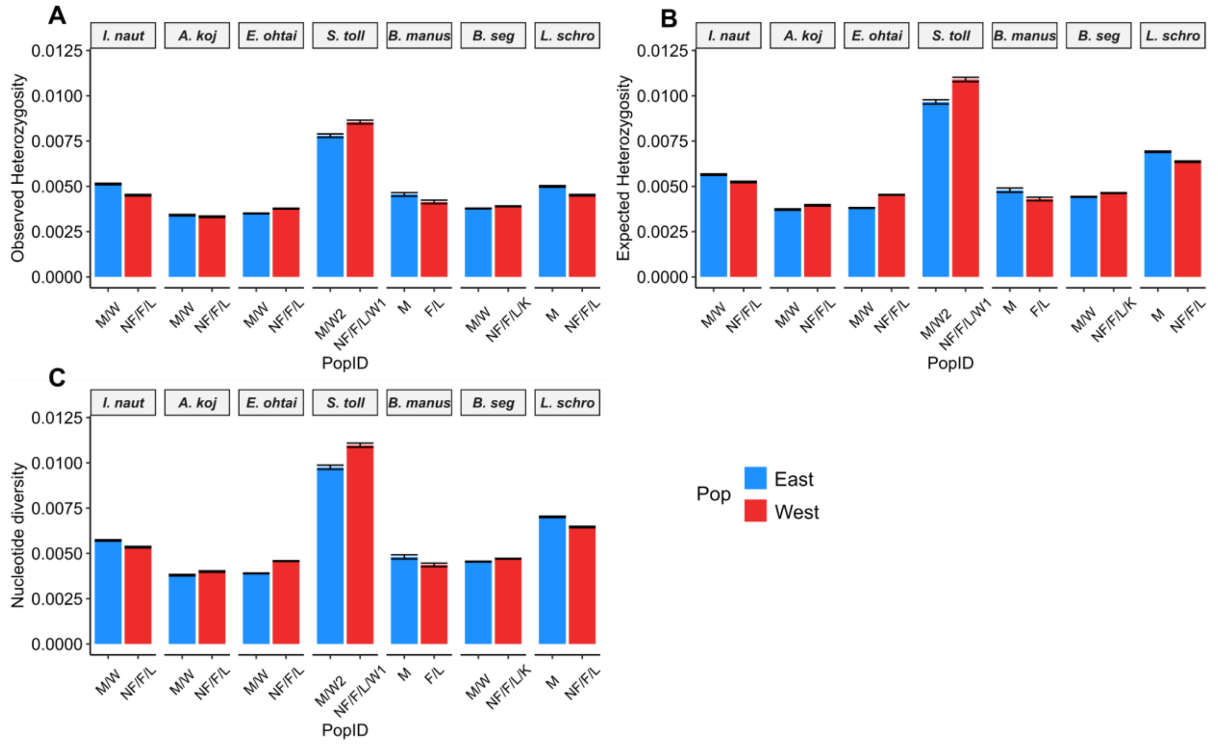


*Figure 8: Observed (Ho) and expected (He) heterozygosities and nucleotide diversity (π) calculated with Stacks2 on the final dataset including all sites (variant and non variant). Error bars represent the confidence interval (CI) estimated at 95%. Colors identify the two main metapopulations (lineages). I. nautilei data are directly taken from Tran Lu Y et al., 2022. I. naut : I. nautilei, A. koj: A. kojimai, S. toll: S. tollmanni; B. manus:B. manusensis, B. seg: B. segonzaci, L. schro: L. schrolli & aff schrolli*

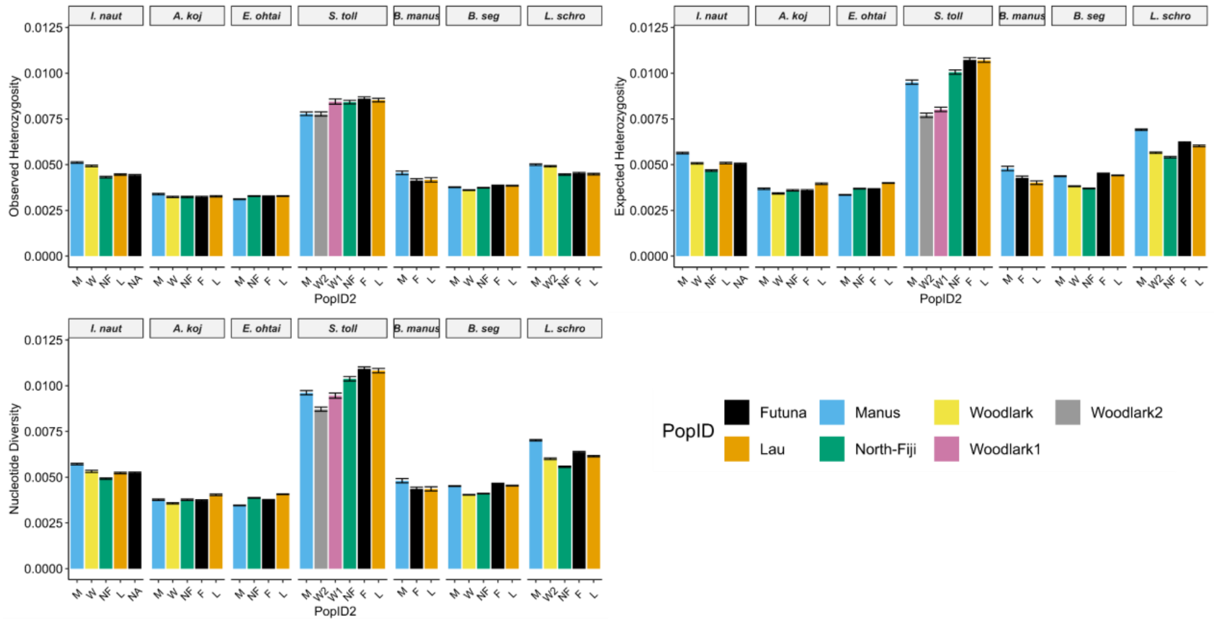


*Figure 9: Observed (Ho) and expected (He) heterozygosities and nucleotide diversity (π) calculated with Stacks2 on the final dataset including all sites (variant and non variant). Error bars represent the confidence interval (CI) estimated at 95%. Colors*

*identify each BAB. I. nautilei data are directly taken from Tran Lu Y et al., 2022. I. nautilei, A. koj: A. kojimai, S. toll: S. tollmanni; B. manus:B. manusensis, B. seg: B. segonzaci, L. schro: L. schrolli & aff schrolli*

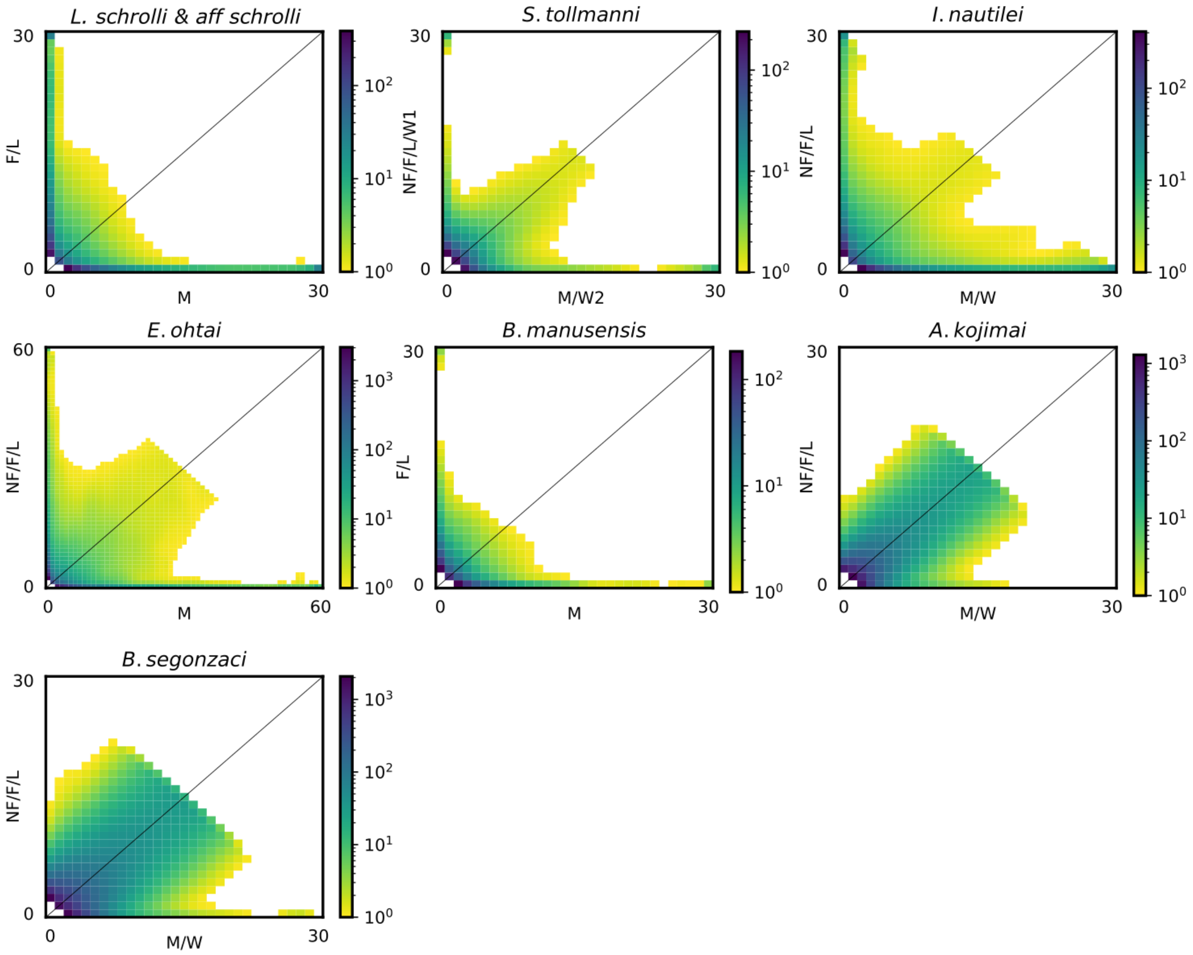


*Figure 10:. For each plot Observed Joint allele frequency spectrum (JAFS) between lineages East and West per species. Color gradients display the frequency of each SNPs class between populations. Population abbreviation for NF: North Fiji, F: Futuna, L: Lau, M: Manus, W: Woodlark.*

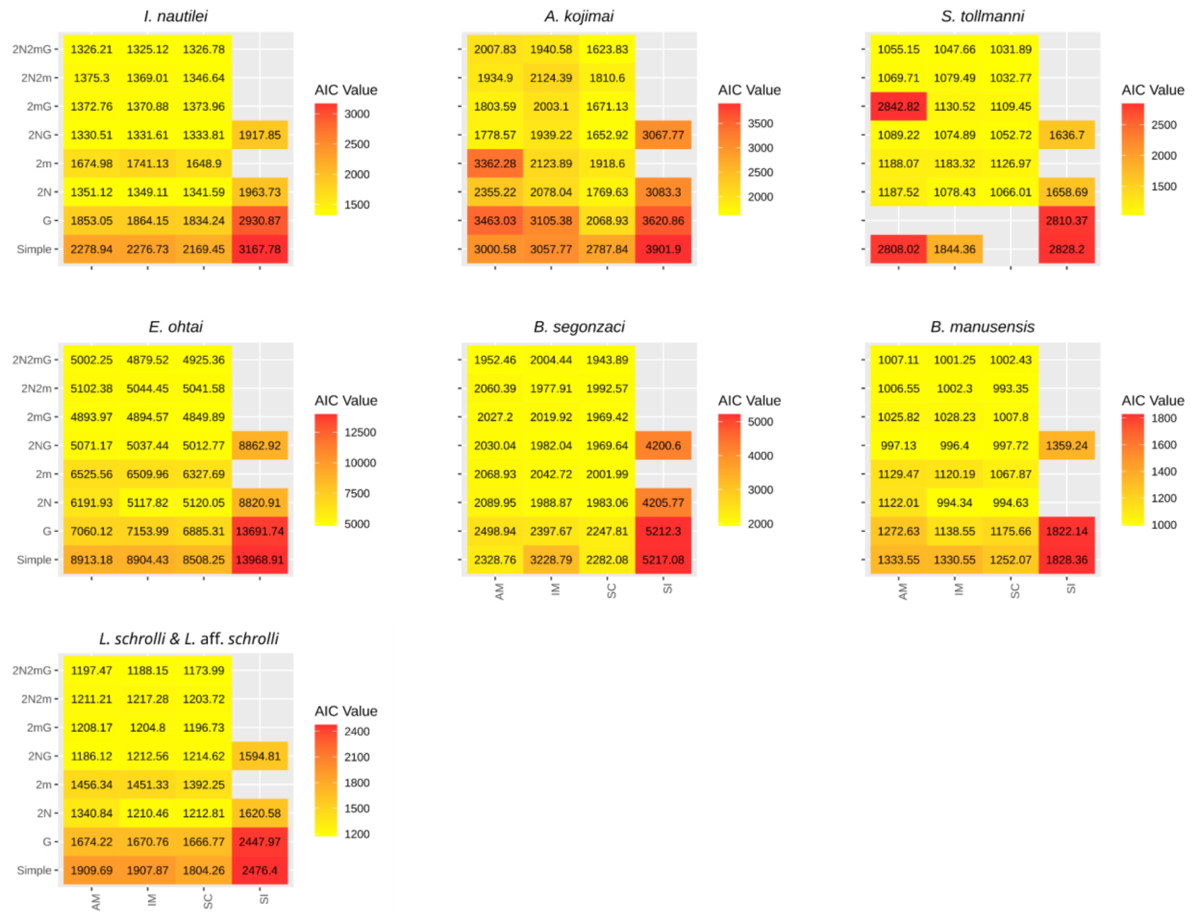


*Figure 11 : Heatmap of AIC values on the best runs for all models and parameters per species. The lowest value represents the best model fit. Ifremeria nautilei data from Tran Lu Y et al. (2022).*

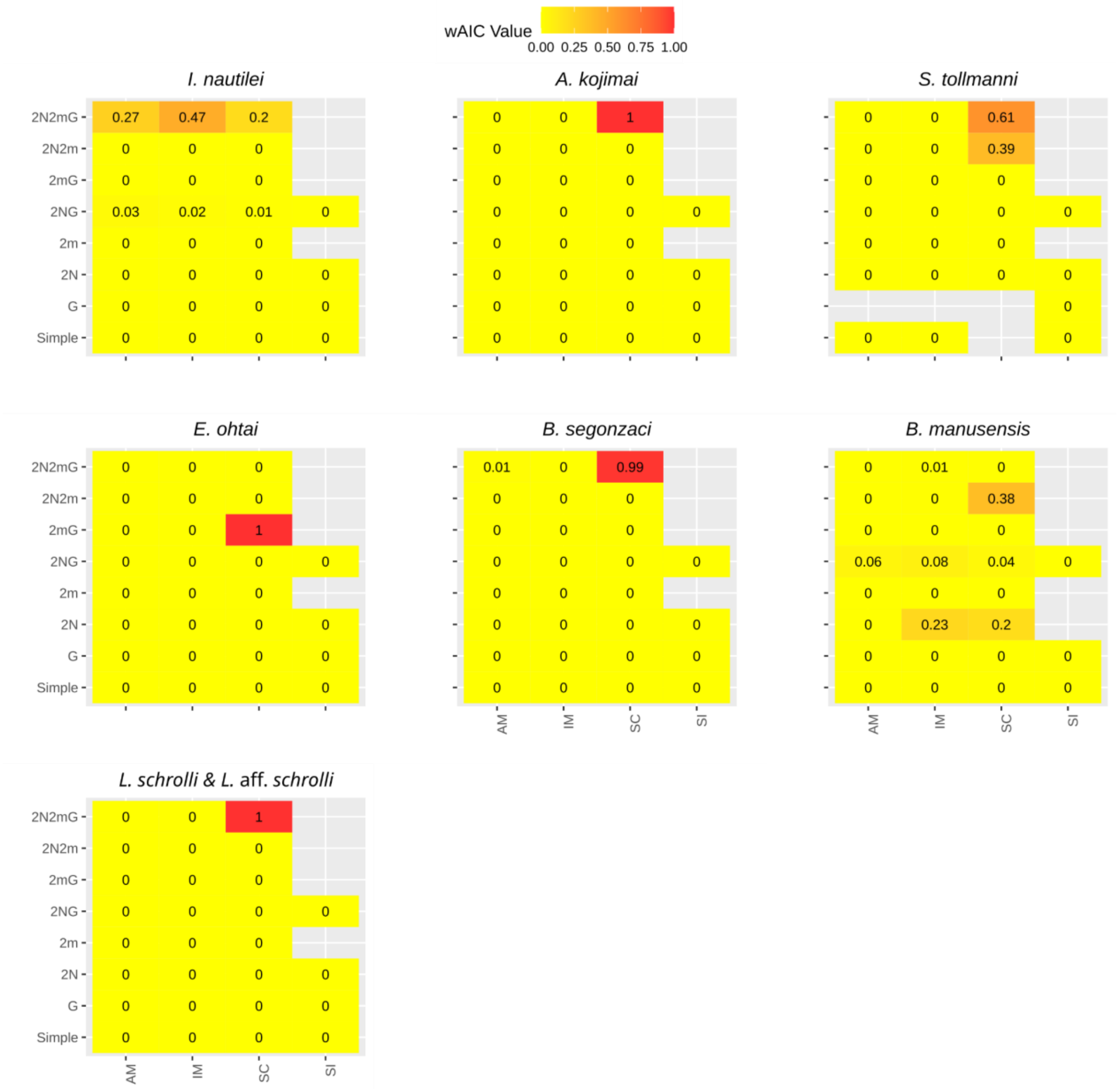


*Figure 12 : Weighted AIC value on the best runs for all models and parameters per species. Highest value represents the best model fit (the lowest AIC value). Ifremeria nautilei data are derived from Tran Lu Y et al. (2022).*

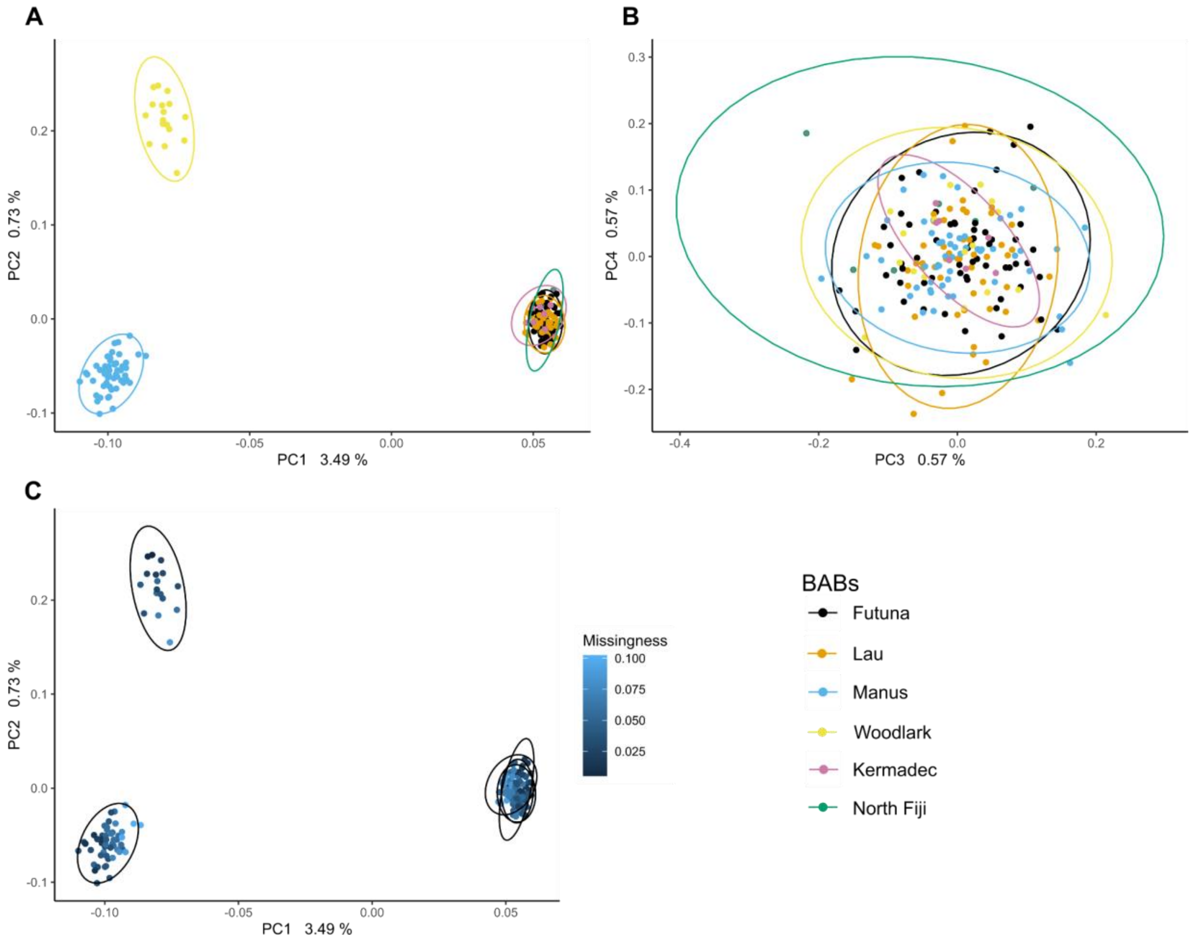


*Figures 13 : PCA plots for B. segonzaci with Kermadec individuals (in black) for first four components and missingness data per individual.*

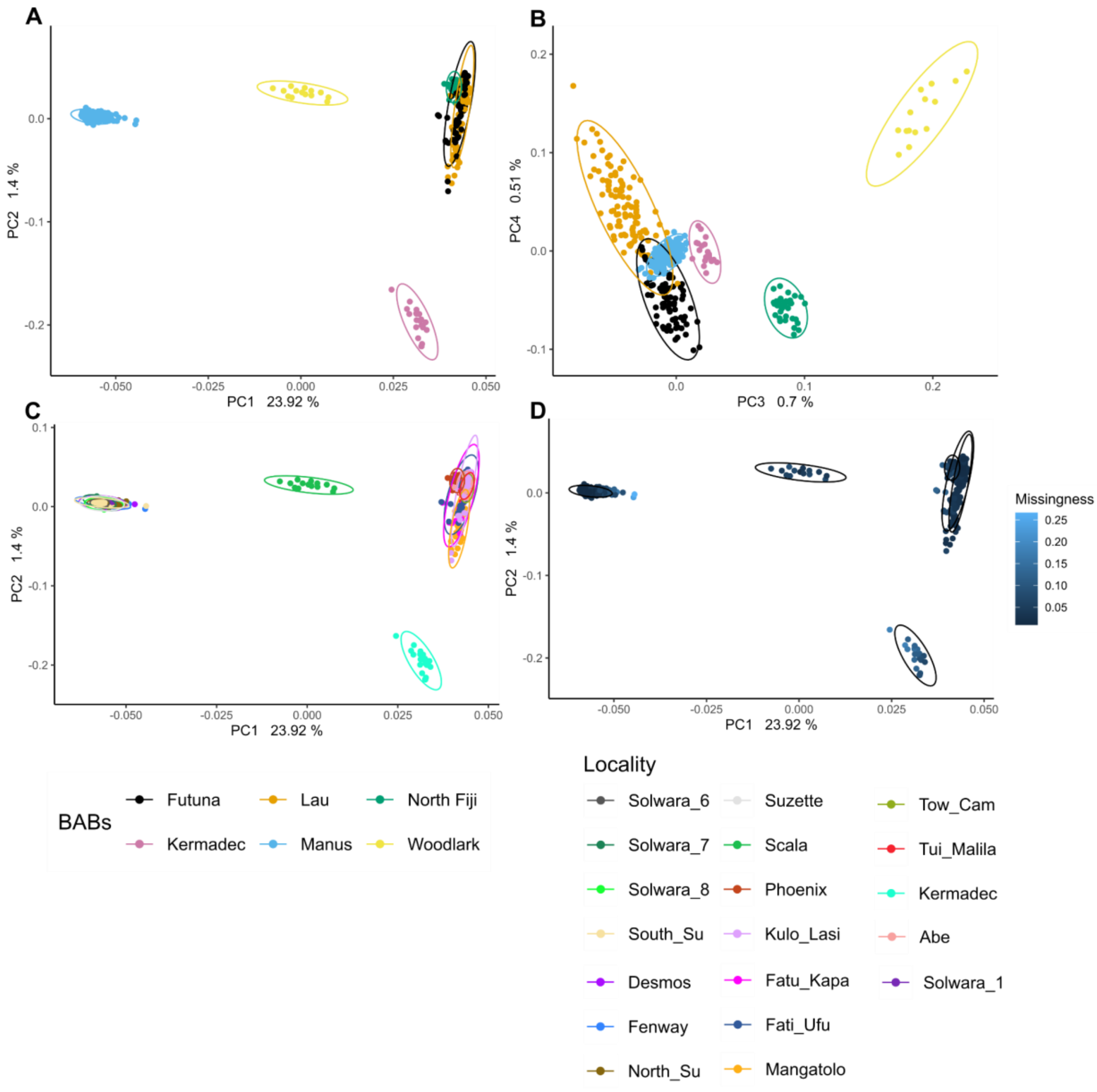


*Figures 14 : PCA plots for L. schrolli & aff schrolli, with Kermadec individuals (in pink) for first four components and missingness data per individual. in A, B colors represents basin. in C colors represents localities and D the pourcent of missing data per sample.*

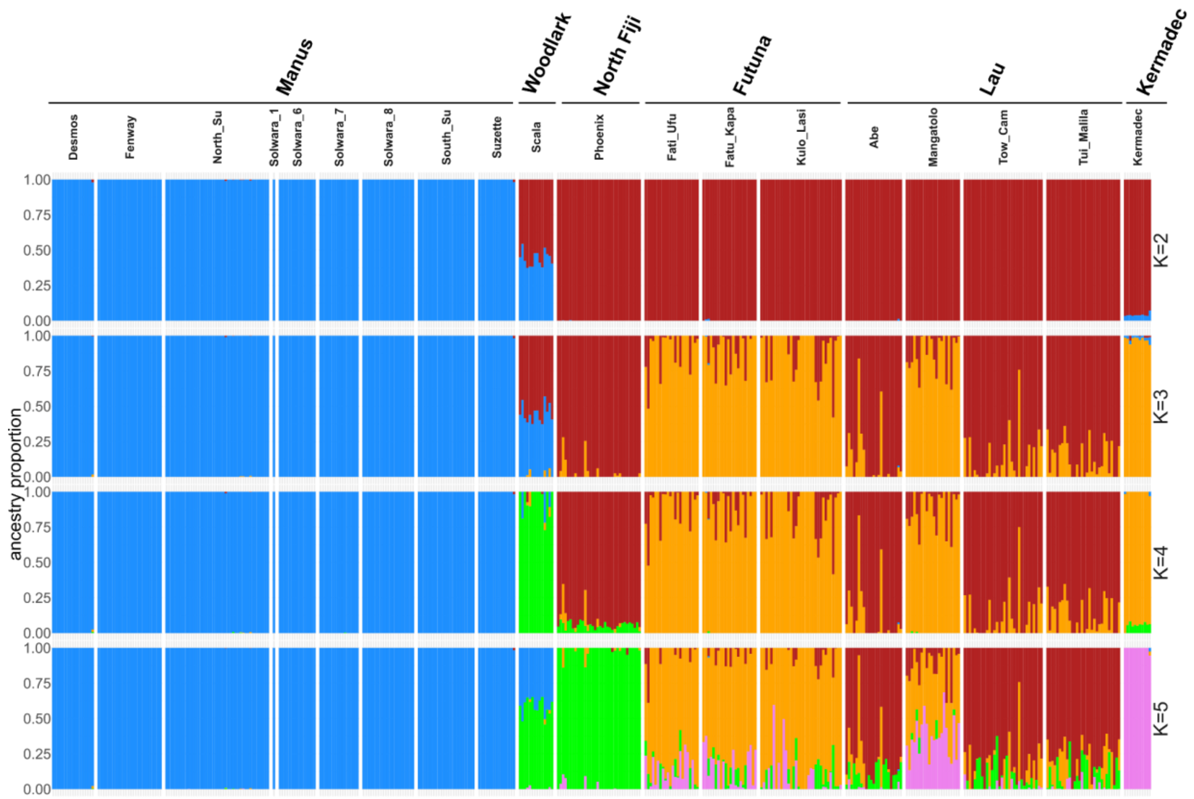


*Figures 15 : Barplot of K=2 to K=5 of Admxiture analysis for L. schrolli &L.* aff. *schrolli* ***Including Kermadec samples.*** *Each color represents a genetic cluster and each bar represents an individual. All individuals are grouped by geographic zone.*

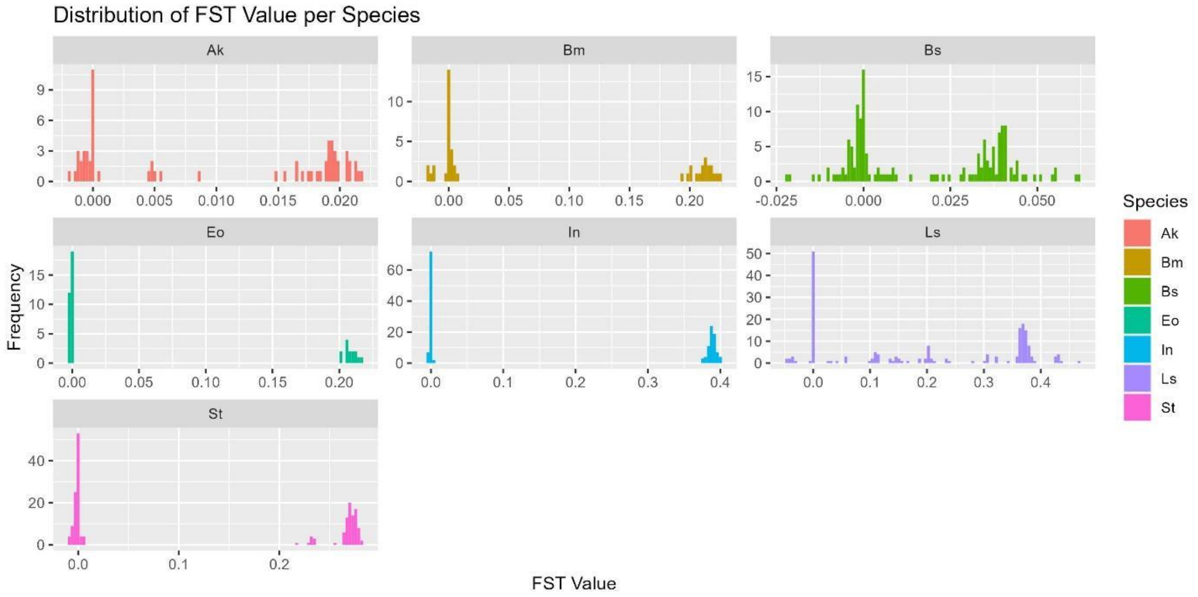


*Figure 16: Histogram of pairwise Fst between all localities. Colors represent species. Ak: A. kojimai, Bm: B. manusensis, Bs: B. segonzaci. Eo: E. ohtai, In: I. nautilei, Ls: L. schrolli & L.* aff. *schrolli. St: S. tollmanni*

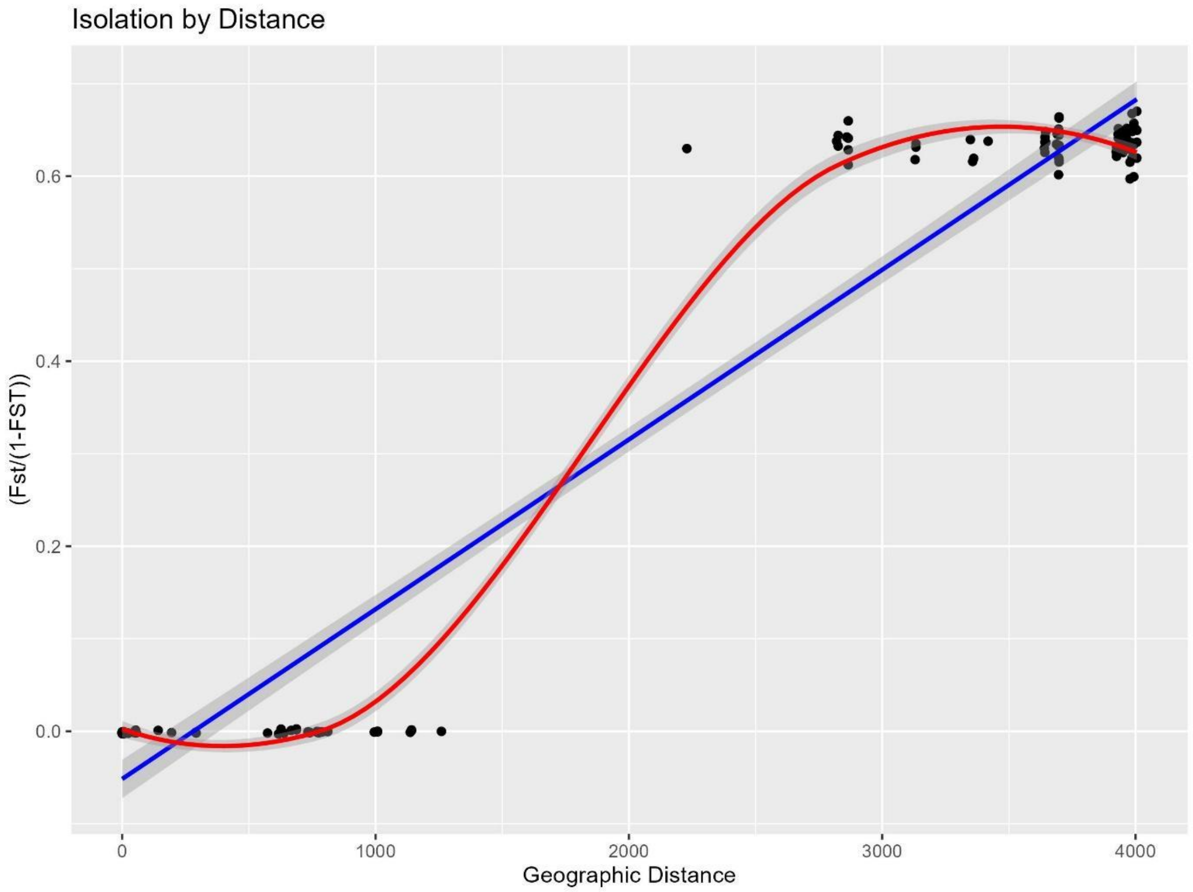


Figure 17: Relationship between genetic differentiation measured as Linearized Fst and geographic distance in km for ***Ifremeria nautilei***. Each point represents a pairwise comparison between localities. Blue line is the model of linear regression (Lm) and the red line the model of polynomial regression (Loess). Grey band represents standart deviations (se) of the model at 95%.

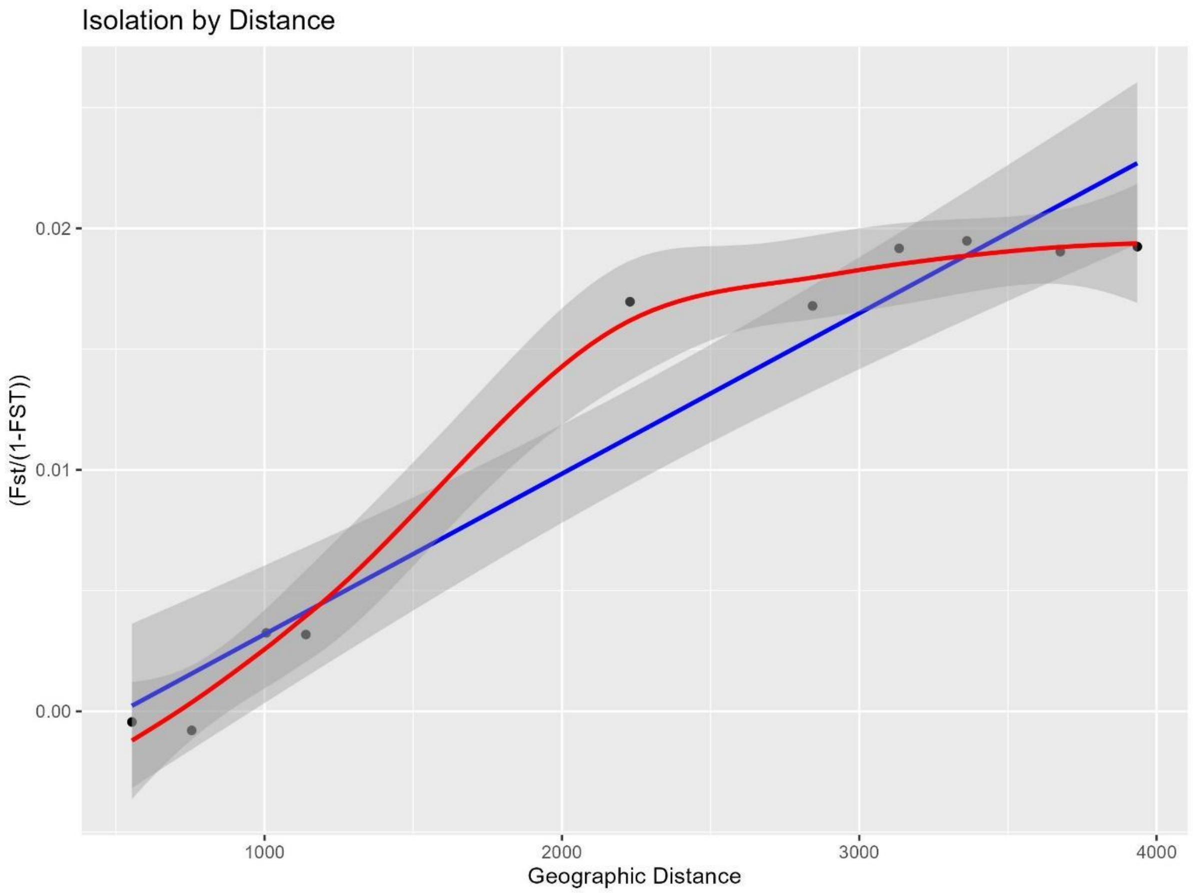


Figure 18: Relationship between genetic differentiation measured Linearized Fst and geographic distance in Km for ***Alviniconcha kojimai***. Each point represents a pairwise comparison between localities. Blue line is the model of linear regression (Lm) and the red line the model of polynomial regression (Loess). Grey band represents of standart deviations (se) of the model at 95%.

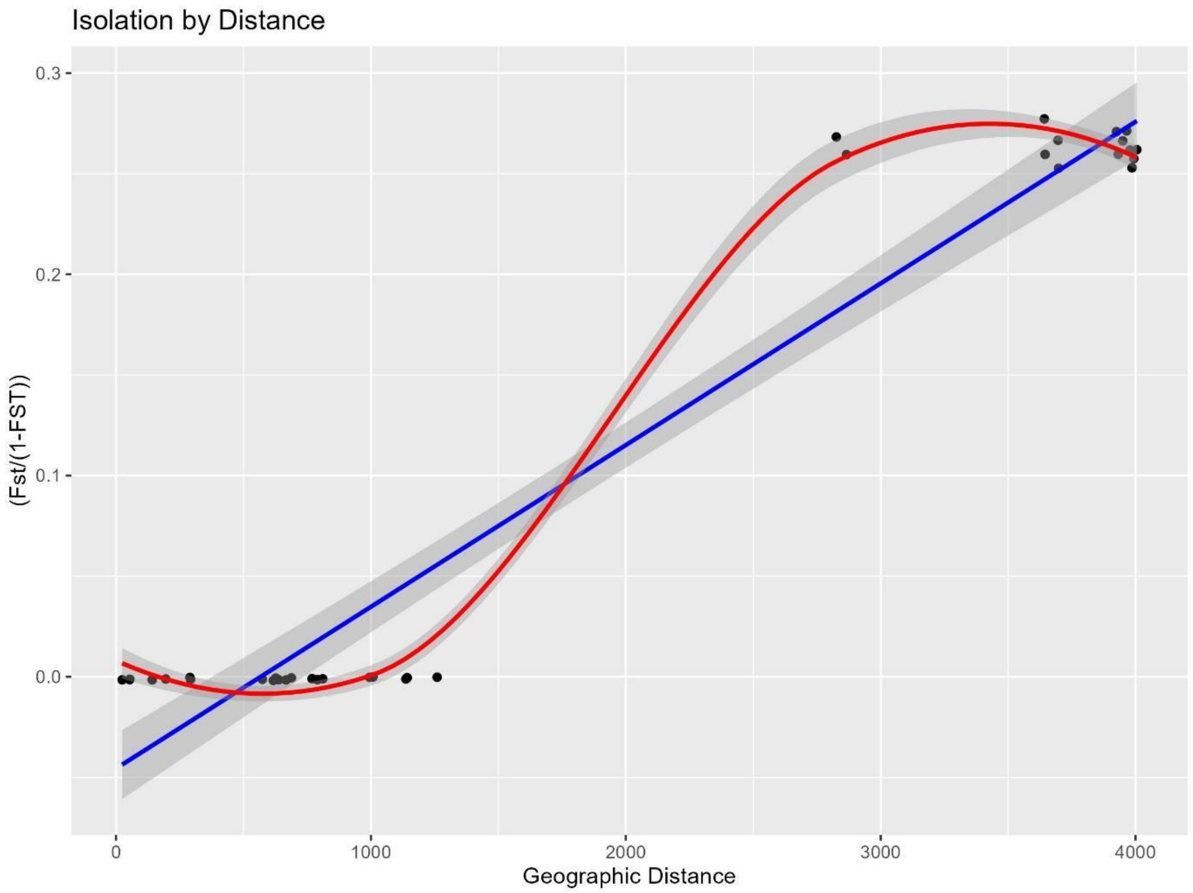


Figure 19: Relationship between genetic differentiation measured as Linearized Fst and geographic distance in km for ***Eochionelasmus ohtai***. Each point represents a pairwise comparison between localities. Blue line is the model of linear regression (Lm) and the red line the model of polynomial regression (Loess). Grey band represents of standart deviations (se) of the model at 95%.

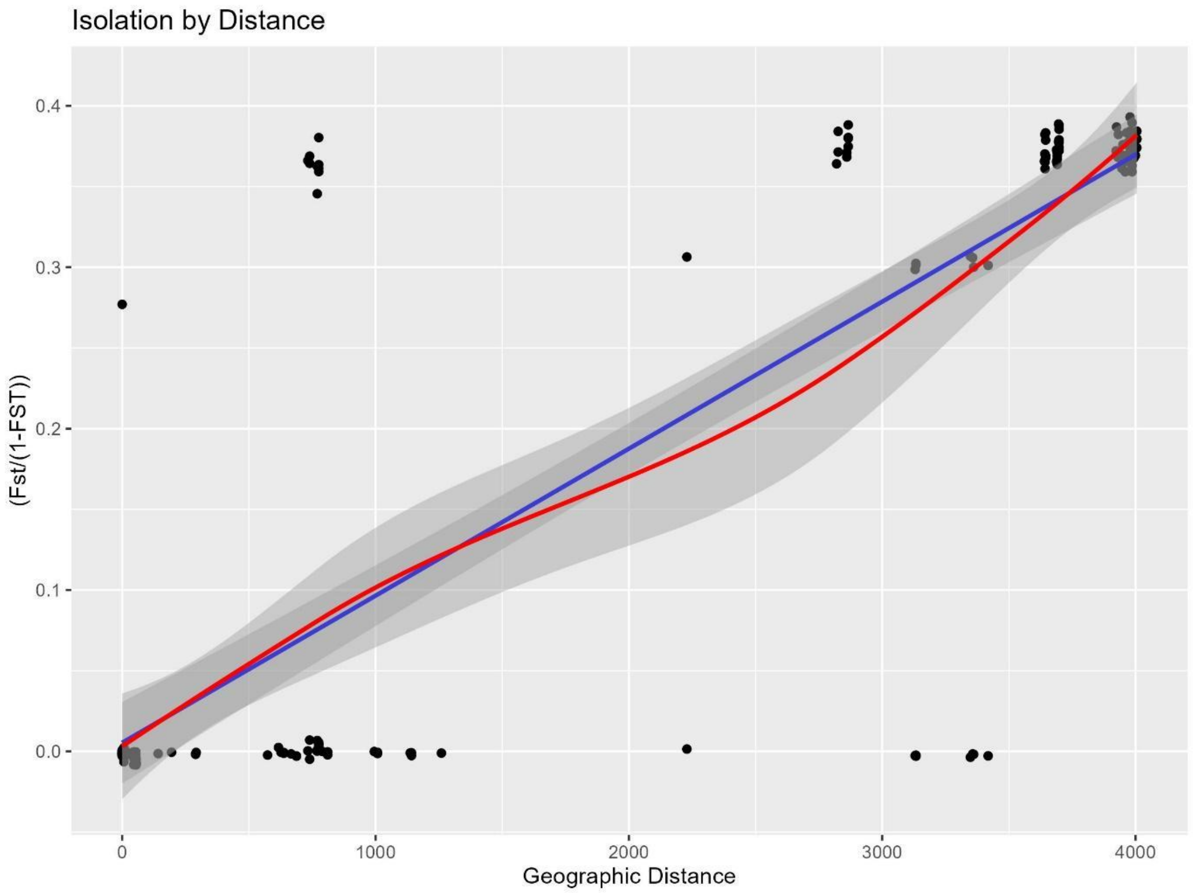


Figure 20: Relationship between genetic differentiation measured as Linearized Fst and geographic distance in km for ***S. tollmanni***. Each point represents a pairwise comparison between localities. Blue line is the model of linear regression (Lm) and the red line the model of polynomial regression (Loess). Grey band represents of standart deviations (se) of the model at 95%.

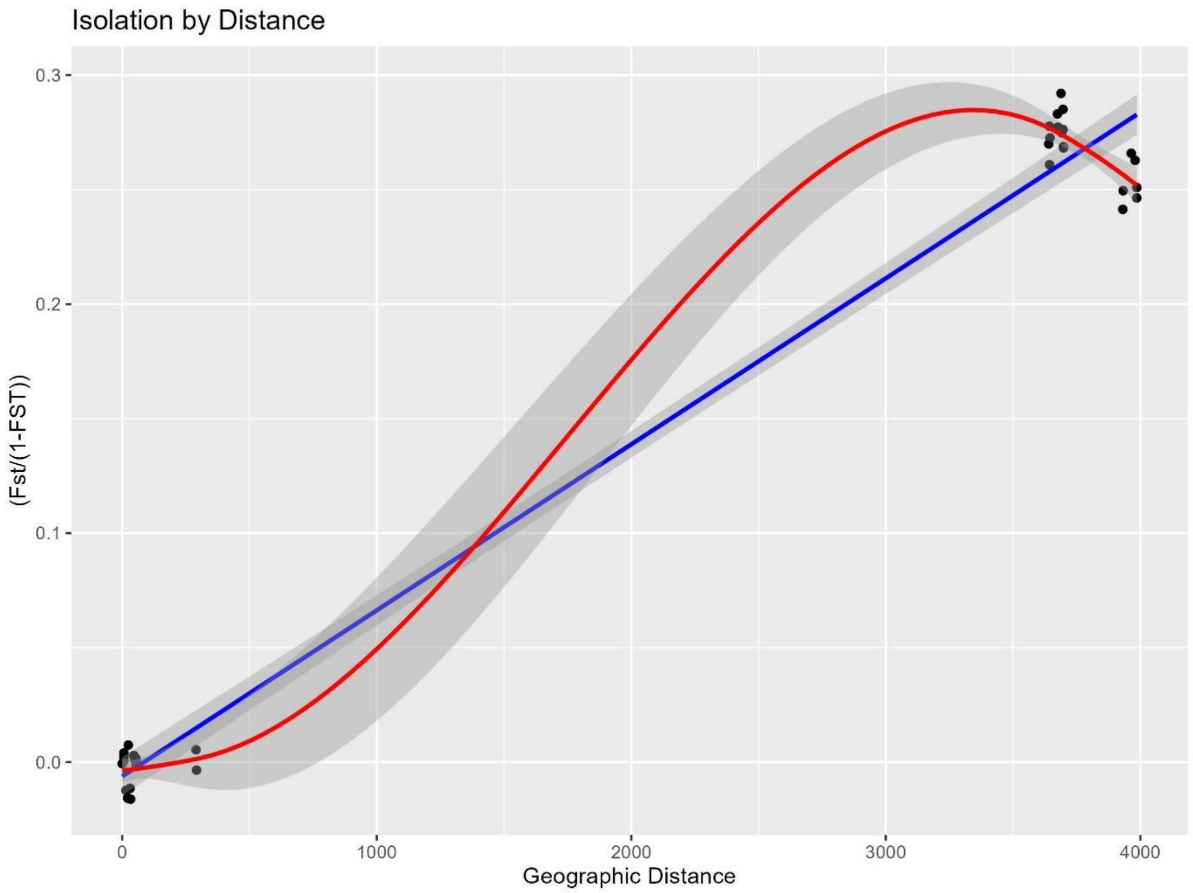


Figure 21: Relationship between genetic differentiation measured as Linearized Fst and geographic distance in km for ***Bathymodiolus manusensis***. Each point represents a pairwise comparison between localities. Blue line is the model of linear regression (Lm) and the red line the model of polynomial regression (Loess). Grey band represents of standart deviations (se) of the model at 95%.

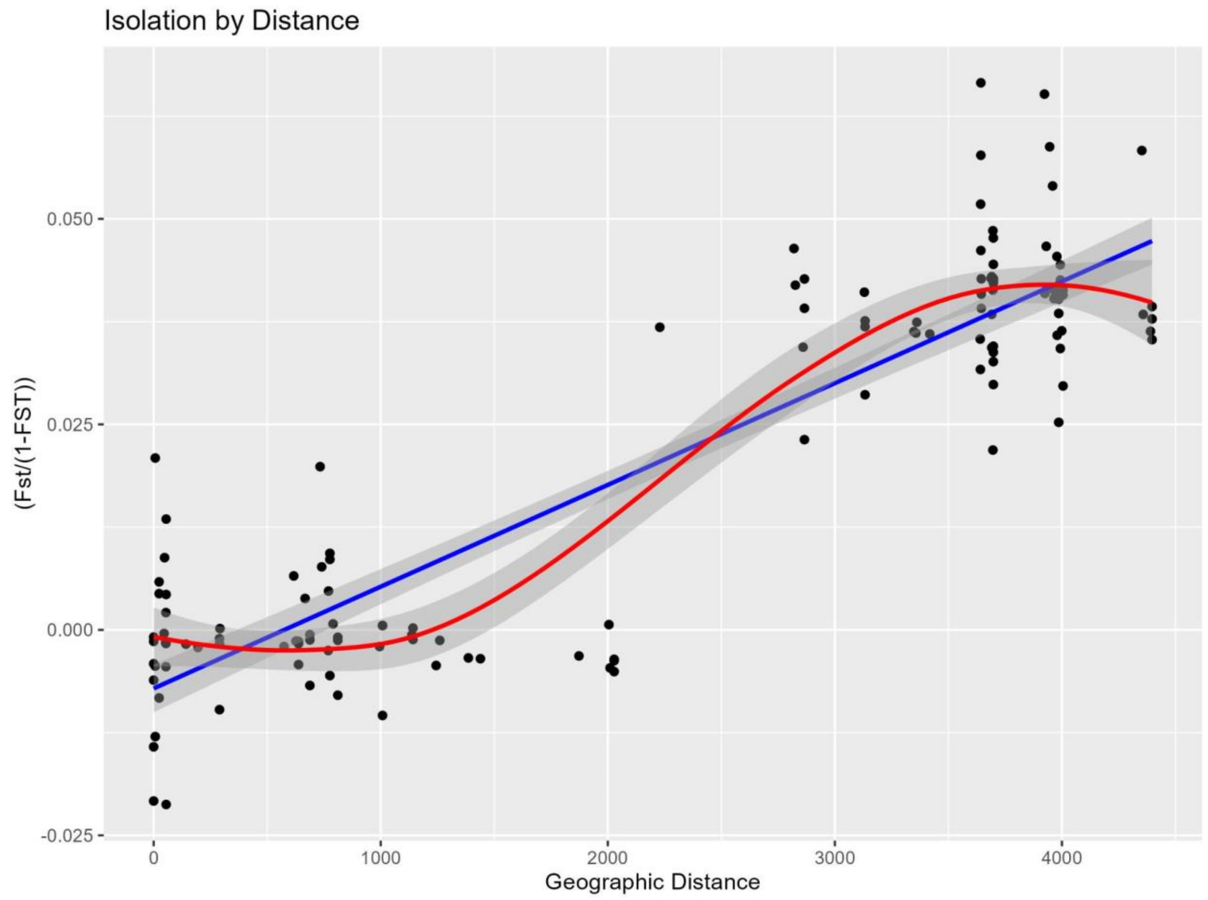


Figure 22: Relationship between genetic differentiation measured as Linearized Fst and geographic distance in km for ***Branchinotogluma segonzaci***. Each point represents a pairwise comparison between localities. Blue line is the model of linear regression (Lm) and the red line the model of polynomial regression (Loess). Grey band represents of standart deviations (se) of the model at 95%.

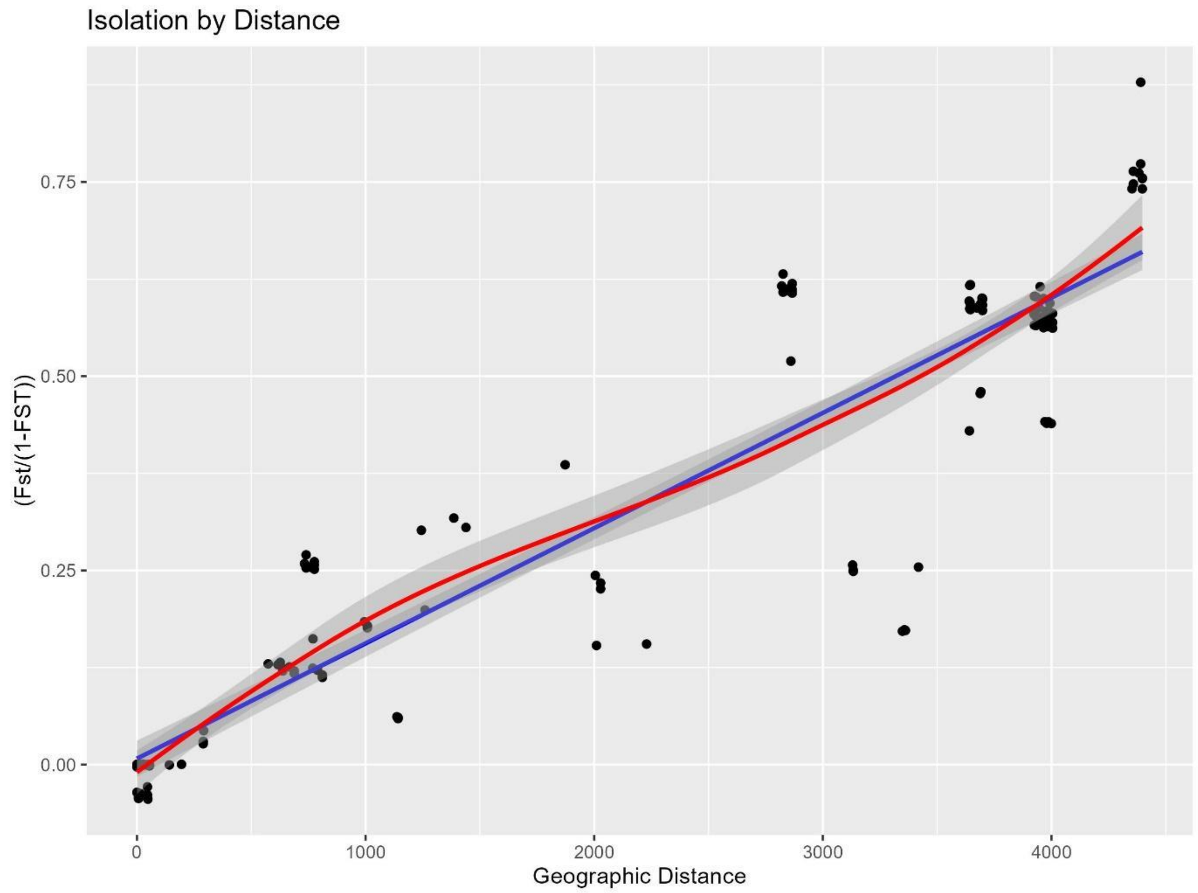


Figure 23: Relationship between genetic differentiation measured as Linearized Fst and geographic distance in km for ***L. schrolli & L.*** **aff.** ***schrolli.*** Each point represents a pairwise comparison between localities. Blue line is the model of linear regression (Lm) and red line the model of polynomial regression (Loess). Grey band represents of standart deviations (se) of the model at 95%.

### Calibration of «*De novo*» Assembly

De novo assembly of raw reads was performed independently for each species using Stacks2. For each species, we apply the same calibration protocol as described in Tran Lu Y et al. (2022), where the assembly and calibration parameter protocol are described in the supplementary information. Briefly, this protocol used the guidelines proposed by Paris et al. (2017) and Mastretta Yanes et al. (2015). These guidelines evaluate various statistics such as the number of assembled loci, the number of polymorphic loci, the number of SNPs, the total number of assembled sites, and the nucleotide diversity to monitor assembly. In addition, genotyping error was investigated using DNA sequencing replicates (the same DNA sample sequenced twice or more, independently). To perform this last step, genotyping error was calculated by measuring the genotyping differences between replicates of the same individual.

*Alviniconcha kojimai*

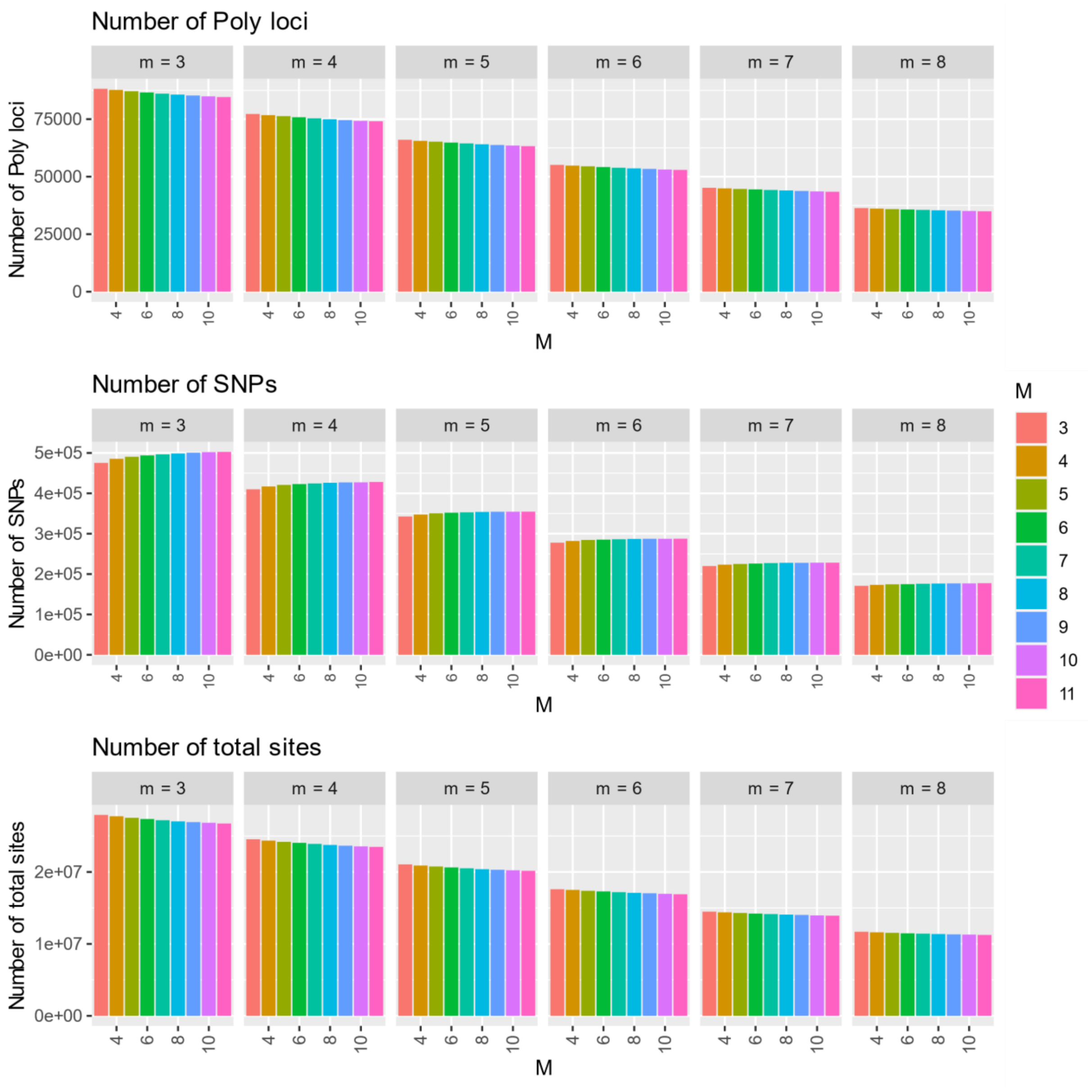


*Figures 1 calibration : Calibration statistics for m (each small box represents a value of m) and M (n=M), representing the number of Polymorphic loci (radtag), number of variants (SNPs) and the total number of sites assembled (variant and non-variant). m is the Minimum stack depth, M the Distance allowed between stacks and n the Distance allowed between catalog loci.*

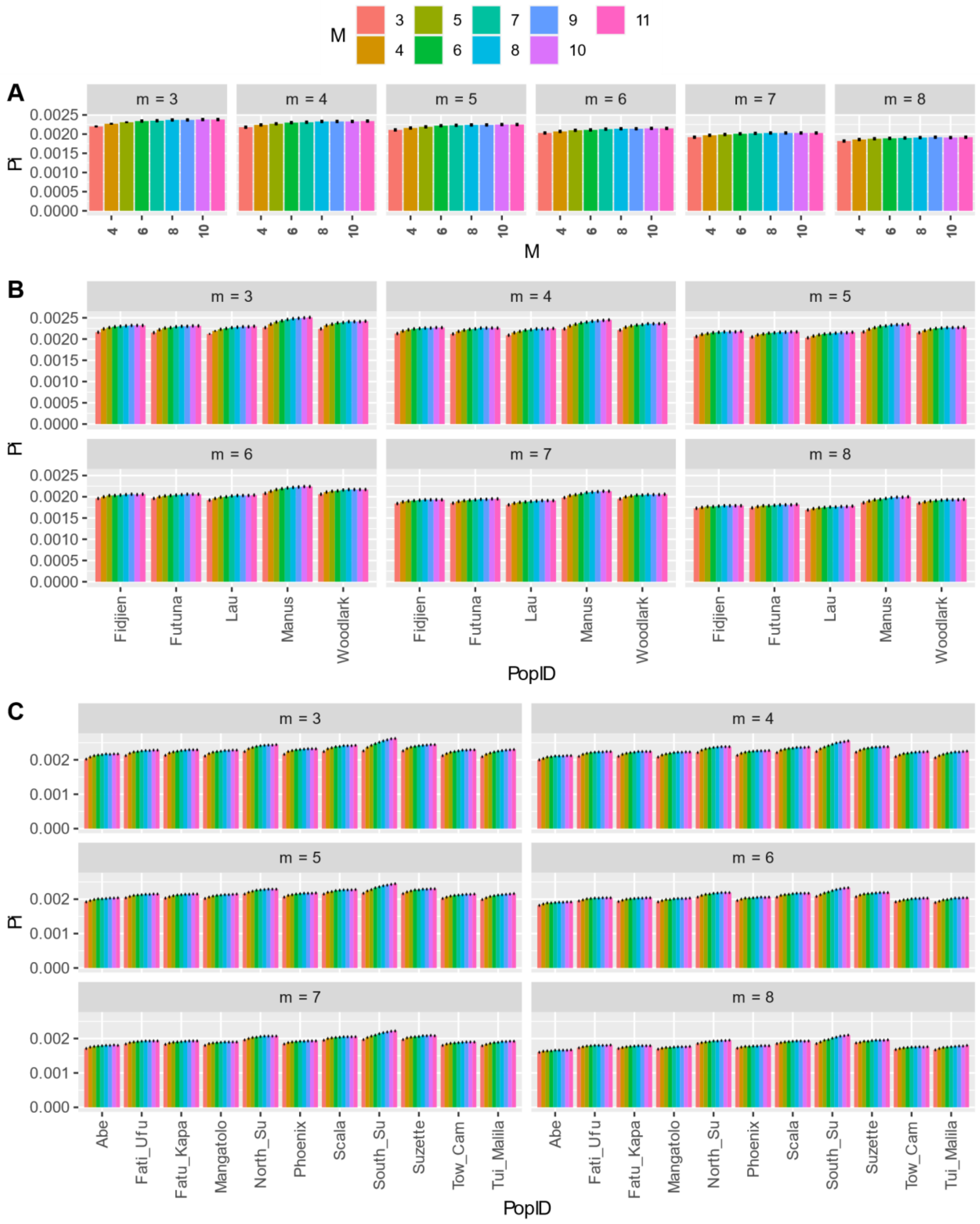


*Figures 2 calibration : Nucleotide diversity (pi) for m (each small box represents a value of m) and M (n=M), estimated with Stacks V2.52 and considering all samples as one population (A), per back-arc-basin (B) and per locality (C). Estimation was performed without replicate. m is the Minimum stack depth, M the Distance allowed between stacks and n the Distance allowed between catalog loci.*

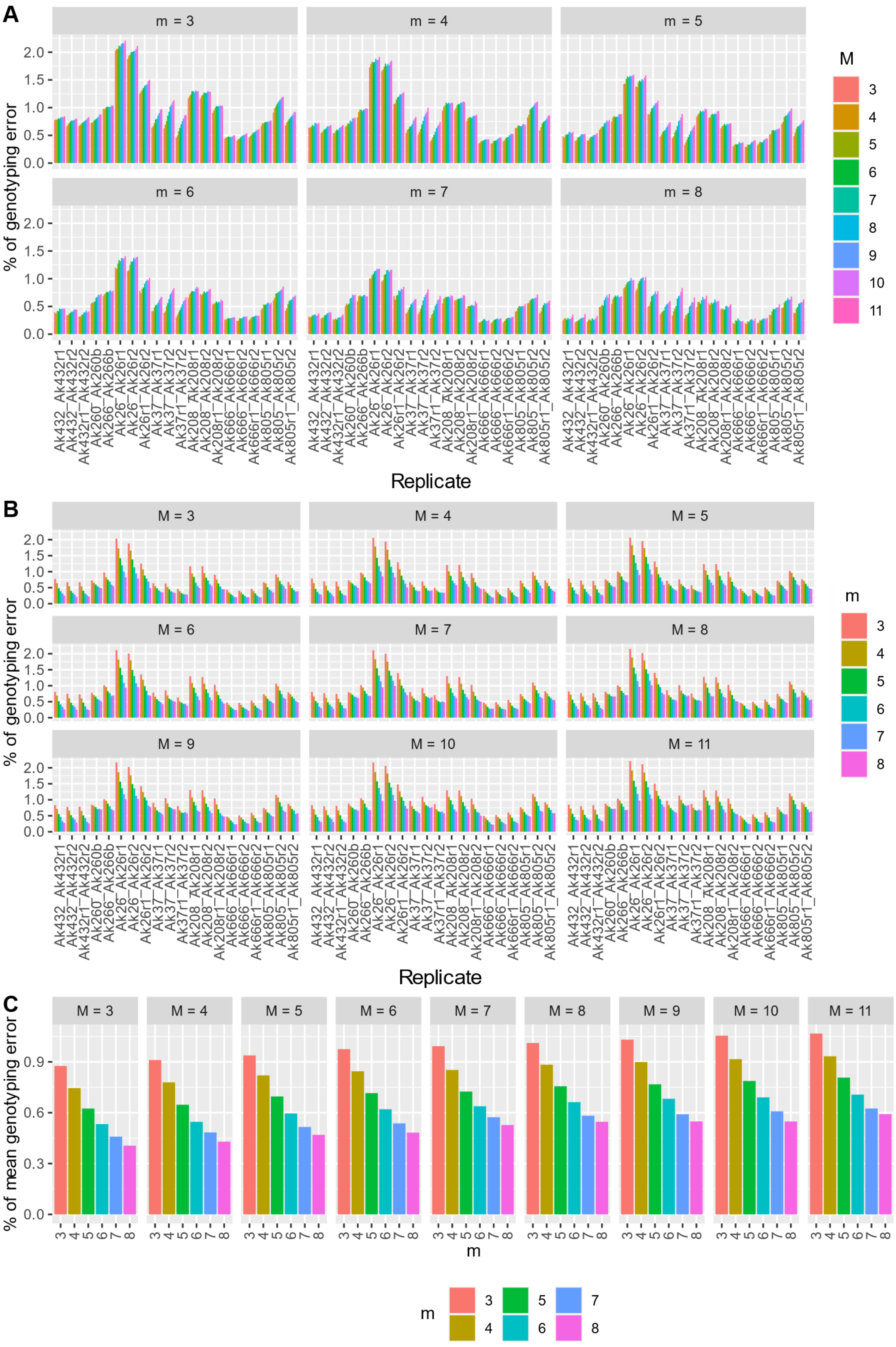
A
m = 3
m = 4
m = 5
m = 6
m = 7
m = 8
% of genotyping error
Replicate
M
B
M = 3
M = 4
M = 5
M = 6
M = 7
M = 8
M = 9
M = 10
M = 11
% of genotyping error
Replicate
m
C
% of mean genotyping error
m
Ak432_Ak432r1
Ak432_Ak432r2
Ak432r1_Ak432r2
Ak260_Ak260b
Ak266_Ak266b
Ak26_Ak26r1
Ak26_Ak26r2
Ak26r1_Ak26r2
Ak37_Ak37r1
Ak37_Ak37r2
Ak37r1_Ak37r2
Ak208_Ak208r1
Ak208_Ak208r2
Ak208r1_Ak208r2
Ak666_Ak666r1
Ak666_Ak666r2
Ak666r1_Ak666r2
Ak805_Ak805r1
Ak805_Ak805r2
Ak805r1_Ak805r2

*Figures 3 calibration : Percent of genotyping error between pairs of replicates for each parameter m and M (n=M). (A) plot of M in function of m. (B) m in function of M. (C) mean genotyping error over all pairs of replicate for each value of M and m. m is the Minimum stack depth, M the Distance allowed between stacks and n the Distance allowed between catalog loci.*

*Shinkailepas tollmanni*

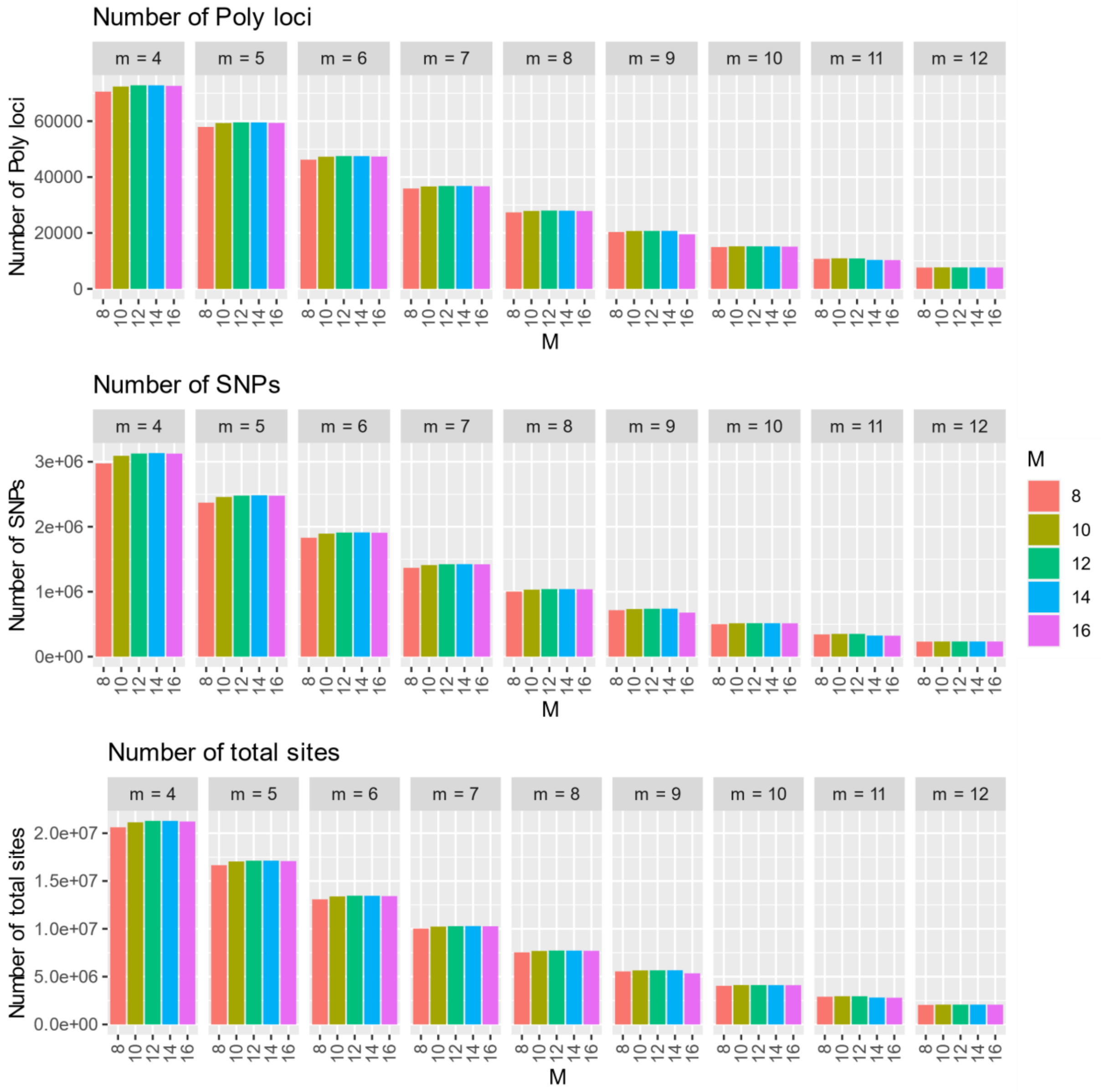


*Figures 4 calibration : Calibration statistics for m (each small box represents a value of m) and M (n=M), representing the number of Polymorphic loci (radtag), number of variants (SNPs) and the total number of sites assembled (variant and non-variant). m is the Minimum stack depth, M the Distance allowed between stacks and n the Distance allowed between catalog loci.*

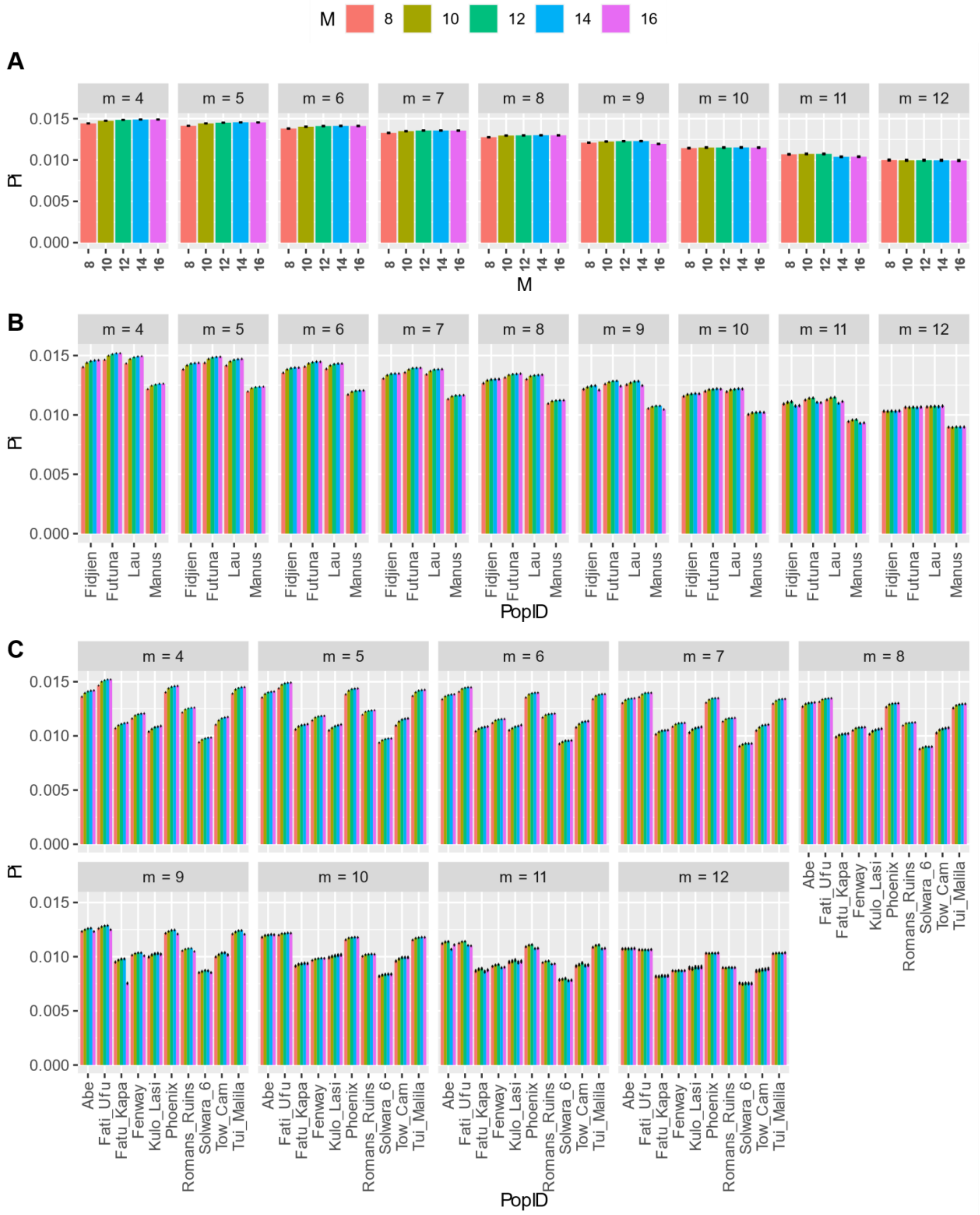


*Figures 5 calibration : Nucleotide diversity (pi) for m (each small box represents a value of m) and M (n=M), estimated with Stacks V2.52 and considering all samples as one population (A), per back-arc-basin (B) and per locality (C). Estimation was performed without replicate. m is the Minimum stack depth, M the Distance allowed between stacks and n the Distance allowed between catalog loci.*

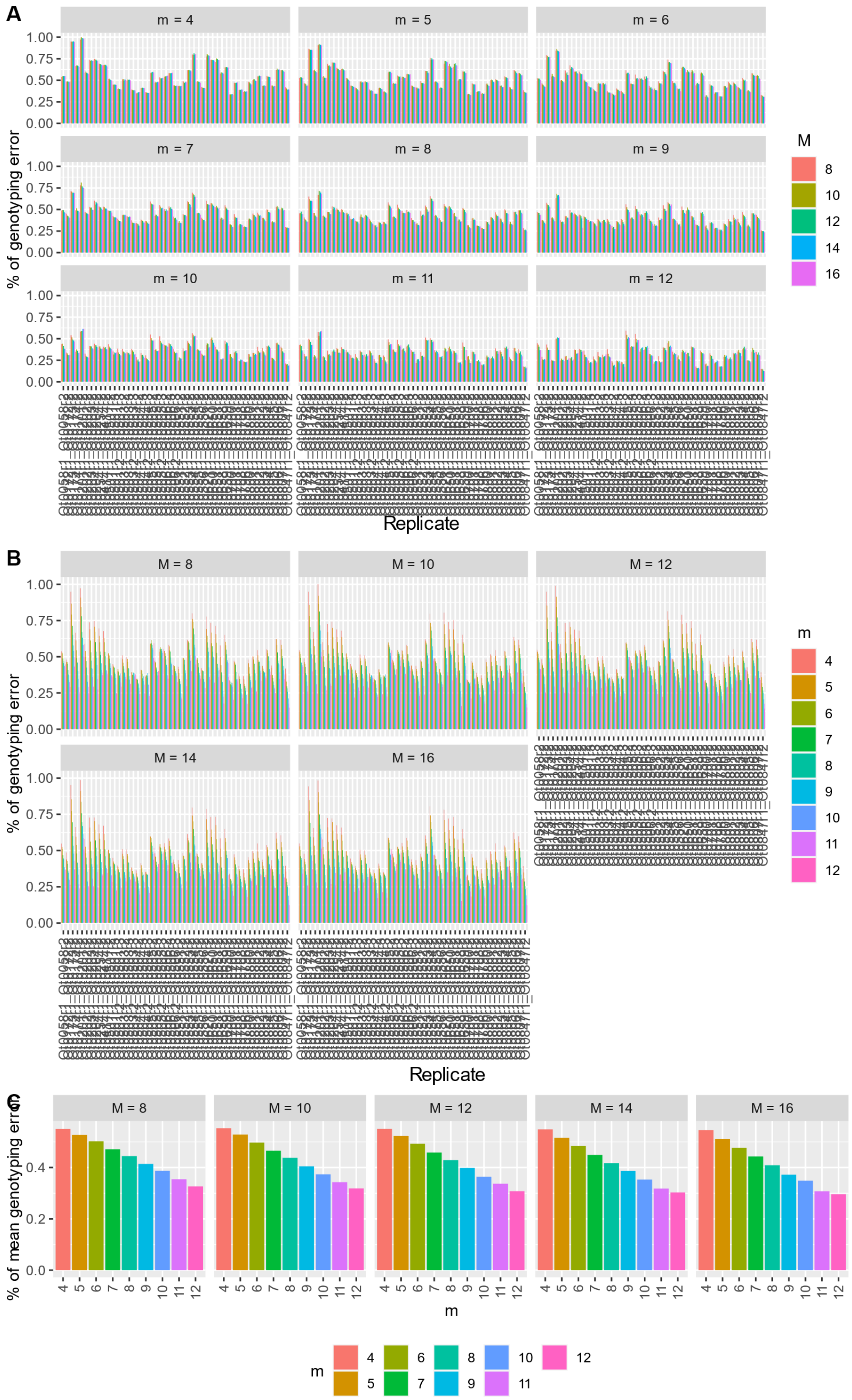
A
m = 4
m = 5
m = 6
m = 7
m = 8
m = 9
m = 10
m = 11
m = 12
% of genotyping error
Replicate
M
8
10
12
14
16
B
M = 8
M = 10
M = 12
M = 14
M = 16
% of genotyping error
Replicate
m
4
5
6
7
8
9
10
11
12
C
% of mean genotyping error
m

*Figures 6 calibration : Percent of genotyping error between pairs of replicates for each parameter m and M (n=M). (A) plot of M in function of m. (B) m in function of M. (C) mean genotyping error over all pairs of replicate for each value of M and m. m is the Minimum stack depth, M the Distance allowed between stacks and n the Distance allowed between catalog loci.*

### *Eochionelasmus ohtai*

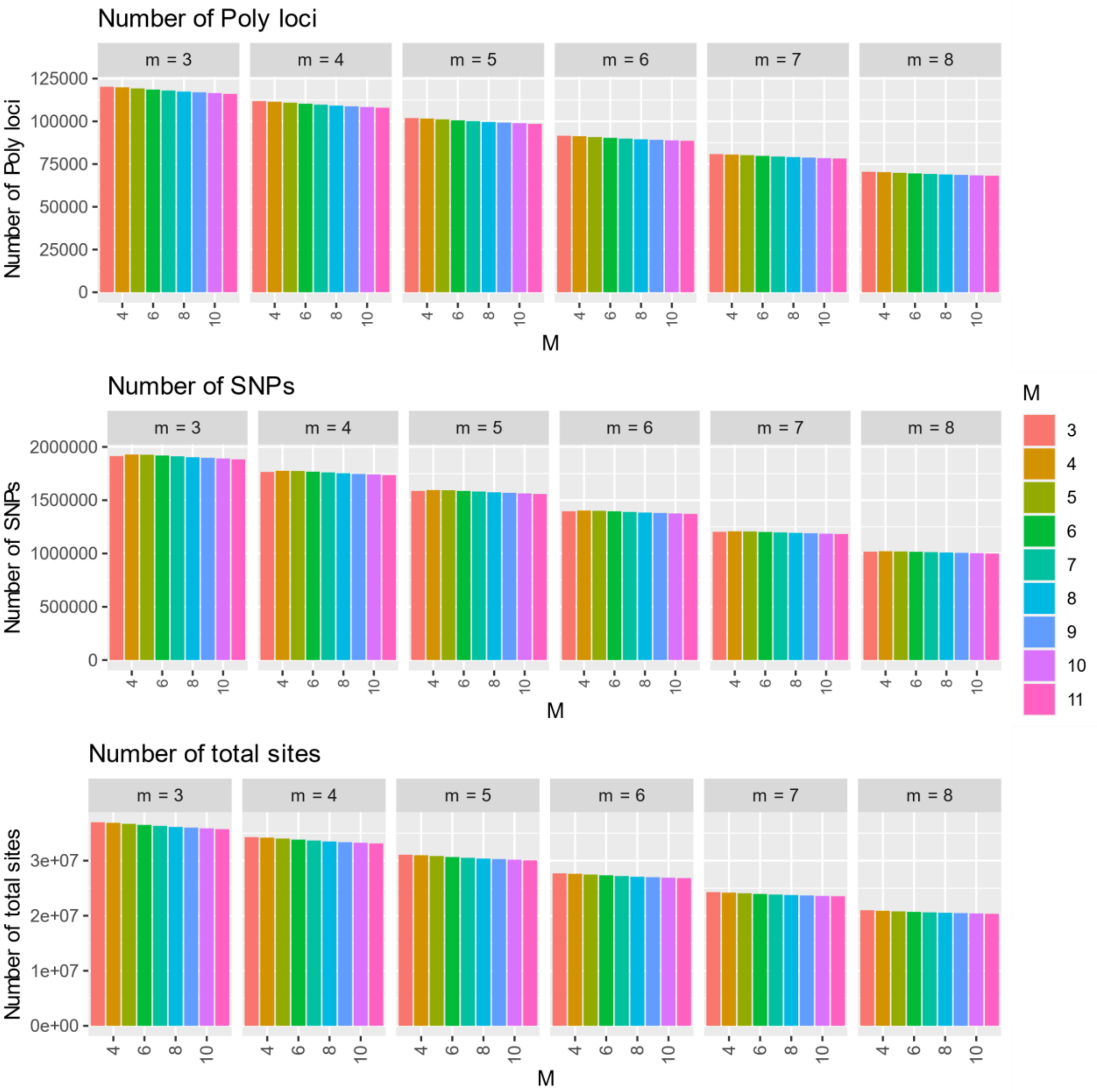


*Figures 7 calibration : Calibration statistics for m (each small box represents a value of m) and M (n=M), representing the number of Polymorphic loci (radtag), number of variants (SNPs) and the total number of sites assembled (variant and non-variant). m is the Minimum stack depth, M the Distance allowed between stacks and n the Distance allowed between catalog loci.*

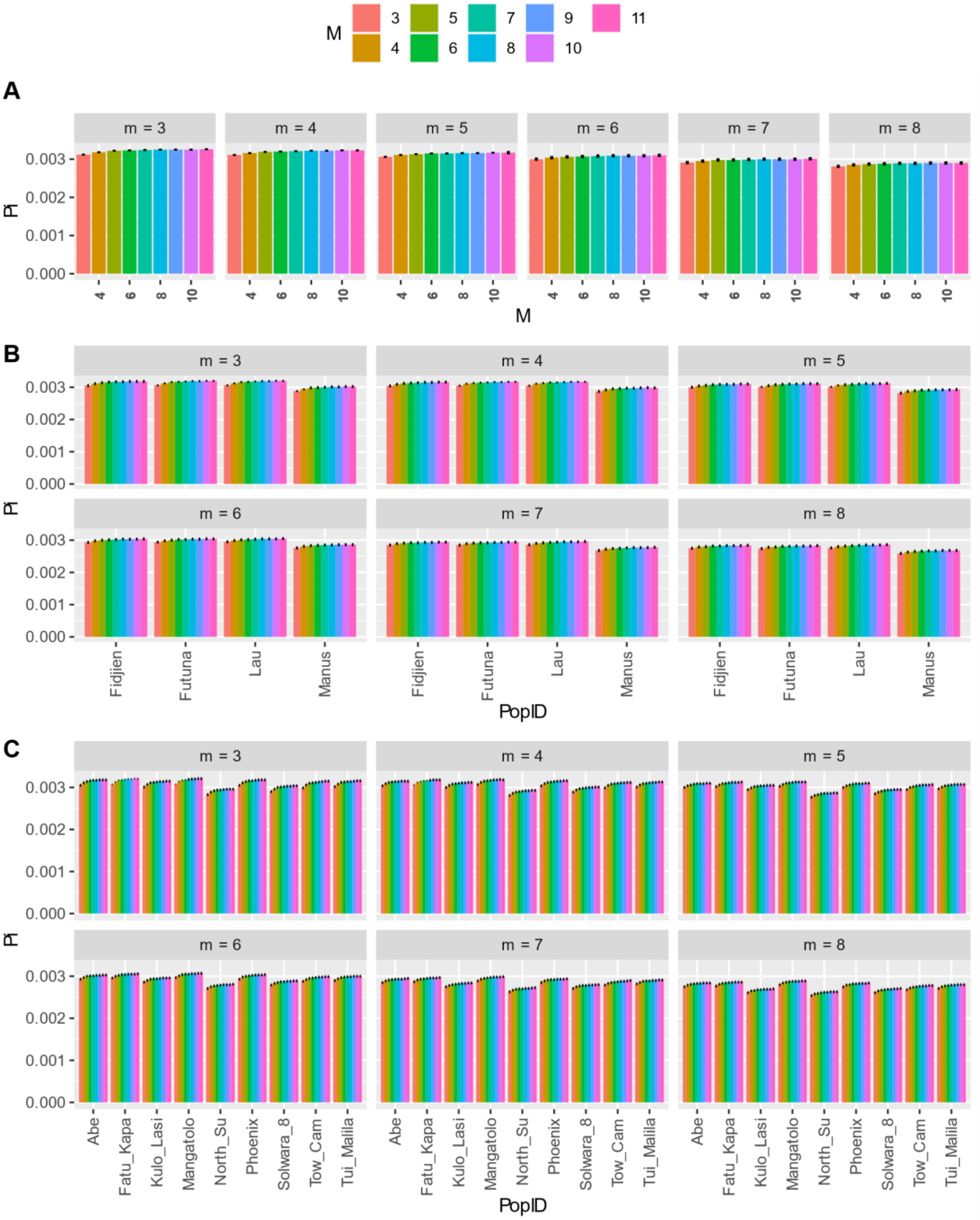


*Figures 8 calibration : Nucleotide diversity (pi) for m (each small box represents a value of m) and M (n=M), estimated with Stacks V2.52 and considering all samples as one population (A), per back-arc-basin (B) and per locality (C). Estimation was performed without replicate. m is the Minimum stack depth, M the Distance allowed between stacks and n the Distance allowed between catalog loci.*

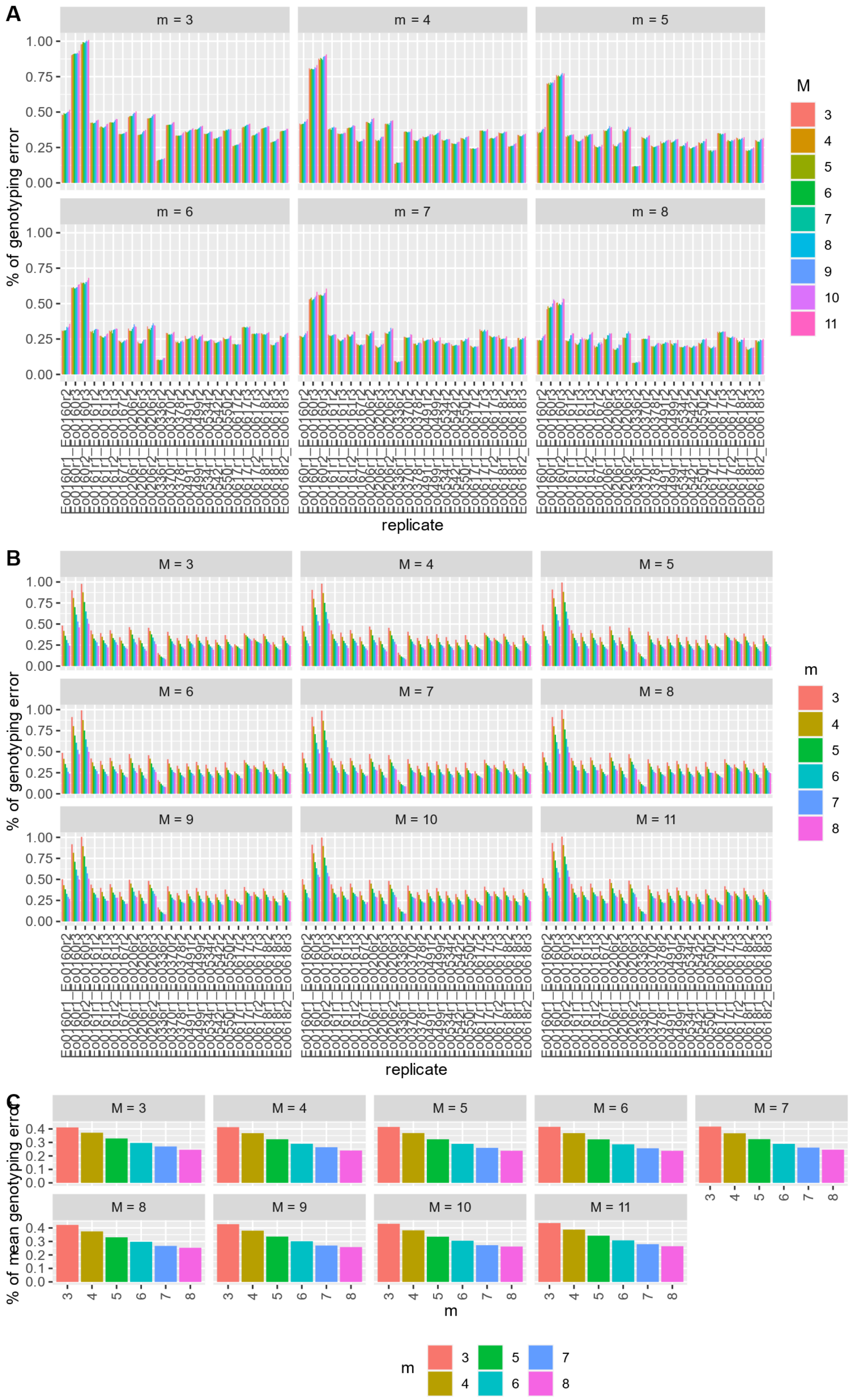
A
m = 3
m = 4
m = 5
m = 6
m = 7
m = 8
% of genotyping error
1.00
0.75
0.50
0.25
0.00
replicate
M
3
4
5
6
7
8
9
10
11
B
M = 3
M = 4
M = 5
M = 6
M = 7
M = 8
M = 9
M = 10
M = 11
% of genotyping error
replicate
m
C
% of mean genotyping error
0.4
0.3
0.2
0.1
0.0
m
3
4
5
6
7
8

*Figures 9 calibration : Percent of genotyping error between pairs of replicates for each parameter m and M (n=M). (A). Plot of M in function of m. (B) m in function of M. (C) mean genotyping error over all pairs of replicate for each value of M and m. m is the Minimum stack depth, M the Distance allowed between stacks and n the Distance allowed between catalog loci.*

#### *Bathymodiolus manusensis*

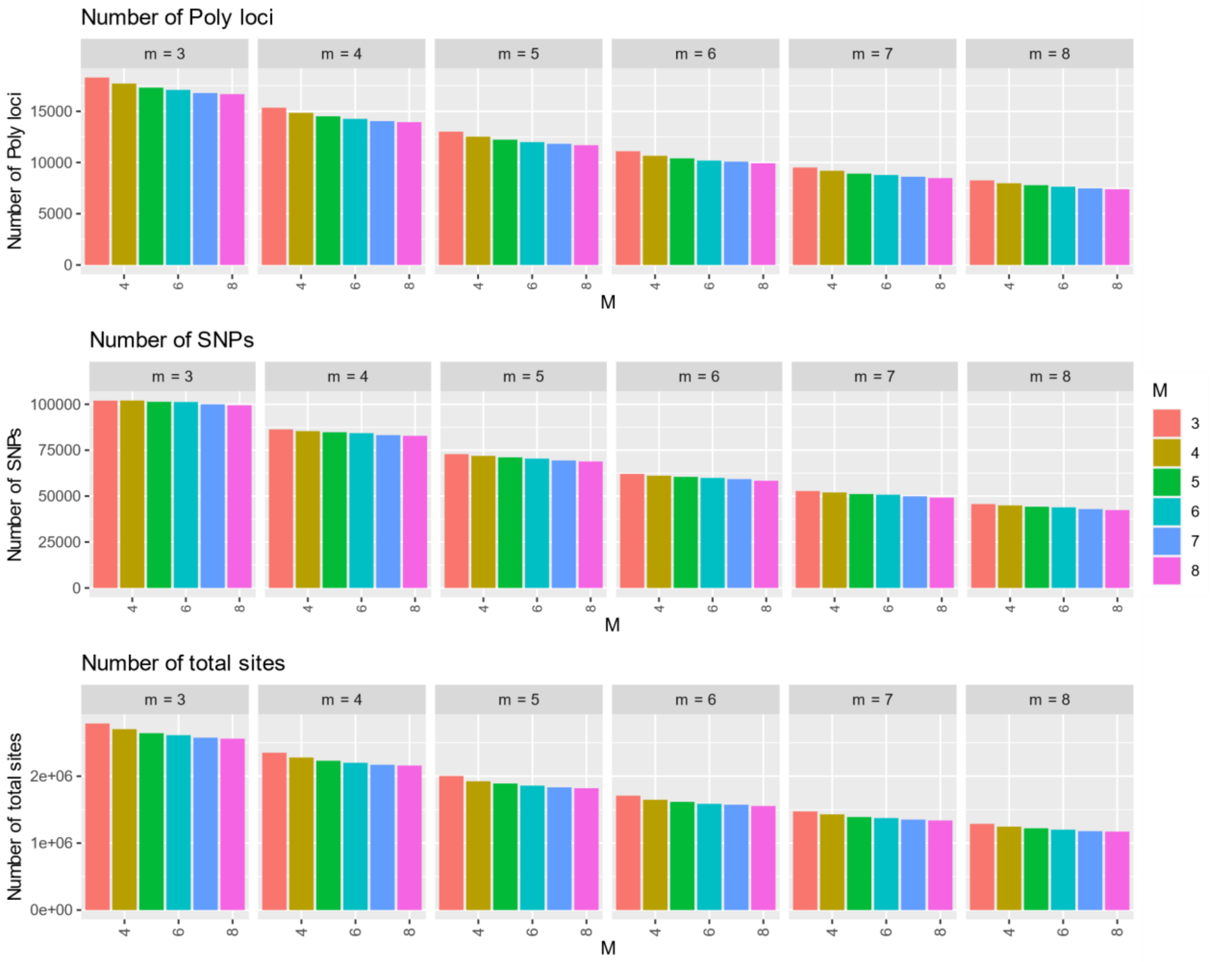


*Figures 10 calibration : Calibration statistics for m (each small box represents a value of m) and M (n=M), representing the number of Polymorphic loci (radtag), number of variants (SNPs) and the total number of sites assembled (variant and non-variant). m is the Minimum stack depth, M the Distance allowed between stacks and n the Distance allowed between catalog loci.*

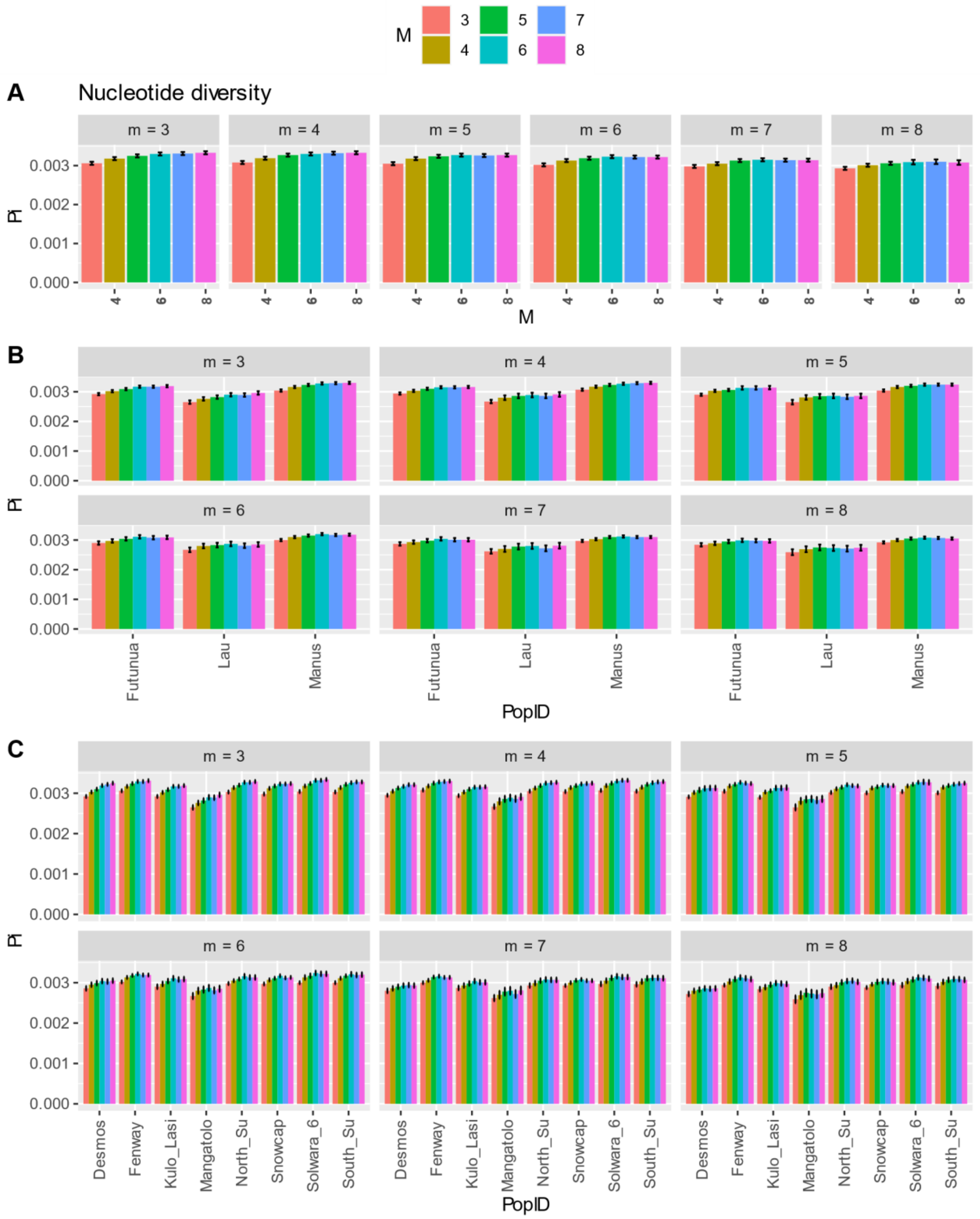


*Figures 11 calibration : Nucleotide diversity (pi) for m (each small box represents a value of m) and M (n=M), estimated with Stacks V2.52 and considering all samples as one population (A), per back-arc-basin (B) and per locality (C). estimation was performed without replicate. m is the Minimum stack depth, M the Distance allowed between stacks and n the Distance allowed between catalog loci.*

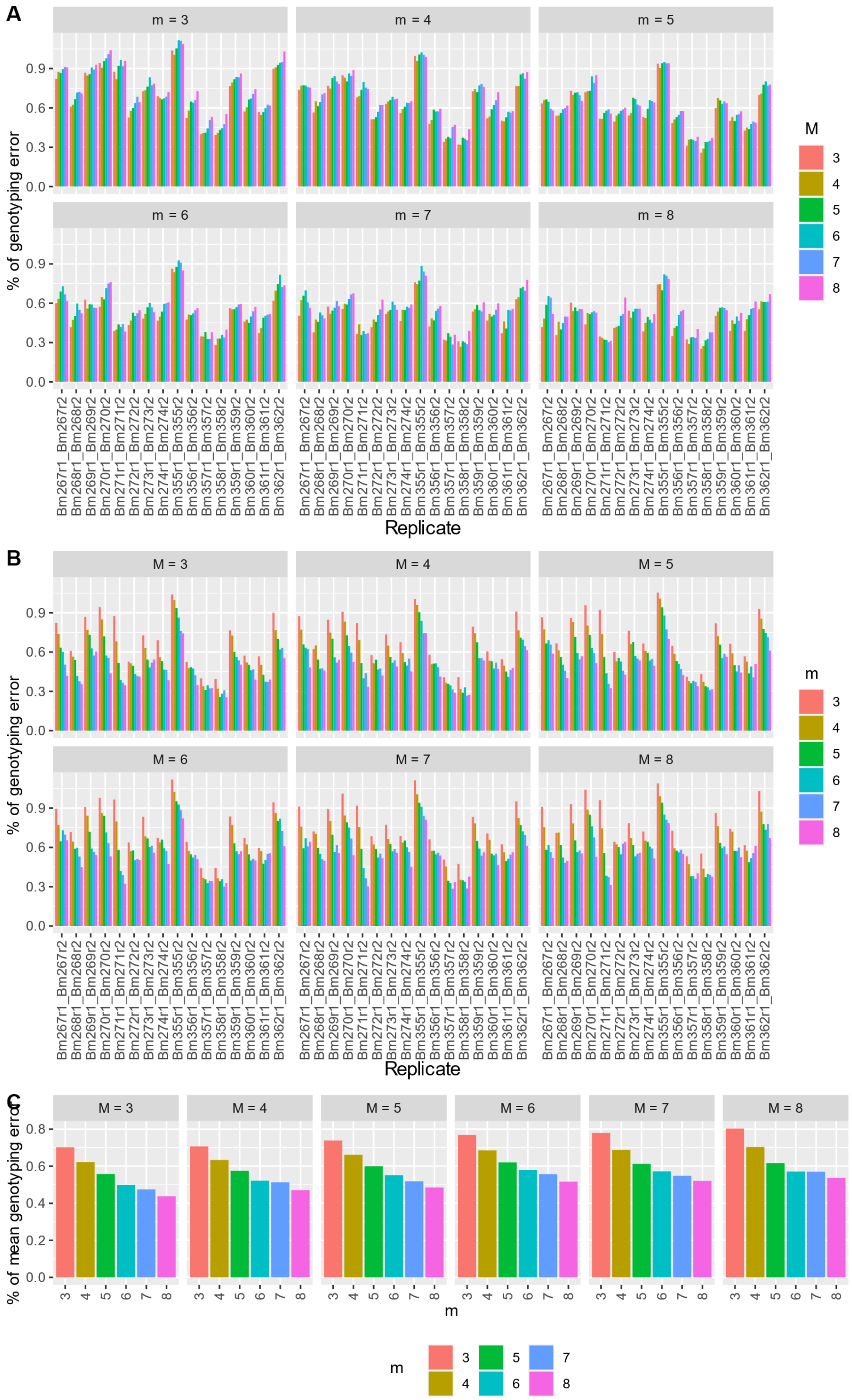
A
m = 3
m = 4
m = 5
m = 6
m = 7
m = 8
% of genotyping error
Replicate
M
B
M = 3
M = 4
M = 5
M = 6
M = 7
M = 8
% of genotyping error
Replicate
m
C
% of mean genotyping error
m
Bm267r1_Bm267r2
Bm268r1_Bm268r2
Bm269r1_Bm269r2
Bm270r1_Bm270r2
Bm271r1_Bm271r2
Bm272r1_Bm272r2
Bm273r1_Bm273r2
Bm274r1_Bm274r2
Bm355r1_Bm355r2
Bm356r1_Bm356r2
Bm357r1_Bm357r2
Bm358r1_Bm358r2
Bm359r1_Bm359r2
Bm360r1_Bm360r2
Bm361r1_Bm361r2
Bm362r1_Bm362r2

*Figures 12 calibration : Percent of genotyping error between pairs of replicates for each parameter m and M (n=M). (A) plot of M in function of m.  (B) m in function of M. (C) mean genotyping error over all pairs of replicate for each value of M and m.  m is the Minimum stack depth, M the Distance allowed between stacks and n the Distance allowed between catalog loci.*

#### *Branchinotogluma segonzaci*

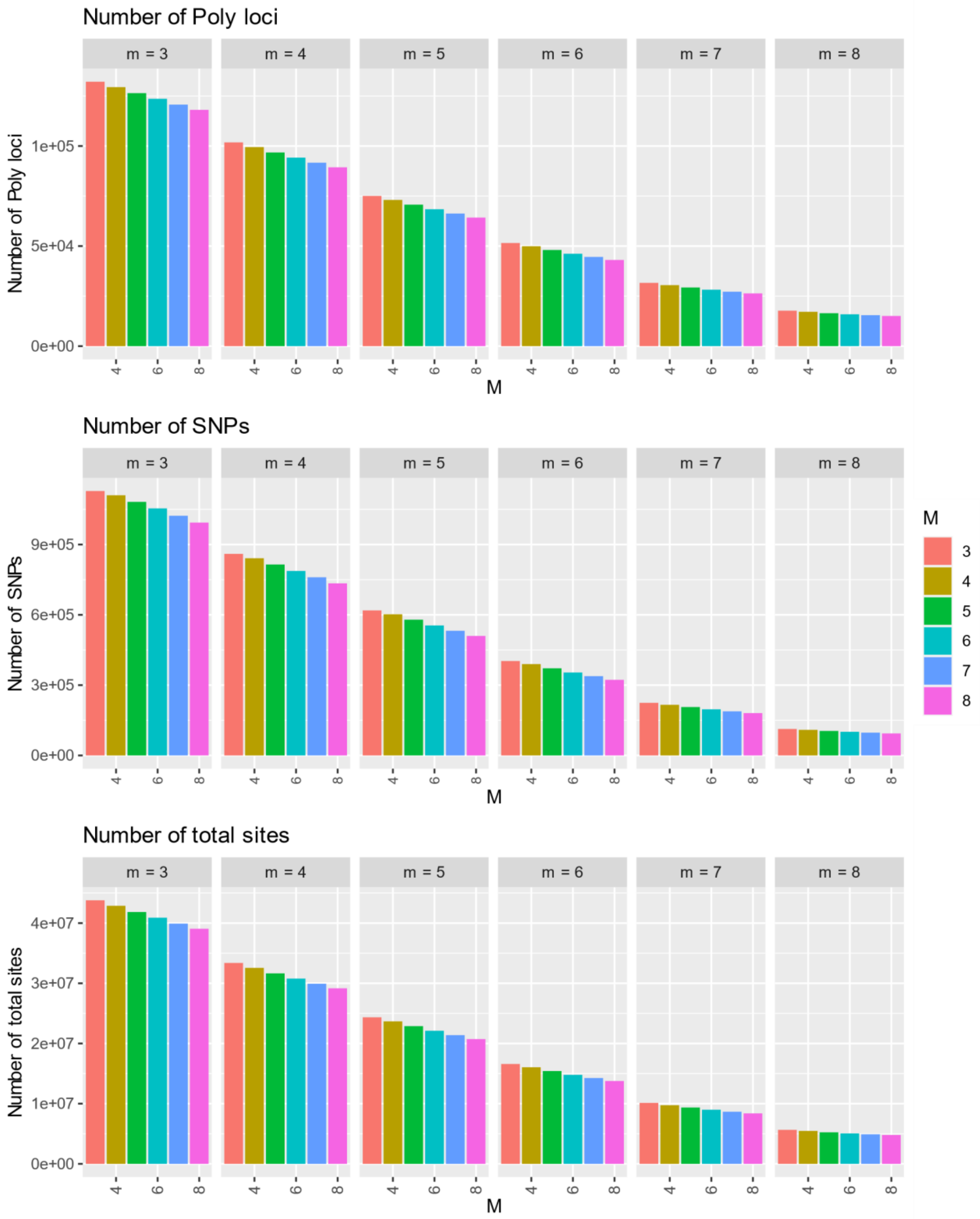


*Figures 13 calibration : Calibration statistics for m (each small box represents a value of m) and M (n=M), representing the number of Polymorphic loci (radtag), number of variants (SNPs) and the total number of sites assembled (variant and non-variant). m is the Minimum stack depth, M the Distance allowed between stacks and n the Distance allowed between catalog loci.*

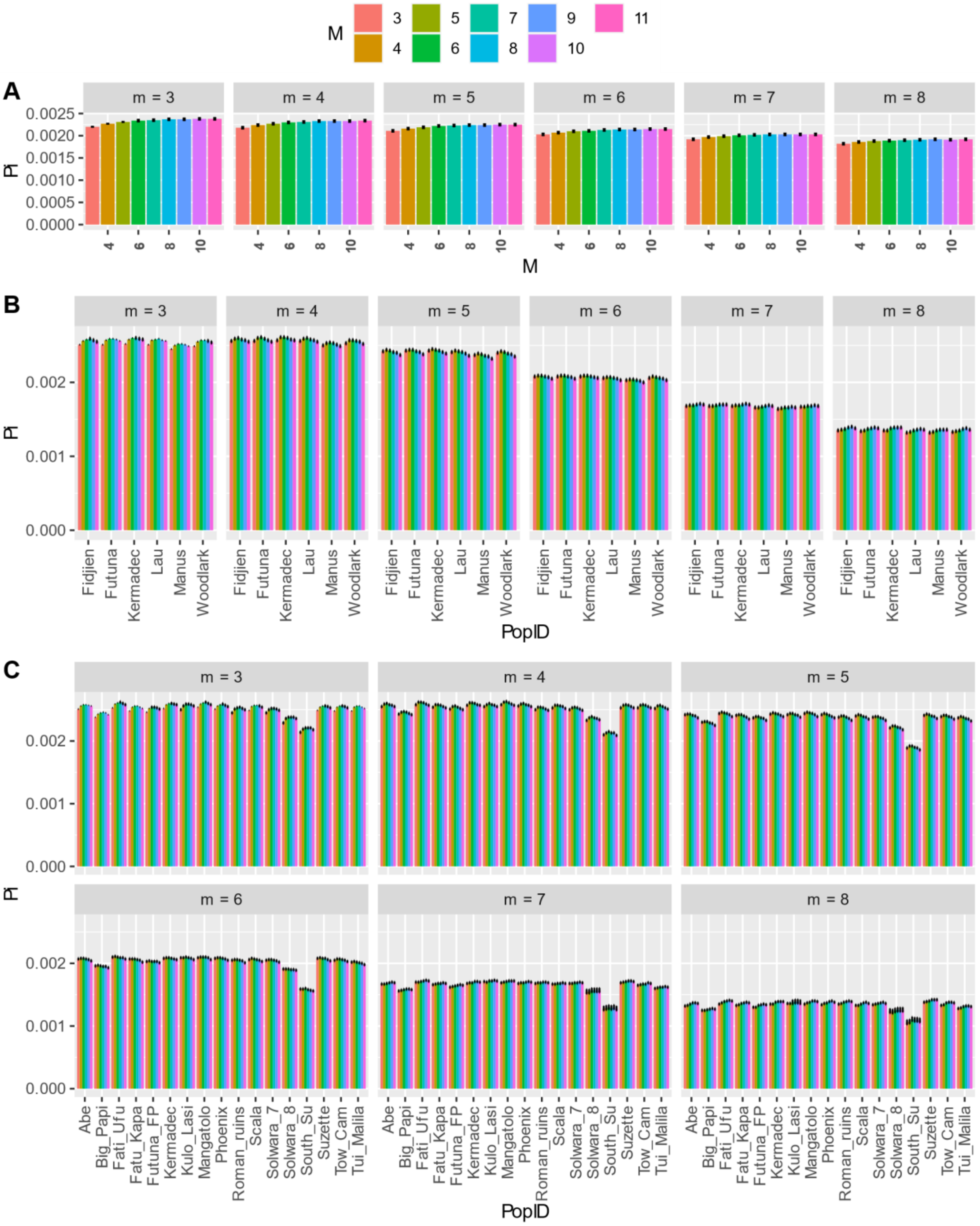


*Figures 14 calibration : Nucleotide diversity (pi) for m (each small box represents a value of m) and M (n=M), estimated with Stacks V2.52 and considering all samples as one population (A), per back-arc-basin (B) and per locality (C). estimation was performed without replicate. m is the Minimum stack depth, M the Distance allowed between stacks and n the Distance allowed between catalog loci.*

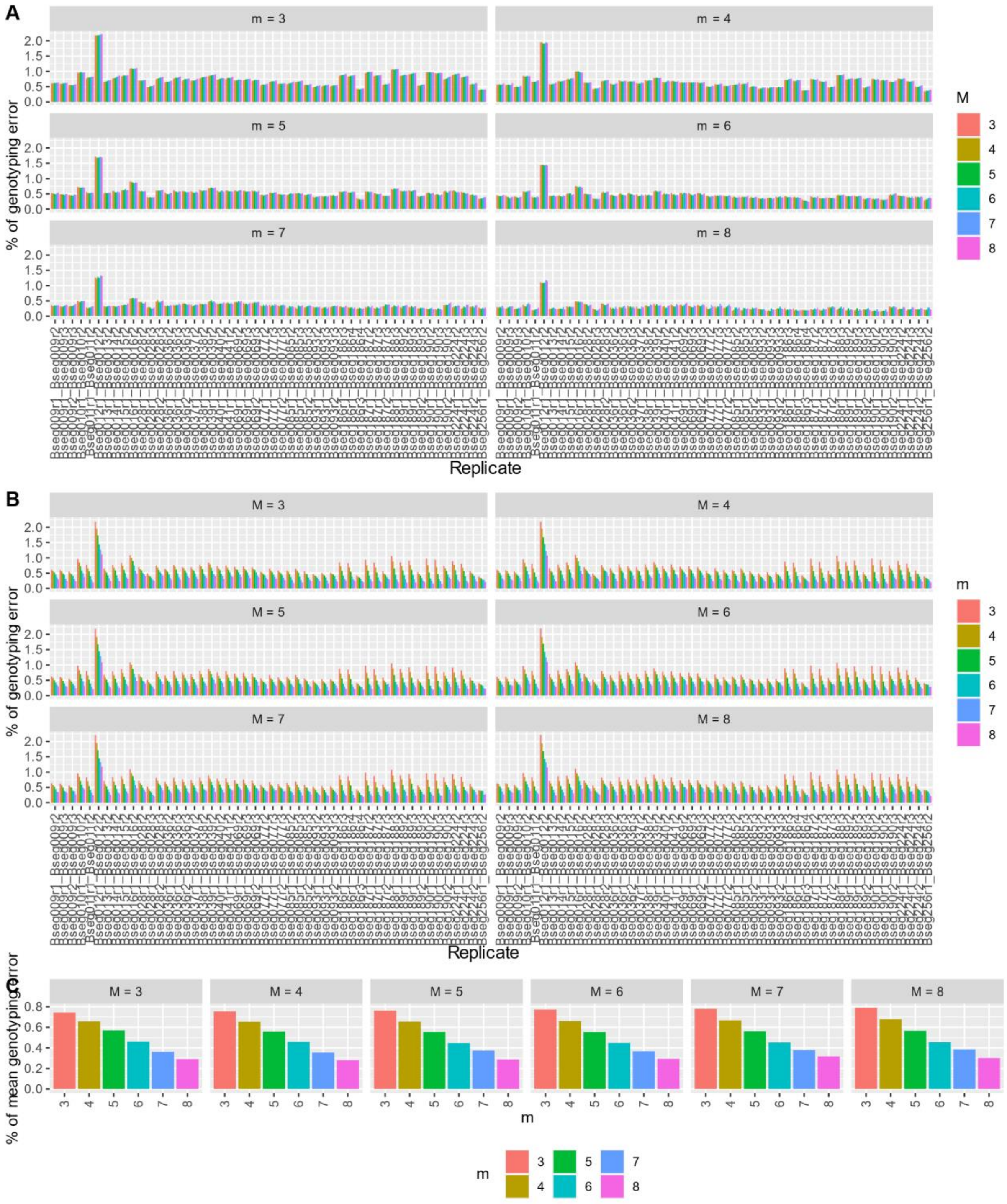


*Figures 15 calibration : Percent of genotyping error between pairs of replicates for each parameter m and M (n=M). (A) plot of M in function of m. (B) m in function of M. (C) mean genotyping error over all pairs of replicate for each value of M and m. m is the Minimum stack depth, M the Distance allowed between stacks and n the Distance allowed between catalog loci.*

*Lepetodrillus schrolli* & *L.* aff. *schrolli*

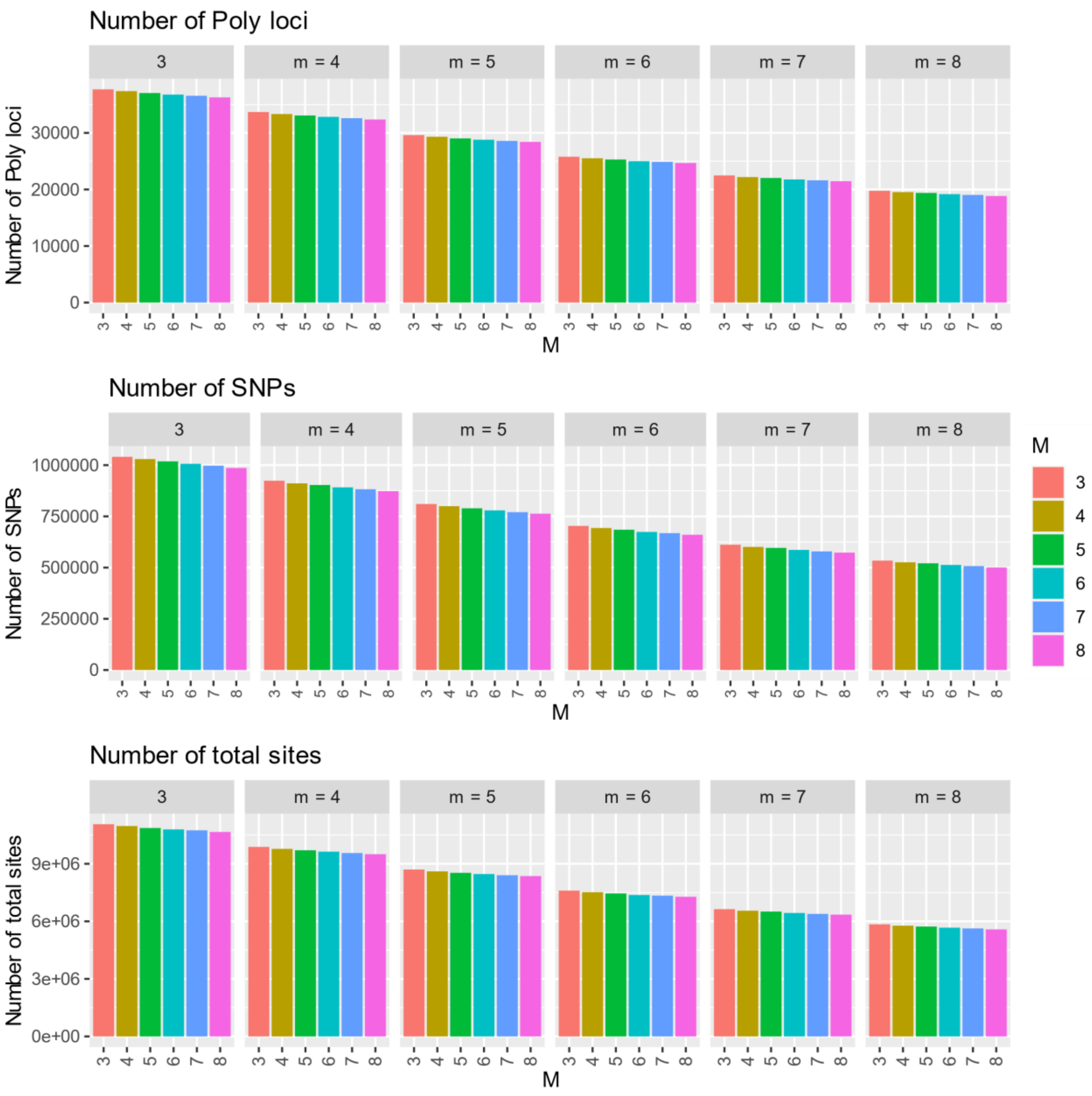


*Figures 16 calibration : Calibration statistics for m (each small box represents a value of m) and M (n=M), representing the number of Polymorphic loci (radtag), number of variants (SNPs) and the total number of sites assembled (variant and non-variant). m is the Minimum stack depth, M the Distance allowed between stacks and n the Distance allowed between catalog loci.*

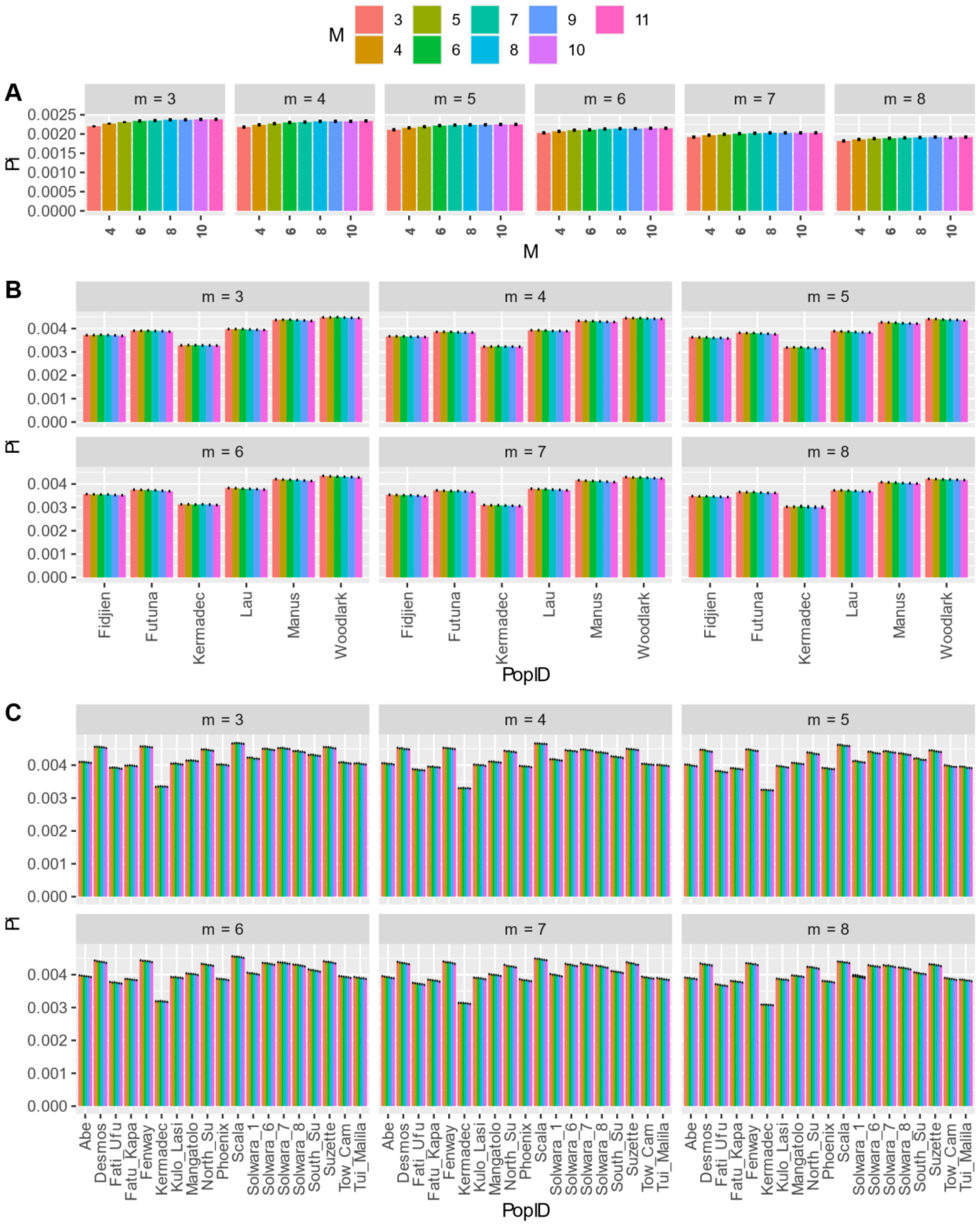


*Figures 17 calibration : Nucleotide diversity (pi) for m (each small box represents a value of m) and M (n=M), estimated with Stacks V2.52 and considering all samples as one population (A), per back-arc-basin (B) and per locality (C). estimation was*

*performed without replicate. m is the Minimum stack depth, M the Distance allowed between stacks and n the Distance allowed between catalog loci.*

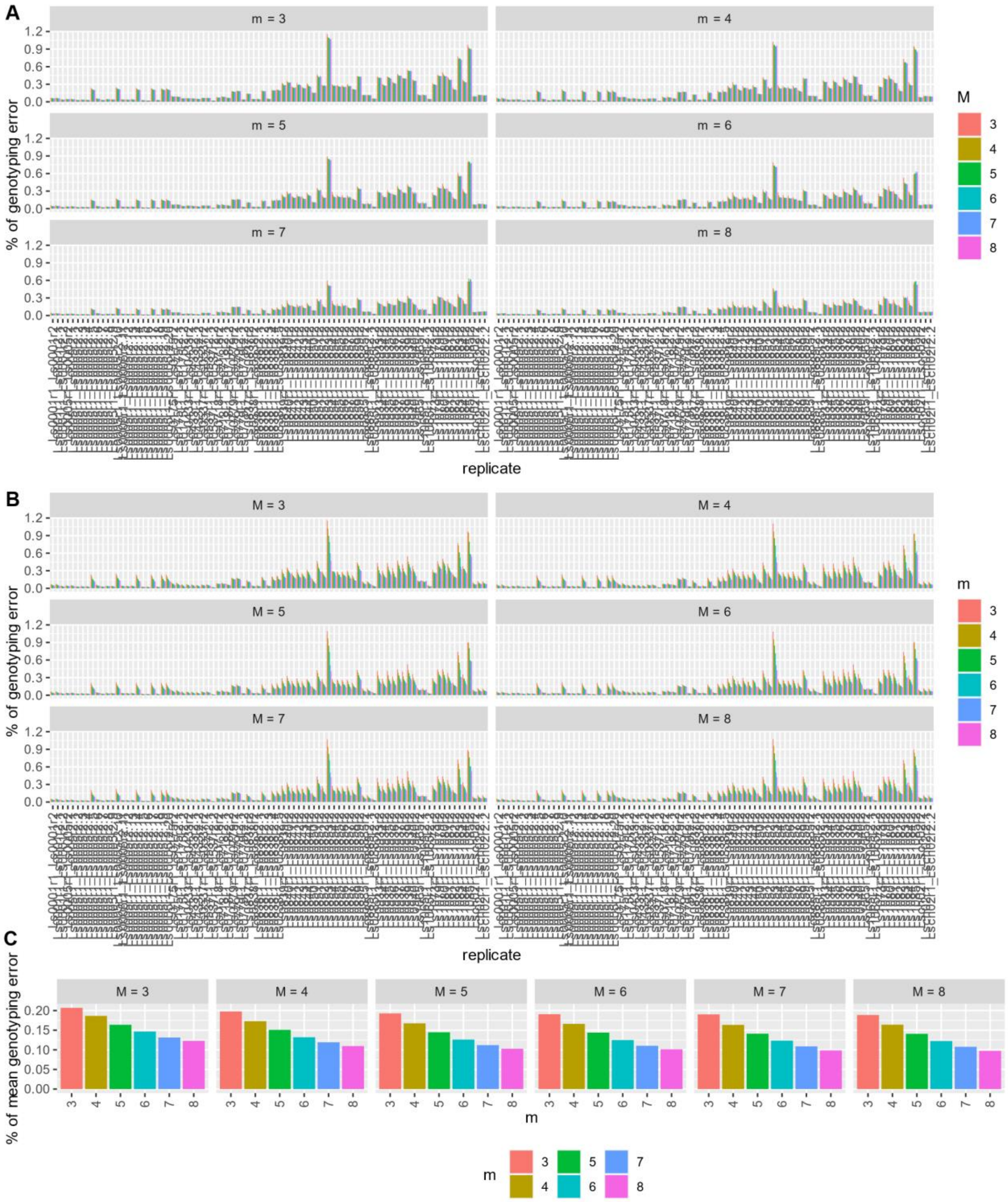

*Figures 18 calibration : Percent of genotyping error between pairs of replicates for each parameter m and M (n=M). (A) plot of M in function of m.  (B) m in function of M. (C) mean genotyping error over all pairs of replicate for each value of M and m.  m is the Minimum stack depth, M the Distance allowed between stacks and n the Distance allowed between catalog loci.*
